# Broad-scale variation in human genetic diversity levels is predicted by purifying selection on coding and non-coding elements

**DOI:** 10.1101/2021.07.02.450762

**Authors:** David Murphy, Eyal Elyashiv, Guy Amster, Guy Sella

## Abstract

Analyses of genetic variation in many taxa have established that neutral genetic diversity is shaped by natural selection at linked sites. Whether the mode of selection is primarily the fixation of strongly beneficial alleles (selective sweeps) or purifying selection on deleterious mutations (background selection) remains unknown, however. We address this question in humans by fitting a model of the joint effects of selective sweeps and background selection to autosomal polymorphism data from the 1000 Genomes Project. After controlling for variation in mutation rates along the genome, a model of background selection alone explains ∼60% of the variance in diversity levels at the megabase scale. Adding the effects of selective sweeps driven by adaptive substitutions to the model does not improve the fit, and when both modes of selection are considered jointly, selective sweeps are estimated to have had little or no effect on linked neutral diversity. The regions under purifying selection are best predicted by phylogenetic conservation, with ∼80% of the deleterious mutations affecting neutral diversity occurring in non-exonic regions. Thus, background selection is the dominant mode of linked selection in humans, with marked effects on diversity levels throughout autosomes.

## Introduction

Selection at a given locus in the genome affects diversity levels at sites linked to it (*1-9*). In particular, when a new, strongly beneficial mutation increases in frequency to fixation in the population, it carries with it the haplotype on which it arose, thus reducing levels of neutral diversity nearby, in what is sometimes called a ‘hard selective sweep’ (*2, 3*). ‘Soft sweeps’, particularly those in which an allele segregates at low frequency before becoming beneficial and sweeping to fixation, and ‘partial sweeps’, in which a beneficial mutation rapidly increases to intermediate frequencies, also reduce neutral diversity levels near the selected sites (*10-15*). Similarly, when deleterious mutations are eliminated from the population by selection, so are the haplotypes on which they lie. This process too reduces diversity levels near selected sites, in a phenomenon known as ‘background selection’ (*5-7, 16-18*). Because the lengths of the haplotypes associated with selected alleles depend on the recombination rate, selection causes a greater reduction in levels of linked neutral genetic diversity in regions with lower rates of recombination or a greater density of selected sites. These predicted relationships have been observed in numerous taxa, including plants, Drosophila, rodents and primates, establishing the effects of linked selection in a wide range of species (*4, 9, 19-28*).

More recently, the advent of large genomic datasets and detailed functional annotations have made it possible to infer the effects of linked selection and build maps that predict levels of diversity along the genome (*29, 30*) (also see (*6, 7, 31*)). The first effort predated the availability of genome-wide resequencing data, relying instead on information about incomplete lineage sorting among human, chimpanzee and gorilla, which reflects variation in diversity levels along the genome in the common ancestor of humans and chimpanzees (*29*). This pioneering paper showed that a model of background selection fits variation in human-chimpanzee divergence levels along the genome remarkably well, with only a few parameters. Nonetheless, the estimate of the deleterious mutation rate was unrealistically high, much greater than estimates of the total mutation rate per site in humans (*32, 33*) (SOM Section 5), raising the possibility that it was soaking up effects of other modes of selection, notably those of selective sweeps. Subsequent work suggested that selective sweeps have had little effect on diversity levels in humans, with no more of a reduction of diversity around nonsynonymous substitutions than around synonymous ones (*34, 35*), but the interpretation of these findings was contested. Notably, it was suggested that the effects of background selection are more pronounced around synonymous substitutions than nonsynonymous substitutions, thereby masking the effects of selective sweeps (*36*). Thus, we still lack an understanding of the relative contribution of adaptation and purifying selection (*37*), as well as a map of their effects on human diversity levels.

## Results

### Model and inference

Here, we resolve these issues by considering the effects of background selection and selective sweeps on diversity levels jointly (Fig. 1 and SOM Section 1). To this end, we model the effects of background selection on neutral diversity levels as a function of genetic distance from regions that may be under purifying selection (following (*6*) and (*38*)), where the deleterious mutation rate per site and distribution of selection effects in a given type of region (e.g., exons) are parameters to be estimated (Fig. 1A). In turn, the effects of sweeps are modeled as a function of genetic distance from substitutions on the human lineage that may have been beneficial (following (*39*) and (*40*)), where the fraction of substitutions of a given type (e.g., nonsynonymous) that were beneficial and their distribution of selection effects are again parameters to be estimated (Fig. 1B). Importantly, our model should capture the effects of any kind of sweeps, be they hard, partial or soft, so long as they eventually resulted in a substitution and affected diversity levels nearby (see SOM Section D in (*30*)). Given the positions of different types of putatively selected regions and substitutions, their corresponding selection parameters, and a fine-scale genetic map, the model allows us to calculate the marginal probability that any given neutral site in the genome is polymorphic in a sample (Fig. 1C). Provided measurements of polymorphism at neutral positions throughout the genome, we combine information across sites and samples to calculate the composite likelihood of selection parameters, and find the parameter values that maximize this likelihood (Fig. 1 and SOM Section 1). In addition to parameter estimation, this approach yields a map of the expected neutral diversity levels along the genome (Fig. 1C).

**Fig. 1.**
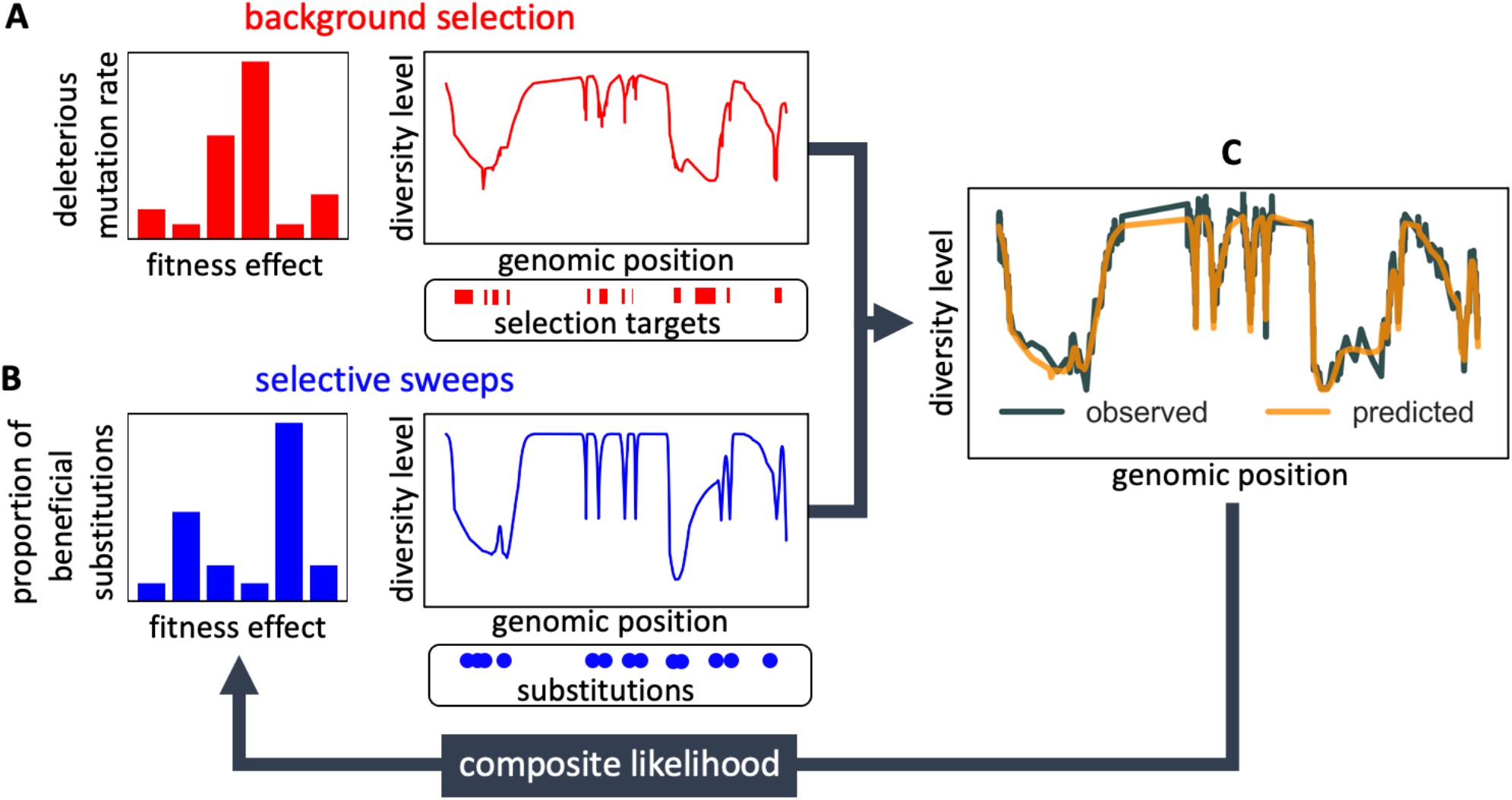
Modeling and inferring the effects of linked selection in humans. Given the putative targets of selection and corresponding selection parameters (**A** and **B**), we calculate the expected neutral diversity levels along the genome (**C**). We infer the selection parameters by maximizing their composite likelihood given observed diversity levels (C). Based on these parameter estimates, we calculate a map of the expected effects of selection on linked diversity levels.

To infer the effects of background selection and selective sweeps on human diversity levels, we analyze autosomal polymorphism data from 26 human populations, collected in Phase III of the 1000 Genomes Project (*41*). Here, we focus on data from 108 genomes sampled from the Yoruba population (YRI), but we get similar results for the other populations (SOM Sections 7 and 9). To estimate diversity levels at neutral sites, we focus on non-genic autosomal sites that are the least conserved in a multiple sequence alignment of 25 supra-primates (see SOM Section 3.1). To control for variation in the mutation rates across neutral sites, we use estimates of the relative mutation rate for contiguous, non-overlapping blocks of 6000 putatively neutral sites, obtained from substitution rates in an eight-primate phylogeny (see SOM Section 3.3). To minimize the confounding of recombination rate estimates and diversity levels, we use a high-resolution genetic map inferred from ancestry switches in African-Americans (*42*), which is highly correlated with other maps (*42*) but is least dependent on diversity levels.

### Background selection

We first focus on two of our best-fitting models of the effects of background selection (see below and SOM Section 4). In both cases, we take as putative targets of purifying selection the 6% of autosomal sites estimated as most likely to be under selective constraint. In one, we choose these sites using phastCons conservation scores obtained for a 99-vertebrate phylogeny that excludes humans (*43*). In the other, we rely on Combined Annotation-Dependent Depletion (CADD) scores, which are based primarily on phylogenetic conservation (excluding humans) but also on information from functional genomic assays (*44, 45*); to avoid circularity, we use scores that were generated without the McVicker et al. (*29*) *B*-map as input (see SOM Section 2.5). From these models, we obtain a map of predicted diversity levels (accounting for variation in mutation rates), which we can then compare to observed data (Figs. 2A and S24). As a measure of the precision of our predictions, we consider the variance in diversity levels explained in non-overlapping autosomal windows (Fig. 2B). Our predictions explain a large proportion of the variance across spatial scales: at the 1 Mb scale, the predictions based on CADD scores account for 59.9% of the variance in diversity levels compared to 32% explained by previous work (*29*) (see SOM Section 4.6).

**Fig. 2.**
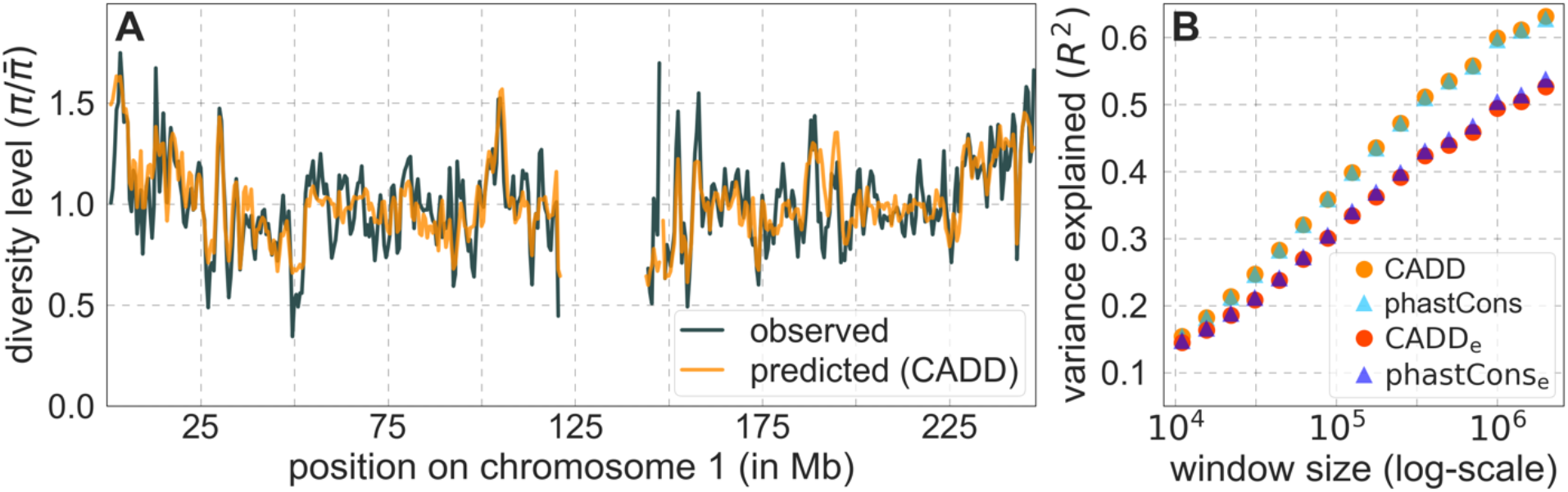
Comparison of diversity levels predicted by our best-fitting maps of background selection effects with observations. (**A**) Predicted and observed diversity levels along chromosome 1 in the YRI sample. Diversity levels are measured in 1 Mb windows, with a 0.5 Mb overlap, with the autosomal mean set to 1. (**B**) The proportion of variance in YRI diversity levels explained by background selection models at different spatial scales. Shown are the results for four choices of putative targets of selection: all sites with the highest 6% of CADD or phastCons scores (denoted CADD and phastCons, respectively) and the subset of these sites that are exonic (denoted CADD_ex_ and phastCons_ex_, respectively). See SOM Section 4 for similar graphs with other choices, and SOM Sections 7 and 9 for other populations. We show that our estimates are not inflated by over-fitting in SOM Section 6.

### Selective sweeps

Next, we examine whether incorporating selective sweeps alongside background selection improves our predictions. Our inference should be able to tease apart the effects of selective sweeps, primarily because their effects, unlike those of background selection, should be centered around the locations of substitutions. Moreover, as noted, we expect to capture the effects of selective sweeps, be they hard, partial or soft (*2, 3, 10-15*), so long as they resulted in substitutions and substantially affected diversity levels (see SOM Section D in (*30*)). Indeed, previous work that applied a similar methodology to data from *Drosophila melanogaster* was able to identify distinct effects of background selection and sweeps (*30*). To examine whether we can identify such effects in humans, we consider several choices of putatively selected substitutions along the human lineage, including any nonsynonymous substitutions or any nonsynonymous and non-coding substitutions in constrained regions, allowing each type to have its own selection parameters and considering different measures of constraint (see SOM Section 4.5). Regardless of the types of substitutions considered, incorporating sweeps does not improve our fit. In fact, in all cases, our estimates of the proportion of substitutions resulting in sweeps with discernable effects on neutral diversity is approximately 0.

Moreover, in contrast to previous attempts (*29, 35*), our model of background selection alone provides good quantitative fits to the diversity levels observed around different genomic features and in particular around nonsynonymous and synonymous substitutions (Figs. 3 and S48). Together, these results refute the hypothesis that reduced diversity levels around nonsynonymous substitutions in humans reflect ‘masked’ effects of selective sweeps (*36*); more generally, they indicate that selective sweeps resulting in substitutions had little effect on diversity levels in contemporary humans.

**Fig. 3.**
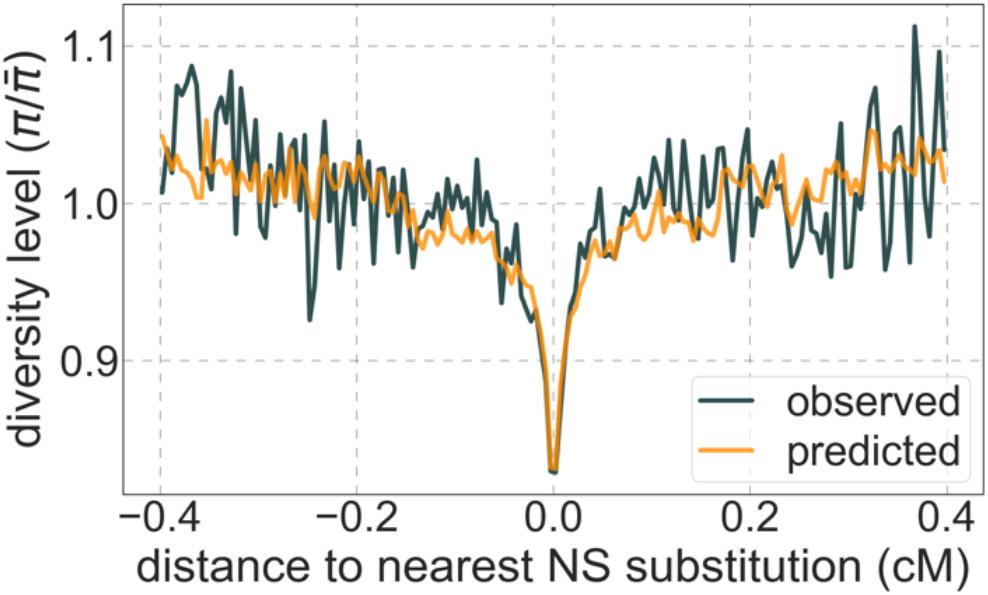
A background selection model predicts neutral diversity levels observed around human-specific nonsynonymous (NS) substitutions. Shown are the results for putatively neutral sites as a function of their genetic distance to the nearest nonsynonymous substitution (in 160 bins, each spanning 0.005cM). For observed values, we average diversity levels within each bin. For predicted values, we average diversity levels predicted by our best-fitting CADD-based model and correct for relative mutation rate in each bin (using substitution data; see SOM Section 3.3). Both observed and predicted diversity levels are plotted relative to the autosomal mean. See Figs. S48 and S50 for similar graphs for other genomic features and using data from other populations.

The lack of a footprint of sweeps in neutral diversity data does not imply that adaptation was rare in recent human evolution. Instead, much of it may have been driven by selection on genetically complex traits (*34, 35, 46-48*), i.e., traits with heritable variation arising from many segregating loci. The response to a shift in selection pressures on complex traits is expected to be highly polygenic, with short-term, rapid but tiny changes to allele frequencies at many loci that segregated before the shift, and a longer term, small excess in the fixation of alleles that change the trait in the direction favored by selection (*49*). Importantly, polygenic adaptation is unlikely to result in sweeps and is likely to introduce only minor perturbations to the allele trajectories expected when selection pressures on traits remain constant (*49*), implying that its effects on neutral diversity levels should be minor (*50, 51*). In contrast, ongoing stabilizing selection on complex traits, i.e., selection acting to maintain traits near an optimal value, is thought to be common (*48, 52-54*) and could have a substantial effect on neutral diversity levels (*49*). Stabilizing selection induces purifying selection against minor alleles that affect complex traits (*53, 55, 56*), and purifying selection on these alleles could be a major source of background selection (*49*). In other words, if much of the selection in humans is driven by ongoing and changing selection pressures on complex traits, we may expect background selection to be the dominant mode of linked selection, as our results indicate.

### The source of background selection

Focusing then on models of background selection alone, we ask which genomic annotations appear to be the sources of purifying selection. Previous work found selection on non-exonic regions to contribute little, to the extent that removing conserved non-exonic sites from a model of background selection had little effect on predicted diversity levels (*29*). In contrast, when we include only conserved exonic regions in our inference, our predictive ability is considerably diminished (Fig. 2B).

Moreover, in models that include separate selection parameters for conserved exonic and non-exonic regions, purifying selection on non-exonic regions accounts for most of the reduction in linked neutral diversity (SOM Section 4.3). Our estimates suggest that ∼80% of deleterious mutations affecting neutral diversity occur in non-exonic regions (e.g., in the model with the top 6% of phastCons scores, ∼84% of selected sites and ∼76% of mutations are non-exonic; with the top 6% of CADD scores, ∼83% of selected sites and ∼85% of deleterious mutations are non-exonic; see SOM Sections 4.3 and 4.6). Our estimates of the average strength of selection differ between exonic and non-exonic regions, but because the total reduction in diversity levels caused by background selection is fairly insensitive to the strength of selection (with the reduction being more localized for weakly selected mutations than for strong ones), the proportions of deleterious mutations that occur in these regions approximate their relative effects on neutral diversity levels (*57*) (see SOM Sections 4.3, 4.4 and 4.6). Thus, our estimates suggest that purifying selection on non-exonic regions accounts for ∼80% of the reduction in linked neutral diversity. Moreover, including separate selection parameters for conserved exonic and non-exonic regions does not improve our predictions (SOM Section 4.3 and Fig. S19).

Incorporating additional functional genomic information also does little to improve our predictions (SOM Sections 4.2 and 4.4). Notably, when we do not incorporate information on phylogenetic conservation, but include separate selection parameters for coding regions and for each of the Encyclopedia of DNA Elements (ENCODE) classes of candidate cis-regulatory elements (cCRE) (*58*), our predictive ability is considerably diminished (SOM Section 4.4). Moreover, using CADD scores (*44, 45*), which augment information on phylogenetic conservation with functional genomic information, offers little improvement over relying on conservation alone (e.g., explaining 59.9% compared to 59.7% of the variance in diversity levels in 1Mb windows, a difference that is not statistically significant; SOM Section 6). Thus, at present, functional annotations that do not incorporate phylogenetic conservation appear to provide poorer predictions of the effects of linked selection and those that do offer little improvement over conservation alone (see SOM Sections 4.1-4).

In turn, our predictions based on conservation are fairly insensitive to the phylogenetic depth of the alignments used to infer conservation levels, although we do slightly better using a 99-vertebrate alignment (excluding humans) compared to its monophyletic subsets (e.g., Figs. S14 and S33 and SOM Section 6.2). Our best-fitting models by a variety of metrics, are obtained using 5-7% of sites with the top CADD or phastCons scores as selection targets (Figs. S16 and S26). This percentage is in good accordance with more direct estimates of the proportion of the human genome subject to functional constraint (*59, 60*).

### Estimates of the deleterious mutation rate

Reassuringly, the deleterious mutation rates that we estimate for our best-fitting models are plausible (Fig. 4). Current estimates of the average mutation rate per site per generation in humans, including point mutations (*32, 33*), indels (*33*), mobile element insertions (*61*), and structural mutations (*62, 63*) lie in the range of 1.29 × 10^−8^ − 1.38 × 10^−8^ per base pair per generation (SOM Section 5). Further accounting for the length of deletions (*33*)—whereby a deletion that starts at a neutral site and includes selected sites should contribute to the our estimate of the deleterious mutation rate, but deletions that affects one or several selected sites should have the same contribution—suggests that the upper bound on estimates of the deleterious mutations rate at putatively selected sites should fall in the range of 1.29 × 10^−8^ − 1.51 × 10^−8^ per base pair per generation (SOM Section 5). The estimates for all of our best-fitting models fall well below this bound (Fig. 4). This is expected, because not every mutation at putatively selected sites will be deleterious: some sites are misclassified as constrained and some mutations at selected sites are selectively neutral.

**Fig. 4.**
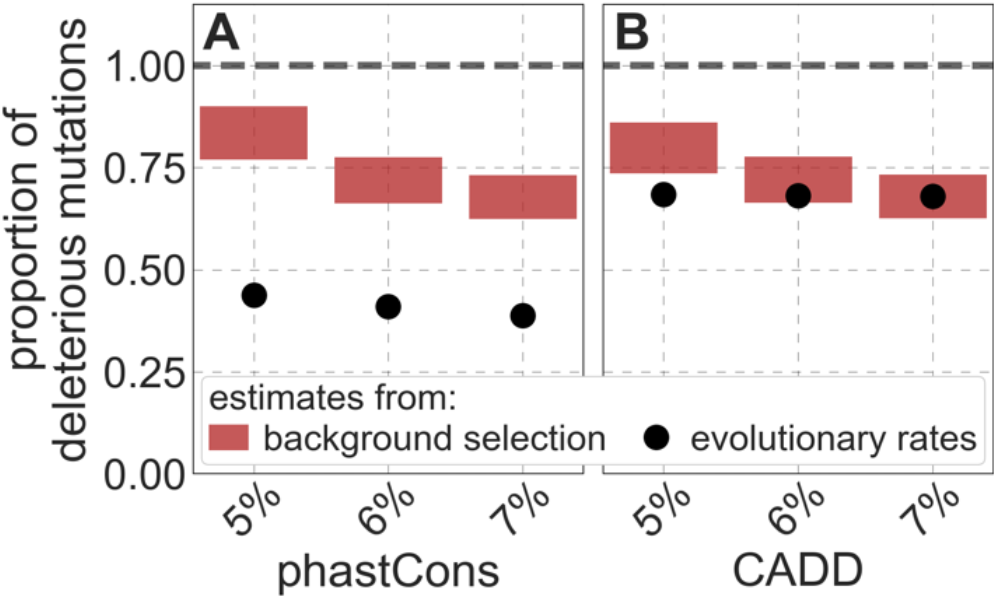
Estimates of the proportion of mutations at putatively selected sites that are deleterious. Shown are the results using 5-7% of sites with the highest phastCons scores (**A**) and CADD scores (**B**) as selection targets. For estimates based on fitting background selection models, we divide our estimates of the deleterious mutation rate per selected site by the estimate of the total mutation rate per site, where the ranges correspond to the range of estimates of the total rate, i.e., 1.29 × 10^−8^ − 1.51 × 10^−8^ per site per generation (SOM Section 5.1). For estimates based on evolutionary rates (on the human lineage from the common ancestor of humans and chimpanzees), we take the ratio of the estimated rates at putatively selected sites and at matched sets of putatively neutral sites (see text and SOM Section 5.2 for details).

To test whether our estimates of the proportion of mutations that are deleterious are plausible, we compare them with independent estimates based on the relative reduction in evolutionary rates at putatively selected vs. neutral sites along the human lineage (these sets of sites were identified from an alignment that excludes humans; SOM Sections 3.1, 4.1 and 4.4). The relative reduction allows us to estimate the proportion of deleterious mutations because deleterious mutations at selected sites rarely fix in the population whereas neutral mutations fix at a much higher rate, which is the same at selected and neutral sites (*64*). In estimating the reduction at putatively selected sites, we matched the set of putatively neutral sites for the AT/GC ratio, and checked that our estimates were insensitive to the composition of other genomic features associated with mutation rates and with other non-selective processes that affect substitution rates (e.g., triplet context, methylated CpGs and recombination rates, which affect rates of biased gene conversion; SOM Section 5).

Our estimates based on evolutionary rates are closer to (and even overlap) those obtained from fitting models of background selection based on CADD scores compared to those based on phastCons scores (Fig. 4). This is expected given that CADD scores are much better than phastCons scores at identifying constraint on a single site resolution (*44, 45*), which markedly influences evolutionary rates at putatively selected sites (but not the predictions of background selection effects). We expect the two estimates to be similar but not identical, both because weak selection has a larger effect on evolutionary rates than on linked diversity levels (*8, 16, 17, 65, 66*) and because estimates based on the effects of background selection may absorb the deleterious mutation rate at selected sites that were not included in our sets but are closely linked to sites in them (SOM Section 5). In summary, given the fit to data and plausible estimates of the deleterious rates, it is natural to interpret our maps as reflecting the effects of background selection, i.e., as maps of *B* (defined as the ratio of expected diversity levels with background selection, π, and in its absence, π_0_) (*5*).

### Background selection on autosomes

Our maps are also well calibrated (Fig. 5). When we stratify diversity levels at putatively neutral sites by our predictions, predicted and observed diversity levels are similar throughout nearly the entire range of predicted values (e.g., *R*^2^ = 0.96 when sites are in predicted percentile bins). One exception is for ∼4% of sites in which background selection is predicted to be the strongest (i.e., with the lowest *B*), where our predictions are imprecise. This behavior is due to a technical approximation we employ in fitting the models (see SOM Section 1.5). The other exception is for ∼2% of sites in which background selection is predicted to be the weakest (i.e., with *B* near 1), where observed diversity levels are markedly greater than expected. We observe similar behavior in all the human populations examined (Fig. S51), and we cannot fully explain it by known mutational and recombination effects (e.g., of base composition and biased gene conversion; SOM Section 8). This behavior could reflect ancient introgression of archaic human DNA into ancestors of contemporary humans (SOM Section 8.3), indicated also in other population genetic signatures (*67-73*). Such introgressed regions are expected to increase genetic diversity and persist the longest in regions with low functional density and high recombination, corresponding to weak background selection effects (*70, 74, 75*).

**Fig. 5.**
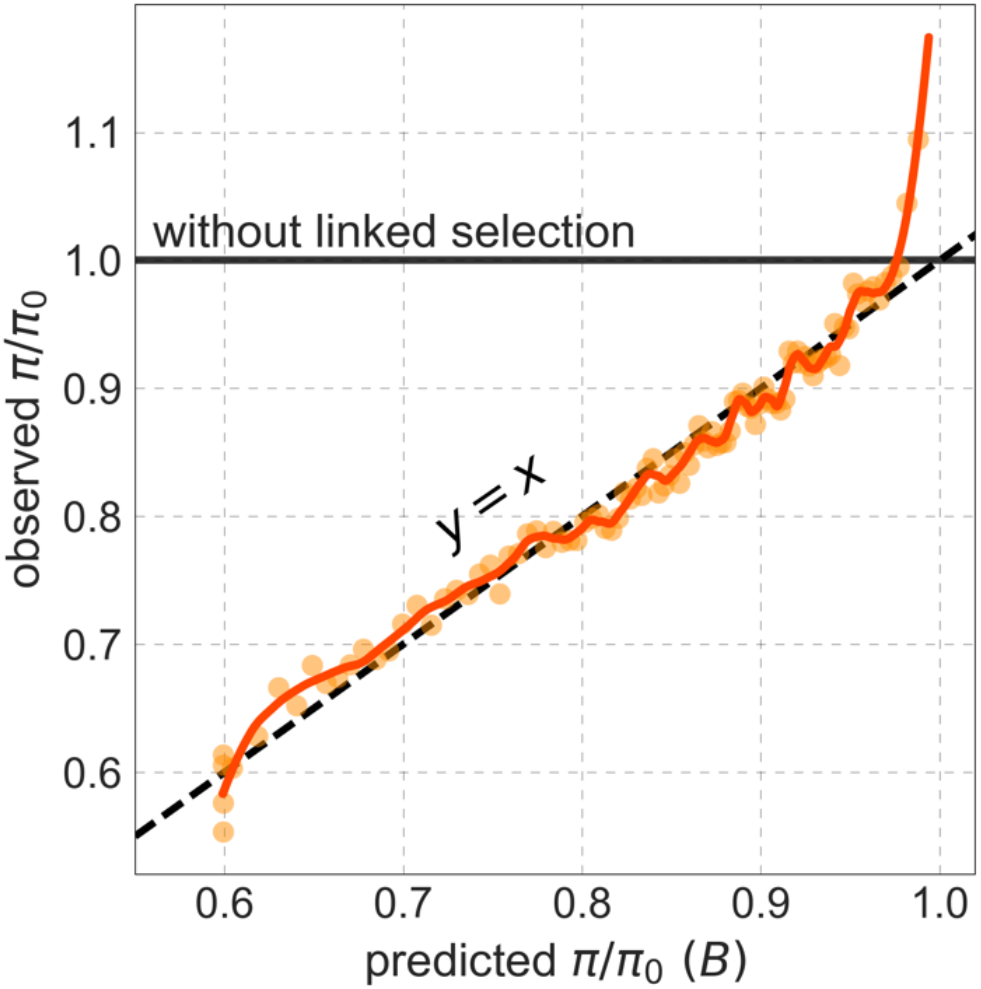
Observed vs. predicted neutral diversity levels across the autosomes. Shown are the results for the best-fitting CADD-based model. Light orange scatter plot: we divide putatively neutral sites into 100 equally sized bins based on the predicted *B*. For predicted values (x-axis), we average the predicted *B* in each bin. For observed values (y-axis), we divide the average diversity level by the estimate of the average relative mutation rate (obtained from substitution data; see SOM Section 3.3) in each bin, and normalize by the autosomal average of π_0_ (estimated from fitting the model; see SOM Section 1.1). Dark orange curve: the LOESS fit for a similarly defined scatter plot but with 2000 rather than 100 bins (with span=0.1). For similar graphs corresponding to other models and using data from other populations, see SOM Sections 4 and 9, respectively.

Setting these outlier regions aside, we can use the maps to characterize the distribution of background selection effects in human autosomes. We note that background selection effects that are not captured by our models would cause us to underestimate the range and extent of background selection effects (*30*). We find that diversity levels throughout almost all of the autosomes are affected by background selection, with a ∼37% reduction in the 10% most affected sites, a non-zero (∼2.1%) reduction even in the 10% least affected (after excluding outliers in the top 2% of bins; see Fig. 5), and a mean reduction of ∼17%. These conclusions are robust across our best-fitting maps and populations (SOM Section 4 and Figs. S35 and S51). An important implication is that our maps of the effects of background selection provide a more accurate null model than currently used for other population genetic inferences that rely on diversity levels, notably inferences about demographic history (*76, 77*).

## Conclusion

Our results indicate that background selection is the dominant mode of linked selection in human autosomes and the major determinant of neutral diversity levels on the Mb scale. They further reveal that background selection effects arise primarily from purifying selection at non-coding regions of the genome. Non-coding regions are known to exhibit substantial functional turnover on evolutionary timescales (*59, 60*), and yet we find phylogenetic conservation to be the best predictor of selected regions. Moreover, at present, augmenting measures of conservation with functional genomic information in humans offers little improvement. It therefore remains unclear how much our maps can still be improved. Even without these potential refinements, our findings demonstrate that a simple model of background selection, conceived nearly three decades ago (*5*), is able to provide a reliable quantitative prediction of genetic diversity levels throughout human autosomes.

## Acknowledgements

We thank M. Przeworski for helpful discussions throughout this work. We also thank Ipsita Agarwal, Eduardo Amorim, Peter Andolfatto, Ian Mathieson, Priya Moorjani, Itsik Pe’er, Joe Pickrell and Jonathan Pritchard for helpful discussions. We thank Yun Song for sharing unpublished results, and Lusiné Nazaretyan, Philipp Rentzsch, Max Schubach and Martin Kircher from the Kircher lab for generating CADD scores that were tailored for our purposes. We thank Ipsita Agarwal, Peter Andolfatto, Jonathan Pritchard and Molly Przeworski for comments on the manuscript. This work was funded by NIH grant GM115889 to GS and NIH training grant T32GM008798 to DAM.

