## Supplementary Material for "Broad-scale variation in human genetic diversity levels is predicted by purifying selection on coding and non-coding elements"

### 1. Model and inference method

Here we detail the model and inference method used in this study. In Section 1.1, we describe our model for the effects of background selection and selective sweeps and our approach to inferring the parameters of these models. This section is adapted from Elyashiv and colleagues (*1*), who applied a similar approach to data from *Drosophila melanogaster*; we reproduce it here for completeness. In Section 1.2, we describe how we calculate lookup tables for the effects of background selection and sweeps, which our inference relies upon. We introduce several changes to the methods used in previous studies (*1, 2*), which allow us to better control the precision of maps of the effects of linked selection. In Section 1.3, we describe how we represent neutral polymorphism data and maps of the effects of linked selection in our calculations in order to increase computational tractability. In Section 1.4, we describe the optimization algorithm that we use to find the selection parameters that maximize our models composite-likelihood, and we apply the optimization to simulated datasets in order to demonstrate its efficacy and robustness. In Section 1.5, we introduce a thresholding approach that contends with biases in our optimization that arise from model misspecification, and we investigate how this thresholding affects our inferences. Finally, in Section 1.6, we provide an overview of the software that we use for inference and for other key analyses in the paper. The software, its documentation, and maps of the effects of linked selection are available for download at (github.com/sellalab/HumanLinkedSelectionMaps).

#### 1.1 Model and inference problem

We model the effects of background selection and selective sweeps on neutral heterozygosity levels (i.e., the probability of observing different alleles in a sample size of two), *π*, at an autosomal position *x*. In a coalescent framework, the model takes the form

| $\pi\left( x \right)=\frac{2u(x)}{2u\left( x \right)+1/(2N_{e}B\left( x \right))+S(x)},$ | $(1)$ |
| --- | --- |

where $u(x)$ is the local mutation rate, $N_{e}$ is the effective population size without linked selection, $B\left( x \right)$ is the local (multiplicative) reduction in the effective population size due to background selection and $S(x)$ is the local coalescence rate caused by selective sweeps (*1, 3*). This approximation can be derived by considering the probability that a mutation occurs (at a rate $2u(x)$ per generation) before the pair of lineages coalesces, owing either to genetic drift ($1/2N_{e}B\left( x \right)$), which includes the effect of background selection, or to a selective sweep ($S(x)$). While we consider autosomes, the model can be extended to sex chromosomes with straightforward modifications.

The model for the effects of background selection, $B\left( x \right)$, follows Hudson & Kaplan (*4*) and Nordborg et al. (*5*) (Fig. S1a). We assume a set of distinct annotations $i_{B}=1,\ldots I_{B}$ under purifying selection (e.g., conserved exonic and non-exonic regions) and positions in the genome $A_{B}=\left\{ a_{B}\left( i_{B} \right) | i_{B}=1,\ldots,I_{B} \right\},$ where $a_{B}(i_{B})$ denotes the set of genomic positions with annotation $i_{B}$**.** The selection parameters at these annotations are given by $\Theta_{B}=\{(u_{d}\left( i_{B} \right), f\left( t | i_{B} \right))|i_{B}=1,\ldots,I_{B}\}$, where $u_{d}$ is the rate of deleterious mutations and $f(t)$ is the distribution of selection coefficients in heterozygotes for a deleterious mutation. The reduction in the effective population size is then

| $B\left( x \vert A_{B}, \Theta_{B}, R \right)=\mathrm{Exp}\left( -\sum_{i_{B}} \sum_{y\in a_{B}\left( i_{B} \right)} \int\frac{u_{d}\left( i_{B} \right)}{t\left( 1+{r\left( x,y \right)\left( 1-t \right)}/t \right)^{2}}f\left( t \vert i_{B} \right)dt \right),$ | $(2)$ |
| --- | --- |

where $R$ is the genetic map and $r\left( x,y \right)$ is the genetic distance between the focal position $x$ and positions $y$ (only positions on the same chromosome are considered). The integrand reflects the effect that a site under purifying selection at position $y$ exerts on a neutral site at position $x$. This expression and its combination across sites provide a good approximation to the effect of background selection so long as selection is sufficiently strong (i.e., when$2N_{e}t\gg1$).

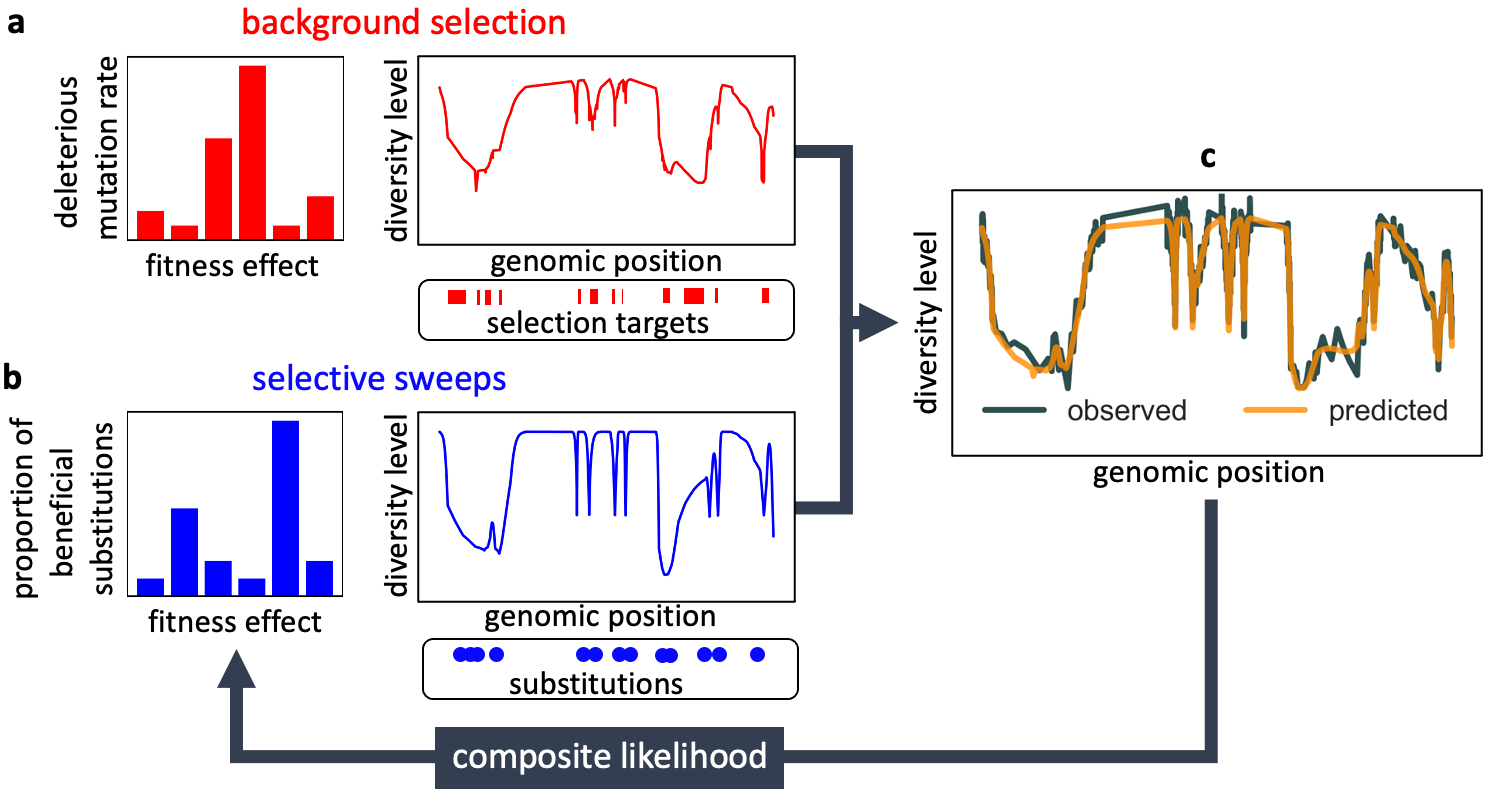

**Figure S1**. Modeling and inferring the effects of linked selection in humans. Given the targets of selection and corresponding selection parameters (a and b), we calculate the expected neutral diversity levels along the genome (c). We infer the selection parameters by maximizing their composite-likelihood given observed diversity levels (c). Based on these parameter estimates, we calculate a map of the expected effects of linked selection on diversity levels.

In turn, the model for the effect of selective sweeps follows from an approximation used by Barton (*6*) and Gillespie (*7*), among others (Fig. S1b). Similarly to the model for background selection, we assume a set of distinct annotations $i_{S}=1,\ldots,I_{S}$ subject to sweeps, but here the specific positions at which substitutions have occurred are known, $A_{S}=\{a_{S}(i_{S})|i_{s}=1,\ldots,I_{S}\}$ with $a_{S}\left( i_{S} \right)$ denoting the set of substitution positions with annotation $i_{S}$**.** The selection parameters at these annotations are $\Theta_{S}=\{(\alpha\left( i_{S} \right), g\left( s | i_{S} \right))|i_{s}=1,\ldots,I_{S}\}$, where $\alpha$ is the fraction of substitutions that are beneficial and $g(s)$ is the distribution of their additive selection coefficients. For autosomes, the expected rate of coalescence per generations at position $x$ due to sweeps is then approximated by

| $S\left( x \vert{A_{S}, \Theta}_{S}, R, \bar{N}_{e}, T \right)=\frac{1}{T}\sum_{i_{S}} \alpha(i_{S})\sum_{y\in a\left( i_{S} \right)} \int\mathrm{Exp}\left( -r(x,y \right)\tau(s,\bar{N}_{e}))g(s\left\vert i_{S} \right)ds,$ | $(3)$ |
| --- | --- |

where $T$ is the length of the lineage (in generations) over which substitutions occurred, the positions of substitutions $y$ are summed over the chromosome with the focal site, $\bar{N}_{e}$ is the average effective population size and $\tau(s,\bar{N}_{e})$ is the expected time to fixation of a beneficial substitution with selection coefficient $s$ and given an effective population size $\bar{N}_{e}$. We use the diffusion approximation for the fixation time

| $\tau\left( s,N_{e} \right)=\frac{2(\ln\left( 4N_{e}s \right)+\gamma-\left( 4N_{e}s \right)^{-1}}{s},$ | $(4)$ |
| --- | --- |

where $\gamma$ is the Euler constant (*8*). This model relies on several simplifying assumptions and approximations. In particular, the term $1/T$ relies on an assumption of one substitution per site per lineage and neglects variation in the length of lineages across loci. In combining the effects over substitutions, we further assume that the timings of beneficial substitutions are independent and uniformly distributed along the lineage, and that they are infrequent enough such that we can ignore interference among them (*9*). The exponent approximates the probability of coalescence of two samples due to a classic sweep with additive selection coefficient $s$ (where $2N_{e}s\gg1$) in a panmictic population of constant effective size $\bar{N}_{e}$. (For the relationships between these expressions and other kinds of sweeps see SOM Section D in (*1*)). In principle, we should use the local $N_{e}$ incorporating the effects of background selection but given the logarithmic dependence of Equation (4) on $N_{e}$, we simply use the average $\bar{N}_{e}$.

To infer the selection parameters $\Theta_{B}$ and $\Theta_{S}$, we use a composite-likelihood approach across sites and samples (*10*) (Fig. S1). We denote the positions of neutral sites by $X$ and the set of samples by $I$. We then summarize the observations by a set of indicator variables across sites and all pairs of samples $O=\{O_{i,j}(x)|x\in X, i\neq j\in I\}$, where $O_{i,j}\left( x \right)=1$ indicates that samples $i$ and $j (j\neq i)$ differ at position $x$ and $O_{i,j}\left( x \right)=0$ indicates that they are the same. In these terms, the composite log-likelihood takes the form

| $\log\left( L \right)=\sum_{x\in X} \sum_{i\neq j\in I} log(Pr\{O_{i,j}\left( x \right)\vert\Theta_{B},\Theta_{S}\}),$ | $(5)$ |
| --- | --- |

where

$$\Pr\left\{ O_{i,j}\left( x \right) | \Theta_{B},\Theta_{S} \right\}=\left\{ \begin{aligned} \pi\left( x | \Theta_{B}, \Theta_{S} \right) O_{i,j}\left( x \right)=1 \\ 1-\pi\left( x | \Theta_{B}, \Theta_{S} \right) O_{i,j}\left( x \right)=0 \end{aligned} \right.$$

Using composite-likelihood circumvents the complications of considering linkage disequilibrium (LD) and of coalescent models for larger sample sizes. Importantly, maximizing this composite-likelihood should yield unbiased point estimates (*11, 12*). Beyond losing the information in LD patterns and in the site frequency spectrum, the main cost of this approach is the difficulty in assessing uncertainty in parameter estimates (as standard asymptotic results do not apply). We therefore use other ways to assess the reliability of our inferences.

To make the composite-likelihood calculations (i.e., the calculation of $\pi\left( x | \Theta_{B}, \Theta_{S} \right)$) feasible genome-wide, we discretize the distribution of selection coefficients on a fixed grid. Given a grid of negative and positive selection coefficients, $t_{g}$ and $s_{k}$, $g=1,\ldots G$ and $k=1,\ldots K$, the distribution of selection coefficients for each annotation becomes a set of weights on this grid, $w\left( t_{g} | i_{B} \right)$ and $w(s_{k}|i_{S})$. (In principle, the grid could also be annotation-specific.) For background selection, these weights reflect the rate of deleterious mutations with a given selection coefficient and their sum should therefore be bound by the maximal deleterious mutation rate per site. For sweeps, the weights reflect the fraction of beneficial substitutions with a given selection coefficient and their sum should be bound by 1. In these terms, the effect of background selection takes the form

| $B\left( x \vert\Theta_{B} \right)=\mathrm{Exp}\left( -\sum_{i_{B}} \sum_{g=1}^{G} w(t_{g}\vert i_{B})b(x\vert t_{g}, i_{B}) \right),$ | $(6)$ |
| --- | --- |

where $Exp\left( -b(x|t_{g}, i_{B}) \right)$ is the proportional reduction in the effective population size induced by having one deleterious mutation per generation per site with selection coefficient $t_{g}$ at all the positions in annotation $i_{B}$. By the same token, the effects of sweeps take the form

| $S(x\vert\Theta_{S})=\frac{1}{T}\sum_{i_{S}} \sum_{k=1}^{K} w\left( s_{k} \vert i_{S} \right)s\left( x \vert s_{k}, i_{S} \right),$ | $(7)$ |
| --- | --- |

where $\frac{1}{T}s\left( x | s_{k}, i_{S} \right)$ is the probability of coalescence per generation induced by sweeps in annotation $i_{S}$, if all the substitutions in this annotation are beneficial with selection coefficient $s_{k}$. By using a grid, we can calculate a lookup table of $b(x|t_{g}, i_{B})$ and $s\left( x | s_{k}, i_{S} \right)$ once and then use it repeatedly to calculate the likelihood of different sets of weights. Moreover, the interpretation of estimated distributions on a grid is arguably simpler than that of the continuous parametric distributions commonly used (e.g., gamma and exponential), which impose rigid interdependencies between the densities associated with different selection coefficients with little justification and while the data is only informative about a subset of the domain. In the next section, we describe additional simplifications in the calculation of $b(x|t_{g}, i_{B})$ and $s\left( x | s_{k}, i_{S} \right)$.

Other parameters are estimated as follows. Consider equation (1) rewritten as

| $\pi\left( x \right)=\frac{\pi_{0}\cdot(u(x)/\bar{u})}{\pi_{0}\cdot(u(x)/u ̅ )+1/B(x) +2N_{e}S(x;\bar{N}_{e},T)},$ | $(8)$ |
| --- | --- |

to clearly specify all the additional parameters required for inference.$\pi_{0}\equiv4N_{e}\bar{u}$ is (approximately) the average neutral heterozygosity, given the effective population size in the absence of linked selection and the average mutation rate per site ($\bar{u}$); $\pi_{0}$ is estimated through the likelihood maximization. The local variation in mutation rate $u(x)/\bar{u}$ is estimated based on substitution rates at putatively neutral sites in an eight-primate phylogeny (excluding humans) in nonoverlapping windows, with a window size chosen to balance true variation in mutation rates and measurement error (see Section 3.3). Finally, $\bar{N}_{e}$ is estimated based on the average genome-wide heterozygosity at putatively neutral sites, after dividing out by a direct estimate of the spontaneous point mutation rate of $1.2\times{10}^{-8}$ per site per generation (*13*), and ${T/2\bar{N}}_{e}$ is estimated by $(\bar{K}/2)/\pi_{0}$, where $\bar{K}$ is the average number of point substitutions per putatively neutral site on the human lineage (see Section 2.7).

#### 1.2 Calculating lookup tables

Here we describe how we calculate the lookup tables for

| $s\left( x \vert s_{k}, i_{S} \right)\equiv\sum_{y\in a\left( i_{S} \right)} \mathrm{Exp}\left( -r(x,y \right)\tau(s_{k},\bar{N}_{e}))$ | $(9)$ |
| --- | --- |

and

| $b(x\vert t_{g}, i_{B})\equiv\sum_{y\in a_{B}\left( i_{B} \right)} \frac{1}{t_{g}\left( 1+r\left( x,y \right)\left( 1-t_{g} \right)/{t_{g}} \right)^{2}}$ | $(10)$ |
| --- | --- |

at all putatively neutral autosomal positions ($x$), given annotations ($i_{B}$and $i_{S}$) and selection coefficients ($t_{g}$ and $s_{k}$). We focus on one annotation and selection coefficient at a time and therefore simplify the notation to $b(x)$ and $s(x)$, and omit the variables in $\tau$ and the subscripts of the selection coefficients. When we refer to accuracy in this section, we assume that there is no model misspecification (e.g., that putatively neutral sites are neutral, that sets of selected sites and selection parameter values are accurate, that genetic maps are accurate, etc.); once we control the accuracy in this sense, the main sources of error in our predictions will be due to model misspecification.

*Our general approach is to calculate* $b(x)$ *and* $s(x)$ *with high accuracy at a subset of positions and to use linear interpolation between them.* The distances between these positions are chosen such that maps built using the lookup tables maintain a preset level of accuracy $\epsilon$. Specifically, we require that our approximation $\tilde{s}$ and $\tilde{b}$ at any position $x$ satisfy

$\left| \frac{\tilde{s}\left( x \right)-s(x)}{s(x)} \right|<\epsilon$ and $\left| \frac{\mathrm{Exp}\left( -u_{M}\cdot\tilde{b}\left( x \right) \right)-\mathrm{Exp}\left( -u_{M}\cdot b\left( x \right) \right)}{\mathrm{Exp}\left( -u_{M}\cdot b\left( x \right) \right)} \right|<\epsilon$,

where $u_{M}$ is an upper bound on the deleterious mutation rate per site per generation. When these conditions are met one can show (based on Eqs. 6 and 7) that the relative accuracy of $S$ and $B$, and consequently of the expected neutral diversity level $\pi$ (based on Eq. 1), are also bound by $\epsilon$.

**Sweeps.** Assume that we have calculated $s$ accurately at position $x$ and consider the distance $\Delta$ at which the relative change in $s$ is bound by $\epsilon$, i.e., where

| $\left\vert\frac{s\left( x+\Delta\right)-s(x)}{s\left( x+\Delta\right)} \right\vert\leq\epsilon.$ | $(11)$ |
| --- | --- |

From Eq. 11, we find that

$$\left| s\left( x+\Delta\right)-s\left( x \right) \right|\leq\sum_{y} \left| \mathrm{Exp}\left( -r\left( x+\Delta,y \right)\cdot\tau\right) -\mathrm{Exp} \left( -r\left( x,y \right)\cdot\tau\right) \right|$$

$$=\sum_{y} \mathrm{Exp}\left( -r\left( x+\Delta,y \right)\cdot\tau\right)\cdot\left| 1 -\mathrm{Exp} \left( \left( r\left( x+\Delta,y \right)-r\left( x,y \right) \right)\cdot\tau\right) \right|$$

$$\approx\sum_{y} \mathrm{Exp}\left( -r\left( x+\Delta,y \right)\cdot\tau\right)\cdot\left| 1 -\mathrm{Exp} \left( r\left( x+\Delta,x \right)\cdot\tau\right) \right|$$

$\approx s(x+\Delta)\cdot\left( r\left( x+\Delta,x \right)\cdot\tau\right)$,

where the approximations assume $r\left( x+\Delta,x \right)\ll1$. Consequently, by solving for $\Delta$ such that

| $r\left( x+\Delta,x \right)=\epsilon/\tau$ | $(12)$ |
| --- | --- |

we assure that the relative accuracy between $x$ and $x+\Delta$ is bound by $\epsilon$. We therefore calculate $s$ at the selected set of positions on a chromosome beginning at one end and choosing our step sizes according to Eq. (12) until we reach the other end.

**Background selection.** Our calculation for background selection is based on the algorithm developed by McVicker et al. (*2*) (their calc_bkgd program) with several important modifications (Fig. S2). The problems that require these modifications are most pronounced for small selection coefficients, whose background selection effects are localized at short genetic distances from selected segments where they can be quite strong. First, McVicker et al. used an additional lookup table to integrate over the effects of background selection exerted by a contiguous selected segment (SI of (*2*)). This lookup table had poor resolution for small selection coefficients at short genetic distances from selected segments, and we have increased the resolution accordingly to fix the problem. Second, the algorithm for choosing the step size $\Delta$ is designed to control the absolute error, such that

$\left| \mathrm{Exp}\left( -u_{M}\cdot\tilde{b}\left( x \right) \right)-\mathrm{Exp}\left( -u_{M}\cdot b\left( x \right) \right) \right|<\epsilon$,

rather than the relative error (Eq. 13), which results in large relative errors when background selection effects are the strongest (which is with small selection coefficients). Third, the choice of step size $\Delta$ is based on the local behavior of background selection at the previous position, and consequently it sometimes skips over selected segments largely ignoring their highly localized effects (which are due to small selection coefficients). We describe how we resolve the last two problems in turn.

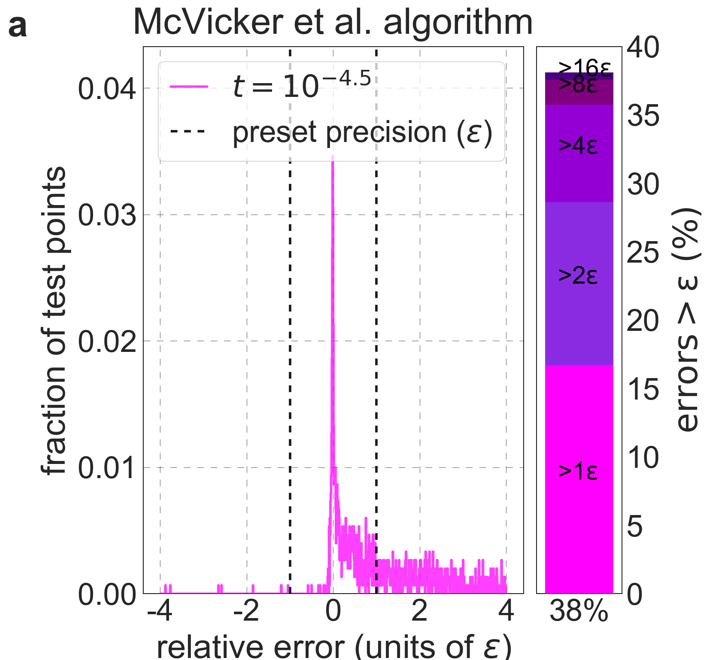

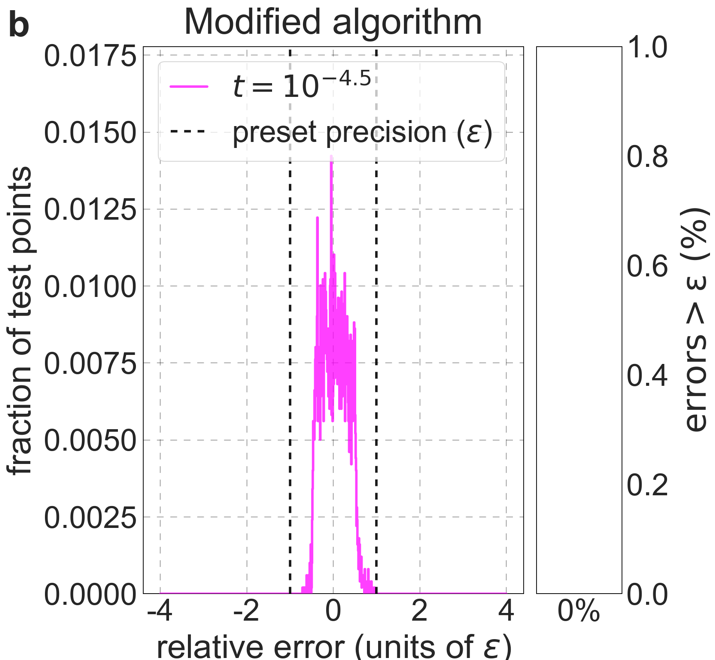

**Figure S2.** Distribution of relative errors in predictions before and after modifying calc_bkgd. We consider the model in which autosomal sites with the top 6% of CADD scores are chosen as selection targets, the deleterious mutation rate is $u_{d}={7.4\cdot10}^{-8}$per bp per generation and the selection coefficient is the lowest in our grid ($t={10}^{-4.5}$), because this is the case most prone to errors (see text). We calculate $B$-values accurately (using Eq. 10) at a million positions picked randomly from the 22 autosomes and use these values to calculate the relative errors based on the McVicker et al. algorithm (a) and on our modified algorithm (b). The side panel shows the proportion of sites in which the error exceeds $\epsilon$ (below), as well as its breakdown in multiples of $\epsilon$.

Assume that we have calculated $b$ accurately at position $x$ and consider the distance $\Delta$ at which the relative change in $\mathrm{Exp}\left( -u_{M}\cdot b \right)$ is bound by $\epsilon$ (see Eq. 13), i.e., where

| $\left\vert\frac{\mathrm{Exp}\left( -u_{M}\cdot b\left( x+\Delta\right) \right)-\mathrm{Exp}\left( -u_{M}\cdot b\left( x \right) \right)}{\mathrm{Exp}\left( -u_{M}\cdot b\left( x+\Delta\right) \right)} \right\vert\leq\epsilon.$ | $(13)$ |
| --- | --- |

Rearranging the left-hand side, we find that

$\left| 1-\mathrm{Exp}\left( -u_{M}\cdot\left( b\left( x \right)-b(x+\Delta) \right) \right) \right|\leq\epsilon$*,*

and assuming that $\left| u_{M}\cdot\left( b\left( x \right)-b(x+\Delta) \right) \right|\ll1$ we find that this requirement is well approximated by

$\left| b\left( x+\Delta\right)-b(x) \right|\approx\left| b^{'}\left( x \right)\cdot\Delta+b^{''}\left( x \right)\cdot{\Delta^{2}}/2 \right|\leq\epsilon/{u_{M}}$.

As our putative step size, we therefore take the (smallest) solution of the quadratic

| $\left\vert b^{'}\left( x \right)\cdot\Delta+b^{''}\left( x \right)\cdot{\Delta^{2}}/2 \right\vert=\epsilon/{u_{M}}$. | $(14)$ |
| --- | --- |

As in the case of sweeps, we calculate $b$ at a selected set of positions on a chromosome, beginning on one end and choosing our step sizes in a way that maintains the preset relative accuracy $\epsilon$ until we reach the other end. Assuming that we have calculated $b$ accurately at position $x$, our algorithm for choosing the step size consists of the following steps:

1. If $x$ is at the end of the chromosome, stop.
2. Calculate a candidate step size $\Delta^{*}$ by solving Eq. 14.
3. If $\Delta^{*}$ is greater than a preset maximal step size $\Delta_{max}$ then set $\Delta^{*}=\Delta_{max}$.
4. If there is a selected segment between positions $x$ and $x+\Delta^{*}$ then set $\Delta^{*}$ such that $x+\Delta^{*}$ is the midpoint between $x$ and the beginning of the (closest) selected segment. This step assures that we do not ‘skip’ selected segments.
5. Convert $\Delta^{*}$ from Morgans to base-pairs, rounding downwards. But if the step $\leq1$ bp then set it to 1 bp, calculate $b\left( x+\Delta^{*} \right)$, set $x$ to $x+\Delta^{*}$, and return to step 1.
6. Calculate $b\left( x+\Delta^{*} \right)$. If $\left| b\left( x+\Delta^{*} \right)-b(x) \right|>\epsilon/{u_{M}}$ then set the step size in Morgans to ${\Delta^{*}}/2$ and return to step 4. Otherwise, set $x$ to $x+\Delta^{*}$ and return to step 1.

**Interpolation and representation of lookup tables.** We calculate $b(x)$ or $s(x)$ at every autosomal position $x$ (for a given selection coefficient and selected annotation) by linear interpolation between adjacent positions at which we calculated $s$ and $b$ accurately. We then discretize the values of $b(x)$ or $s(x)$ on a linear grid of values corresponding to the preset accuracy $\epsilon$, and group together contiguous autosomal segments with the same discrete value. We intersect these segments with our list of putatively neutral sites (Section 3.1) to obtain lookup tables consisting of contiguous segments of putatively neutral sites with the same coarse-grained $s$ and $b$ values for our sets of selected annotations and selection coefficients.

#### 1.3 Binning neutral sites

A direct calculation of the composite log-likelihood function for given sets of selected annotations and selection coefficients and parameters (Eq. 5) requires that we store and access lookup tables and calculate the log-likelihood function at ~$6.5\times{10}^{8}$ putatively neutral autosomal sites (see Section 2.1). Doing so would entail high computation and memory demands in the search for selection parameters that maximize the composite-likelihood. For example, our best-fitting models of background selection (see Main Text) with a grid of 6 selection coefficients would require storing and repeatedly accessing lookup tables that amount to $6.5\times{10}^{8} \times8 \times6\approx32$GB (given a precision of $\epsilon=0.01$), and models involving multiple annotations for background selection and sweeps push the memory requirement to hundreds of GBs.

We reduce the computational and memory demands by dividing the set of putatively neutral sites into bins in which all the effects of background selection and sweeps predicted by the lookup tables and our estimates of the local (relative) mutation rate ($u(x)/\bar{u}$ in Eq. 8; Section 3.3) are identical. The composite log-likelihood function can then be calculated by summing over log-likelihood functions corresponding to bins, where the calculation per bin requires only the bin-specific parameters and bin-specific summaries of polymorphism. The number and identity of bins varies with the sets of selected annotations and selection coefficients and parameters and with the precision ($\epsilon$). For our best-fitting models, the average number of sites per bin is ~100, implying a ~100-fold reduction in demands on memory and in the number of log-likelihood calculations. For our most complex selection models (Section 4), the binning reduces memory and computational demands tenfold.

#### 1.4 Optimization

Here we describe how we developed and tested the algorithm we use in order to find the selection parameters that maximize the composite-likelihood of our different models. The high dimensional parameter space (including up to 55 parameters in the most complex model in Section 4) potentially makes this optimization problem non-trivial.

**One step optimization.** First, we tested the performance of standard optimization algorithms from the SciPy minimization toolkit (*14*). To this end, we generated polymorphism datasets based on our best-fitting model of background selection based on phastCons conservation scores, as follows:

1) We fixed the total deleterious mutation rate to $u_{d}={10}^{-8}$ per base per generation, and randomly divided it among the 6 selection coefficients of the model by sampling from a Dirichlet distribution (with $\alpha=1$). We set the expected neutral diversity level in the absence of background selection to $\pi_{0}=\pi_{YRI}$, where $\pi_{YRI}$ is a value of $\pi_{0}$ from an iteration of our best-fitting phastCons-based model using polymorphism data from the Yoruba (YRI) population (Section 2.1).
2) We generated the map of expected neutral diversity levels in autosomes given the chosen parameters. The map was represented in terms of the expected levels at each bin of putatively neutral sites (see Section 1.3).
3) We generated a polymorphism dataset corresponding to a sample size $n=108$ pairs of (haploid) autosomes by picking the number of pairwise differences in each bin such that the average diversity level in it most closely matched the level predicted by the map. The discretization step introduces small differences between average and expected diversity levels in bins.

We tested each algorithm by applying it to *10 simulated datasets*, with *3 sets of initial conditions* for each dataset, corresponding to weak, intermediate and strong background selection (with $u_{d}=5\times{10}^{-10}$, $5\times{10}^{-9}$, and $5\times{10}^{-8}$ per base per generation, respectively), and *5 randomly chosen initial conditions* in each set (with the total rate divided among the 6 selection coefficients by sampling from a Dirichlet distribution with $\alpha=1$) amounting to *150 runs*. The initial value of $\pi_{0}$ was always set to the average diversity level in the dataset $\bar{\pi}$.

None of the algorithms closely converged to the ground truth parameters in all cases. Nelder-Mead downhill simplex minimization (*15*) (NM) and Constrained Trust Region minimization (*16*) (CTR) performed the best overall, closely recovering the true parameters in ~2/3 of cases. While CTR was slightly more reliable, it was also up to ten times slower than NM. We therefore decide to combine them in order to leverage the relative strengths.

**Two-step minimization algorithm.** After some experimentation we converged on the following two-step algorithm (Fig. S3):

1) We apply NM with multiple initial conditions. For models of background selection with a single selected annotation we generate 3 sets of initial conditions, with 5 randomly chosen initial conditions per set, as we described above. For models of sweeps with a single annotation we generate the initial conditions analogously. Namely, we generate 3 sets of initial conditions corresponding to a low, intermediate and high proportion of beneficial substitutions (with $\alpha=0.0125, 0.125$ and 1, respectively) with 5 randomly chosen initial conditions per set (with the total proportion divided among selection coefficients by sampling from a Dirichlet distribution with $\alpha=1$). For models with background selection and sweeps and/or multiple annotations, we generate 3 sets of initial conditions, corresponding to the weak/low, intermediate, and strong/high categories, with 5 random initial conditions per set that are chosen similarly for each mode and annotation. In all cases, the initial value of $\pi_{0}$ is set to the average diversity level in the dataset $\bar{\pi}$.
2) We apply the CTR algorithm with a single initial condition that is chosen based on the output of the previous step. Specifically, we focus on the sets of selection parameters inferred in the 3 out of 15 initial runs that yielded the highest composite-likelihood, and use their average as our initial condition.

­
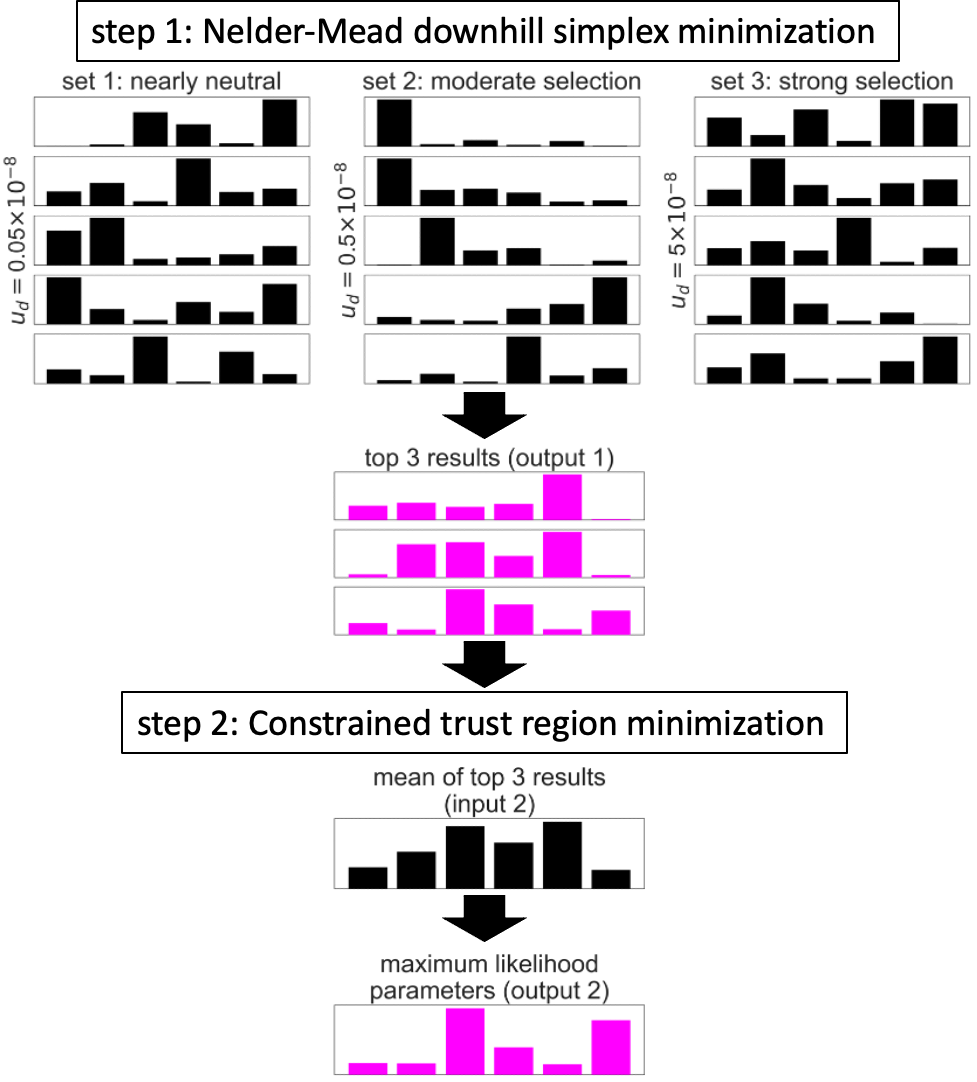

**Figure S3.** Illustration of the two-step algorithm. In this example, the optimization is applied to a model of background selection with a single selected annotation and a grid of 6 selection coefficients. See text for details.

We tested the two-step algorithm under a variety of scenarios. When we applied it to the aforementioned ‘deterministically’ simulated datasets corresponding to the best-fitting model of background selection, it always closely recovered the ground truth parameters (Fig. S4). The tiny differences between predicted and simulated diversity levels introduced by discretizing sometimes caused tiny differences between the inferred and ground-truth parameter values (see e.g., Fig. S4c), but the composite log-likelihood of the inferred parameters was always higher, indicating that the algorithm is working well. Moreover, the runtime of the CTR algorithm in step 2 was typically short, presumably because its initial conditions were close to the true maximum.

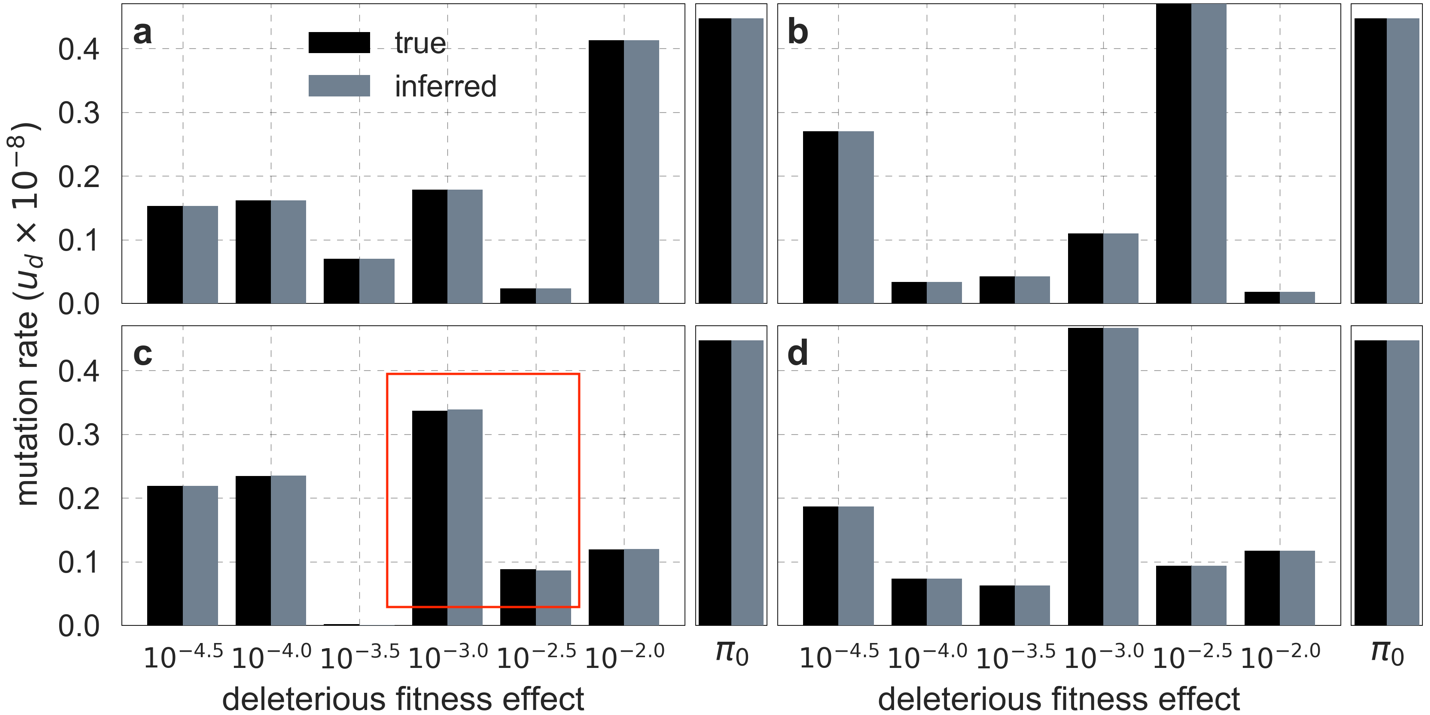

**Figure S4.** Comparison of inferred and ground-truth parameters for datasets simulated ‘deterministically’ under the best-fitting background selection model. Panels a-d correspond to different simulated datasets. Boxed region in (c) highlights the small differences between inferred and ground truth parameters introduced by discretization.

We also tested the algorithm on simulated datasets that include substantial noise in diversity levels. We generated the datasets for a sample size $n=2$ by sampling the number of pairwise differences in a bin of neutral sites from a Binomial distribution with a probability of success that equals the predicted diversity level (replacing step 3 in the simulations described above). The parameters inferred by our optimization algorithm were always similar to those used in the corresponding simulations, but with noticeable differences (Fig. S5). In all cases, however, the composite-likelihood of the inferred parameters was greater than that of the ground-truth parameters indicating that the differences were due to overfitting (which is expected given the noise we introduced in the simulations) rather than a problem in the optimization.

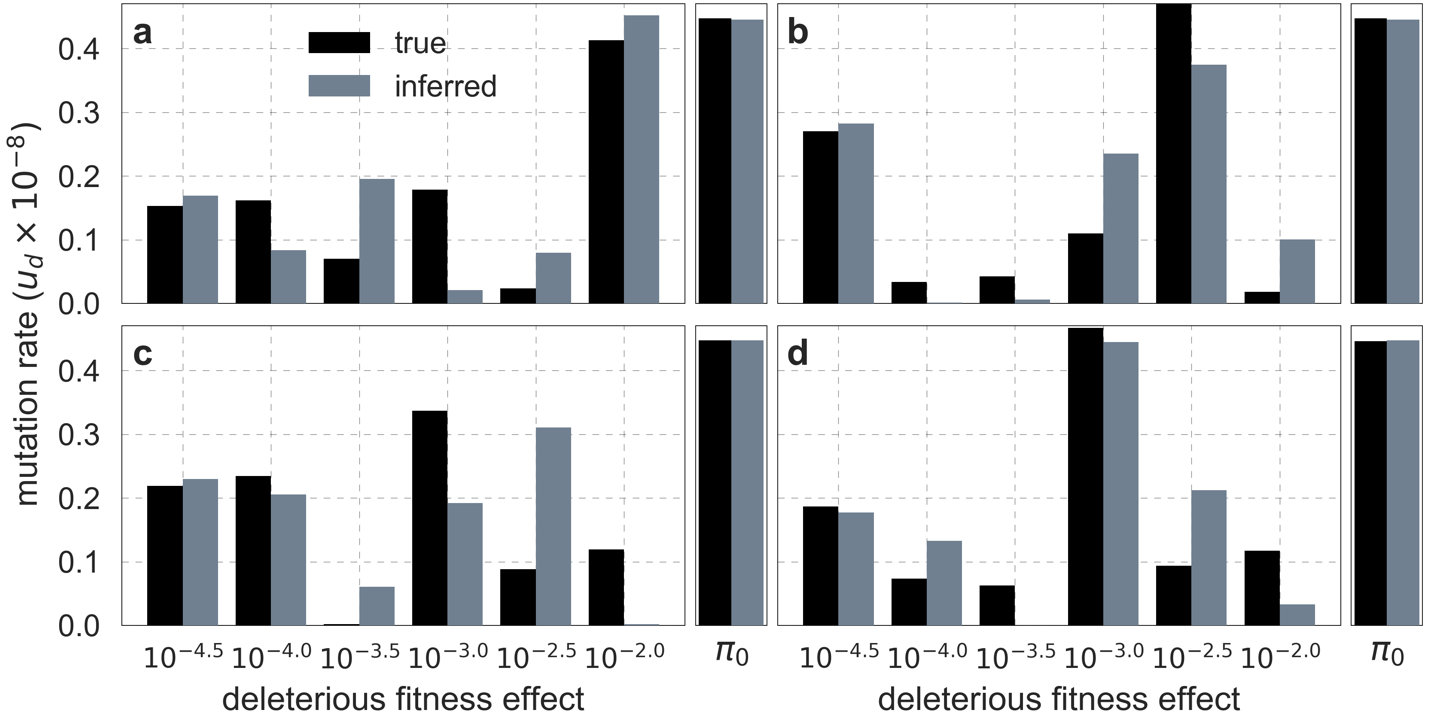

**Figure S5.** Comparison of inferred and ground-truth parameters for datasets simulated with noise under the best-fitting background selection model. Panels a-d correspond to different simulated datasets.

Lastly, we tested the optimization algorithm on datasets simulated under a joint model of background selection and selective sweeps. We modeled the effects of sweeps driven by nonsynonymous substitutions, assuming that they made up $\alpha=0.25$ of the nonsynonymous substitutions on the human lineage since divergence from the common ancestor with chimpanzees (see Section 2.7), and randomly dividing this proportion among 6 selection coefficients of by sampling from a Dirichlet distribution (with $\alpha=1$). We modeled background selection as we detailed above, and generated the dataset using the ‘noisy’ simulation scheme corresponding to a sample size of $n=2$. The parameters inferred by our optimization algorithm were always similar to those used in the simulations, with greater composite-likelihood of inferred than of ground-truth parameters indicative of overfitting (Fig. S6) as we observed in the case with background selection alone. We obtained similar results when we simulated datasets under a variety of scenarios corresponding to the combinations weak, intermediate and strong background selection ($u_{d}=5\times{10}^{-10}$, $5\times{10}^{-9}$ and $5\times{10}^{-8}$ per base per generation, respectively) with low, intermediate, and high proportions of beneficial substitutions ($\alpha=0.0125, 0.125$ and 1, respectively).

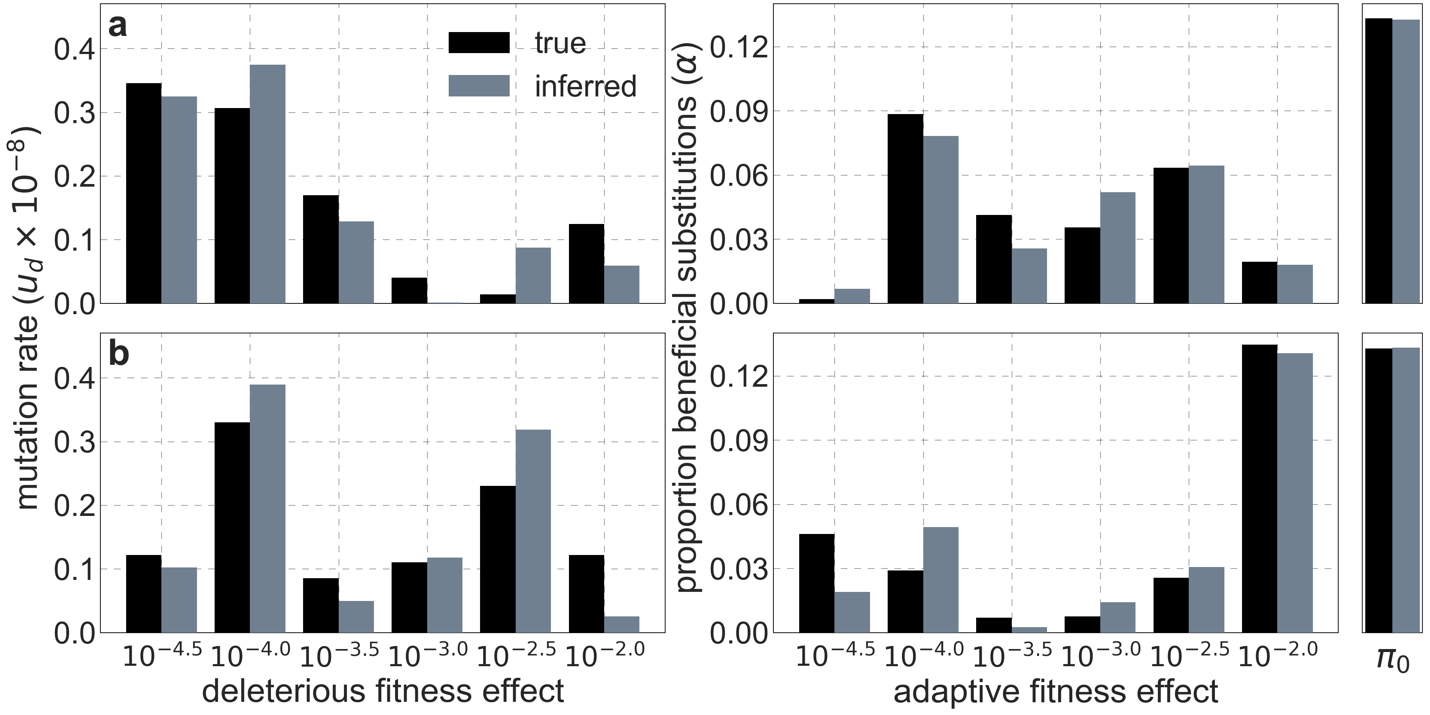

**Figure S6.** Comparison of inferred and ground-truth parameters for datasets simulated with noise under a joint model of background selection and selective sweeps. Panels a and b correspond to different simulated datasets.

#### 1.5 Thresholding

Our inference is strongly affected by forms of model misspecification that cause erroneous predictions of strong background selection effects (i.e., low values of *B*) and thus of low diversity levels at a relatively small proportion of neutral sites in our dataset. (We refer to neutral rather than putatively neutral sites for brevity and because low error in the identification of neutral sites is irrelevant to the problem at hand). These kinds of erroneous predictions can occur, for example, at neutral sites near regions that are incorrectly annotated as conserved or that are truly conserved but have proportionally fewer weakly deleterious mutations than most similarly annotated regions (because weakly deleterious mutations have strong localized effects on diversity levels). Even when neutral sites near such regions make up a small proportion of the dataset, having more of them be polymorphic than predicted can substantially reduce the composite-likelihood of models that may otherwise fit the data well (see Eq. 5), potentially biasing our inference. Here we present evidence for this problem, show how we modify our inference to solve it – by imposing a lower threshold for the value of *B* in the lookup tables or in the optimization, and address the consequences of this modification.

In Figs. S7-S9, we compare the results of our inference with and without thresholding for our best-fitting CADD-based model (the results for other models are qualitatively similar). Under the aforementioned forms of model misspecification, we might expect excess neutral diversity in regions where background selection is predicted to be strongest. Accordingly, when we apply the inference with little or no thresholding and focus on 1% of neutral sites where background selection is predicted to be the strongest, we find that observed diversity levels are up to twofold higher than our predictions (Fig. S7a). Additionally, we expect this form of model misspecification to bias the inferred distribution of selection effects toward larger selection coefficients, because smaller selection effects cause a more localized reduction in diversity levels and are therefore expected to be heavily penalized by having even relatively few misspecified regions. Accordingly, we find that the inferred distribution without thresholding is shifted toward greater selection coefficients ($t\geq{10}^{-2.5}$) compared to the distributions with thresholding (Fig. S7c(i)).

Importantly, the map of background selection effects generated without thresholding fits the data more poorly than the maps with thresholding. Notably, when we compare observed and predicted diversity levels around nonsynonymous substitutions, we find that the predictions generated without thresholding underestimate the reduction in diversity levels near nonsynonymous substitutions (inset in Fig. S7b). This can be explained by the bias toward larger selection coefficients, which causes the inference without thresholding to underestimate the reduction in diversity levels near conserved regions that are specified correctly (in order to avoid the reduction in diversity levels near misspecified regions). Additionally, when we compare the fit of maps with and without thresholding, we find that without thresholding the composite-likelihood is lower (Fig. S7c(iv)), the variance in diversity levels explained throughout the range of window sizes is lower (Figs. S7d and S8) and the calibration of our predictions is poorer (Fig. S7a; this remains the case when we exclude the top and bottom 5% of our predicted values, such that the predictions with and without thresholding span the same ranges of values; e.g., Pearson $R^{2}$ of 0.99 and 0.97 with a threshold of $B=0.6$ and without thresholding, respectively).

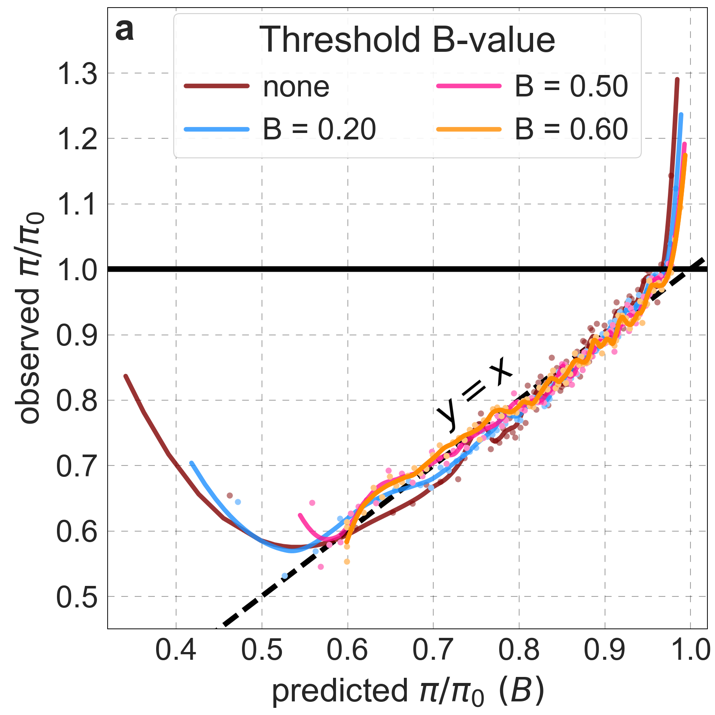

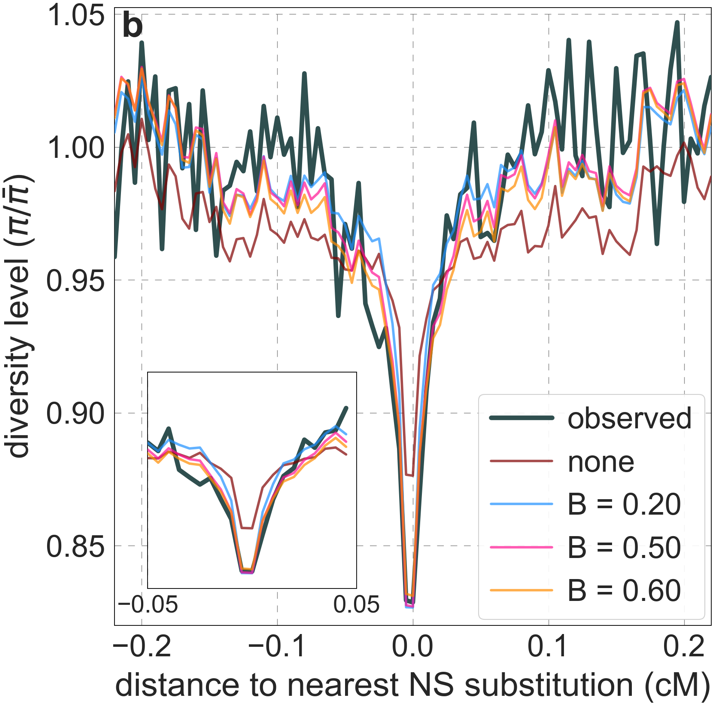

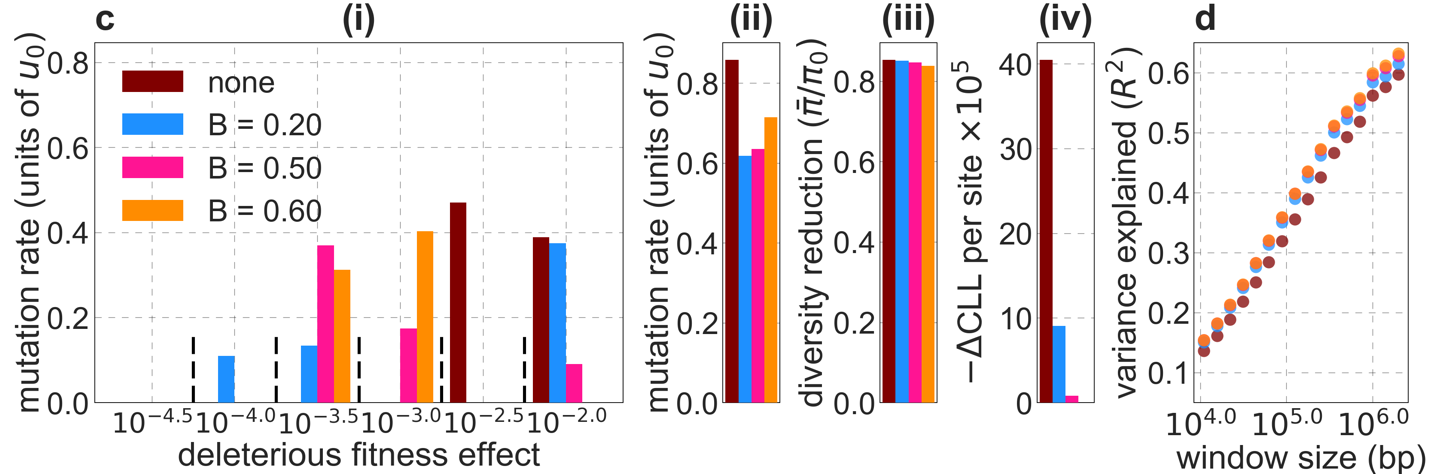

**Figure S7.** Comparison of inference results with and without thresholding. The results shown correspond to our best-fitting CADD-based model (see Main Text), with threshold values of $B=0$ (without threshold, labeled ‘none’), 0.2, 0.5 and 0.6 applied in the lookup tables. a) Observed vs. predicted neutral diversity levels across the autosomes. The graph was generated as detailed in Fig. 5. Note that the division of neutral sites among bins varies with the choices of thresholds because it is based on corresponding maps. b) Observed vs. predicted neutral diversity levels as a function of genetic distance from human-specific nonsynonymous (NS) substitutions. The graph was generated as detailed in Fig. 3, using a narrower range of genetic distances to NS substitutions to highlight differences among thresholds. c) Parameter estimates and summaries of the inferences. From left to right: i) The estimated distribution of fitness effects, described in terms of the rate of mutation per generation with a given selection coefficient. Mutation rates (throughout) are measured relative to the estimate of the total mutation rate in humans, $u_{0}=1.4\cdot{10}^{-8}$ per bp per generation (see Section 5). ii) The total deleterious mutation rate ($u_{d}$) measured in units of $u_{0}$. iii) Our prediction of the mean reduction in neutral diversity level due to background selection, measured as the ratio of the average predicted level across the genome, $\bar{\pi}$, to the predicted level in the absence of selection at linked sites, $\pi_{0}$. iv) The reduction in composite log-likelihood (CLL) per site relative to the model with the highest CLL. Differences in CLL should be interpreted with caution, as this measure does not account for linkage disequilibrium. d) The proportion of variance in diversity levels explained ($R^{2}$) on different spatial scales (measured in non-overlapping windows).

We considered two ways of thresholding, where in both we set any value of $B$ that is below the threshold to the threshold value: 1) applying the threshold in the lookup tables, i.e., *before* the composite-likelihood maximization step, and 2) applying the threshold *at each step* of the maximization, when *B* values are calculated for a given distribution of selection effects (see Eq. 6). The two approaches yield similar improvements in fit at equivalent threshold levels, and even applying a relatively low threshold improves fits markedly compared to *B-*maps without thresholding (Fig. S8). Based on our metrics of fit, we find that applying a threshold of $B=0.6$ in the lookup tables yields the best fits (Fig. S7-9), although thresholds within the range $0.45\leq B\leq0.65$ yield comparable results. Nonetheless, lower thresholds yield better fits to data in regions of the genome where selection is particularly strong (e.g., Fig. S7a-b, red box in Fig. S9). It may therefore be useful to use a lower threshold when considering regions of the genome that are subject to especially strong background selection. We provide *B*-maps for a range of thresholds that can be downloaded at github.com/sellalab/HumanLinkedSelectionMaps (in addition to the ‘best-fitting *B*-maps’ presented in the Main Text).

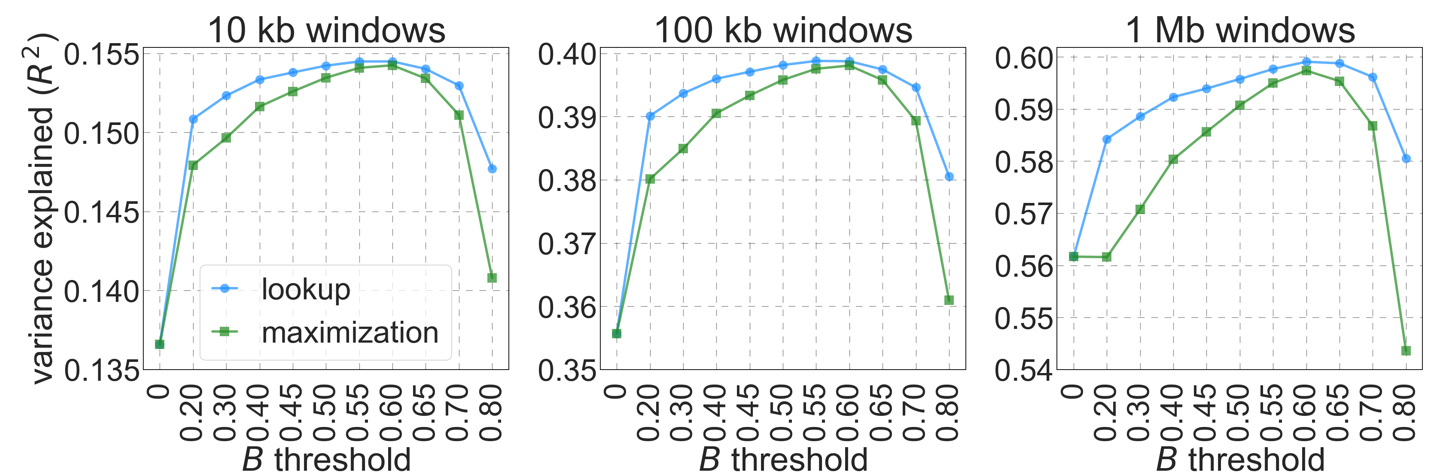

**Figure S8.** The proportion of variance explained in 10 kb, 100 kb and 1 Mb windows using a range of $B$ thresholds applied to lookup tables (‘lookup’) or during maximization (‘maximization’).

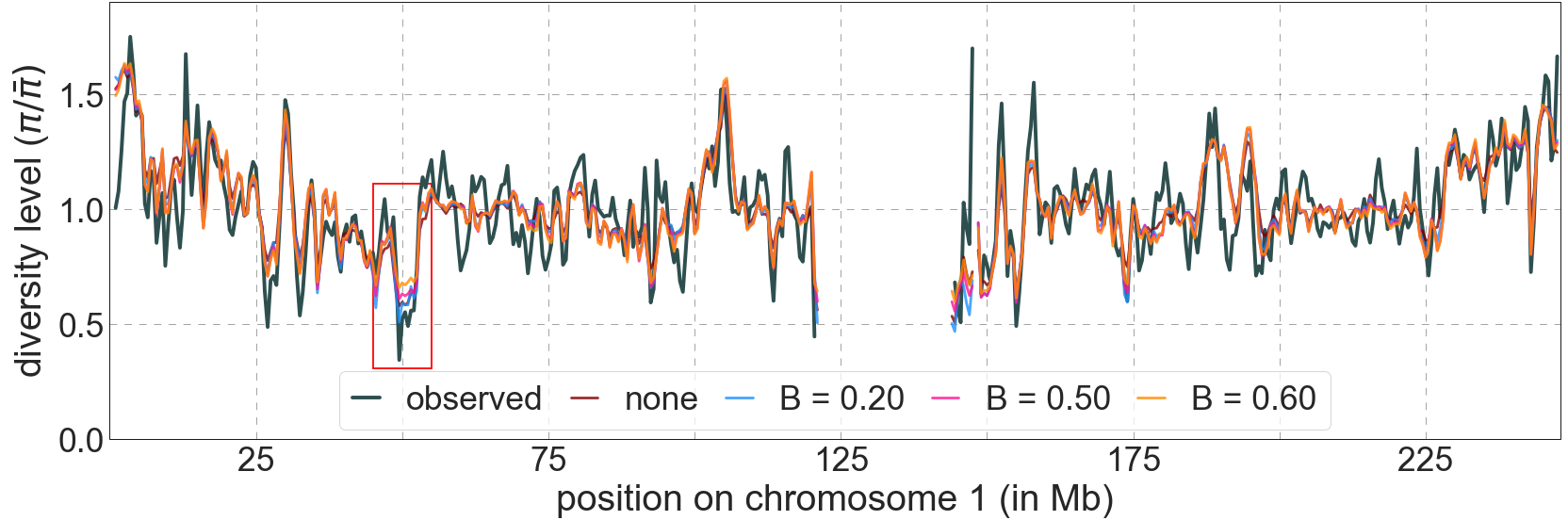

**Figure S9.** Predicted and observed diversity levels along chromosome 1 in the YRI sample. Diversity levels are measured in 1 Mb windows, with a 0.5 Mb overlap, with the autosomal mean set to 1. Thresholds were applied in the lookup tables. Lower thresholds yield better predictions in regions with low diversity levels, e.g., near 50 Mb (red box).

While thresholding largely resolves the aforementioned problem of model misspecification, it also introduces some problems. First, as we already noted, it leads to an underestimation of background selection effects at ~4% of the genome in which background selection effects is predicted to be the strongest. Second, thresholding potentially biases our estimates of the distribution of selection effects. While this bias is probably smaller than the bias without thresholding, its form and magnitude are not obvious. This is why we decided not to report the inferred distributions of selection effects in the Main Text. We are working on more principled ways of resolving the problems introduced by model misspecification, but these fall beyond the scope of the current paper.

#### 1.6 Software

We provide a set of Python programs to download and format the genomic data that we use (see Section 2), infer maps of the effects of linked selection and reproduce all of the analyses and figures described in this study (github.com/sellalab/HumanLinkedSelectionMaps). We rely on publicly available software for some steps, including the PHAST package (*17, 18*) , which we use to identify conserved regions and to estimate substitution rates (see Sections 3 and 5), and a modified version of the calc_bkgd program from McVicker et al. (*2*), which we use to generate lookup tables of the effects of background selection (see Section 1.2).

**Running the inference pipeline.** The inference pipeline is controlled by a data structure called RunStruct, which is initialized with information about input/output file paths used, model parameters and other control variables, such as the precision $\epsilon$ of lookup tables (see Section 1.2) and the *B* threshold (see Section 1.5). Once RunStruct has been initialized, the pipeline proceeds through the following steps:

1) Download and organize input files (annotations, genetic maps, etc.).
2) Create lookup tables of the effects of background selection and/or selective sweeps (Section 1.2) for the given set of selected annotations and grid of selection coefficients.
3) Organize polymorphism dataset that includes polymorphism data at putatively neutral sites (Section 2.1), corresponding estimates of substitution rates (Section 3.3) and corresponding values of lookup table into our compressed bins format (Section 1.3).
4) Run the two-step optimization algorithm to obtain estimates of model parameters, a map of the predicted effects of linked selection, and summary statistics including, e.g., the estimated deleterious mutation rate ($u_{d}$) and proportion of beneficial substitutions ($\alpha)$ associated with different annotations and the average reduction in diversity levels ($\bar{\pi}/{\pi_{0}})$.

**Parallelization and runtimes.** The composite-likelihood calculations during optimization can be partitioned into sums over subsets of bins of neutral sites, which in turn allows us to parallelize the optimization. The number of processing cores used in optimization is controlled by RunStruct. For our best-fitting models of background selection, loading lookup tables and neutral polymorphism data and running the two-step optimization requires ~1GB of memory for each of the 15 processes in step 1 and the single process in step 2. Running each process on a single core takes ~12-24 hours or ~200-400 CPU$\times$ GB hours. The computing cluster we used allows up to 12 cores per process and thus using parallelization we were able to run the optimization for the best-fitting models in 1-2 hours. Our most complex models (see Section 4) required up to 10GB of memory per process and took up to 60 hours with using 12 cores (i.e., ~${10}^{4}$ CPU$\times$ GB hours).

### 2. Data sources and filters

#### 2.1 Polymorphism data

We download 1000 Genomes Project phase 3 VCF files for all 26 populations from across the world (*19*). Unless otherwise noted, results in the Main Text and Supplementary Online Materials are based on autosomal data from Yoruba (YRI); the results for other populations are reported in Sections 7 and 9 of this Supplementary Online Materials.

We apply several filters to these data. First, we restrict our analysis to bases that pass all filters, denoted ‘P’ in the 1000 Genomes Project strictMask accessibility mask (*19, 20*). In addition, we remove low-complexity, simple repeats, duplications, and hg19 build gaps using repeatMasker files downloaded from UCSC (*21*). For each population, we restrict polymorphic sites to that population’s subset of biallelic SNPs from VCF files, excluding indels and other variants using VCFTools (*22*). Remaining sites are treated as monomorphic.

We apply additional filters to restrict our analyses to putatively neutral sites. First, we remove the union of genic regions, as detailed in section 2.4. Second, we remove all remaining sites with phastCons conservation scores greater than 0.001 as described in section 3.1. Third, we remove putatively neutral sites at the telomeric ends of autosomes, near the edges of our genetic maps (Section 2.3), as detailed in Section 3.2. Accessibility and repeat masks remove ~33.3% of all autosomal sites; excluding genic regions removes an additional ~3.3%; filtering based on phastCons scores removes another ~40.5%; and filtering sites at the telomeric ends removes ~1.2% more. We are left with a set of ~653M putatively neutral sites, which correspond to ~23% of autosomal sites (based on hg19 build).

#### 2.2 Multiple species alignment data

We rely on multiple sequence alignments to identify phylogenetically conserved and non-conserved regions of the genome, as well as for estimating local variation in neutral substitution rates (see Sections 3 and 4). To this end, we download mutation annotation format (MAF) files containing 99 vertebrate genomes aligned to the human genome (build hg19), using the Multiz software from UCSC (*23*).

#### 2.3 Genetic map

We use the Hinch et al. (*24*) genetic map, which was inferred from ancestry switches in African-Americans. At the >10 kb scale, it is highly correlated to other fine-scale maps (e.g., (*25*)), International HapMap Consortium (*25, 26*)). Among high-resolution genetic maps in humans, however, this one is likely the least confounded by diversity levels along the genome.

#### 2.4 Human gene annotations

We use genic annotations from the UCSC knownGene database (*27*) to identify putative targets of selection as well as regions that should be removed from our set of putative neutral sites. To this end, we rely on exon coordinates from knownGene transcripts to identify four kinds of annotations: 1) upstream/downstream regions, defined as 1 kb upstream of a transcript start and 1 kb downstream of a transcript end; 2) untranslated region (UTR), both 5’ and 3’; 3) protein coding sequences (CDSs); and 4) splice regions, defined as 200 bp from the start and end of each intron.

For the purpose of identifying putative targets of selection, we rely on a non-overlapping subset of these four annotations. For genes with multiple splice variants, we keep only the set of exons within the longest isoform. In rare cases of two overlapping gene predictions, we retain the gene with the longer exonic sequence. For the purpose of removing putatively functional regions from our set of putative neutral sites, however, we remove the union of all four annotations for all gene transcripts.

#### 2.5 CADD scores

We use CADD scores (*28, 29*) in order to annotate putative targets of selection in a couple of models (Sections 4.4 and 4.5). The standard CADD scores rely on the map of background selection effects generated by McVicker et al. (*2*) as one of their inputs. While this input has minor effects on CADD scores (i.e., the top 1-10% of scores; see Supplementary Table 3 (*28*)), in order to avoid any measure of circularity we approached the Kircher Lab (Martin Kircher, Lusiné Nazaretyan, Philip Rentzsch and Max Schubach), who manage the development of CADD scores, and who kindly agreed to generate and share a version of CADD score without the background selection map as input (this set of CADD scores is available on request from either the Kircher or Sella labs). For each site in the genome, we retain the highest of the three CADD scores (corresponding to the three possible point mutations). We use the distribution of scores across the autosomes to determine cutoffs for our annotations (e.g., sites within the top 6% of scores) and use sites with scores that exceed these cutoffs as putative targets of selection (sometimes in conjunction with another annotation, e.g., exons).

#### 2.6 ENCODE cCRE annotations

In two of our models (Section 4.4), we consider regulatory elements identified by the ENCODE project as putative targets of selection (*30*). To this end, we download ENCODE candidate cis-regulatory elements (cCREs) from the Tier 1a group of biosamples, which include experimental support from all relevant assays used to define elements: high DNase signal and high H3K4me3, H3K27ac or CTCF signal (*30*). The resulting cCREs are categorized as 1) enhancer-like signatures (ELS), 2) promoter-like signatures (PLS), 3) CTCF-bound (CTCF) and 4) poised elements marked by DNase and H3K4me3 (H3K4me3). cCRE annotations were downloaded for each individual Tier 1a biosample using the SCREEN tool (*30*) and lifted over from hg38 to hg19 coordinates.

#### 2.7 Substitutions in the human lineage

We rely on an estimate of the human-chimpanzee ancestor inferred using the Enredo-Pecan-Ortheus (EPO) 6-species alignment pipeline (*31*) to identify likely substitutions on the human lineage. We use subsets of these substitutions that arose in putative targets of positive selection as candidate substitutions resulting in classic sweeps (Section 4.5). We derive sets of likely substitutions in a couple of different ways. First, we compare the reconstructed ancestral genome with the human hg19 reference, taking the differences as putative substitutions. In this case and others, we do not differentiate between low and high confidence calls (lower and upper case, respectively) in the estimated ancestor. Because the hg19 reference genome is a composite of genomes with different ancestries (*32*), we also consider population-specific inferences of substitutions for YRI and CEU. To this end, we compare the reconstructed ancestral genome with the polymorphism data collected in the 1000 Genome Project for a given population. If a site is monomorphic in the population and differs from the HC ancestor, we include the site in our set of substitutions. For biallelic sites where one of the two alleles is ancestral, we randomly choose one of the alleles with probabilities that are weighted by allele frequency; if the chosen allele is the derived one, the site is considered a substitution. We generate two such samples for a given population to see whether different choices of substitutions affect our results. In practice, each of these sets differs from the set based on the hg19 reference at fewer than 1% of sites, the differences between the two samples for a given population are even smaller, and the results of our inference end up being insensitive to these differences (Section 4.6).

#### 2.8 Covariates of *B*

In Section 8, we ask whether genomic features that covary with *B* could account for the divergence between observed and predicted diversity levels in the ~2% of sites in which background selection is predicted to be the weakest. In addition to annotations of features whose sources were already mentioned, we also use the following datasets: 1) BED files of CpG islands downloaded from the UCSC Table Browser (*21*); 2) BED files of testis CpG methylation levels in downloaded from the GEO database (GEO accession: GSM1127119) (*33*); 3) coordinates of C>G hypermutable regions, given at 1 Mb resolution, taken from the Supplemental Information of Jonsson et al. (*34*); 4) coordinates of centromeres and telomeres taken from the hg19 gaps track in the UCSC Table Browser (*21*); 5) inferred proportions of archaic ancestry in European (CEU) and East-Asian (CHB/CHS) populations based on estimates from Steinrucken et al. (*35*).

### 3. Choice of exogenous parameters

Fitting our model to data requires several choices beyond those of datasets and filters. Here, we describe how we chose our set of putatively neutral sites and estimate the substitution rate at these sites. In Section 4, we describe how our results depend on the choice of targets of selection.

#### 3.1 Choosing putatively neutral sites based on phylogenetic conservation

Our main source of information for choosing the set of putatively neutral sites is the degree of conservation in multiple species alignments. To this end, we rely on running phastCons (*17*) on subsets of the 99-vertebrate alignment (from which we exclude the human genome). PhastCons fits a phylogenetic hidden Markov Model (phylo-HMM) with two states, neutral and conserved, to multiple species alignments of contiguous sites along the genome using the relative substitution rates in the alignment columns to infer conservation. The phastCons score is the posterior probability that any given site is conserved. In principle, including more species in the alignment increases the power to distinguish between conserved and neutral sites (Fig. S10a). However, as the phylogenetic distance from humans increases, sequence conservation might become less informative about conservation in humans because of functional turnover (*36, 37*). In practice, the latter effect is ameliorated by the fact that phastCons only uses information at aligned sites and the proportion of the genome that aligns to the human reference decreases with phylogenetic distance (Fig. S10b), especially in regions with considerable turnover.

In relying on phastCons scores to identify a set of putatively neutral sites, we need to choose two parameters: the phylogenetic depth of species included in the alignment and the cutoff conservation score below which a site will be considered neutral. In both cases, we pick the parameter values that maximize the variance in diversity levels explained by our best-fitting models (Fig. S11). Given these criteria, we chose to base our set of neutral sites on the alignment of supra-primates (Fig. S11a), and use the 35% of sites (in the set remaining after filters and removing genic regions; see Sections 2.1 and 2.4) with the lowest phastCons scores in this alignment, which includes sites with scores ≤ 0.001 (Fig. S11b). These choices are robust to the phylogenetic depth used to specify the selection targets (see Section 4) and to the window size in which we measure the variance explained by our model (we show the results for windows of 1 Mb in Fig. S11).

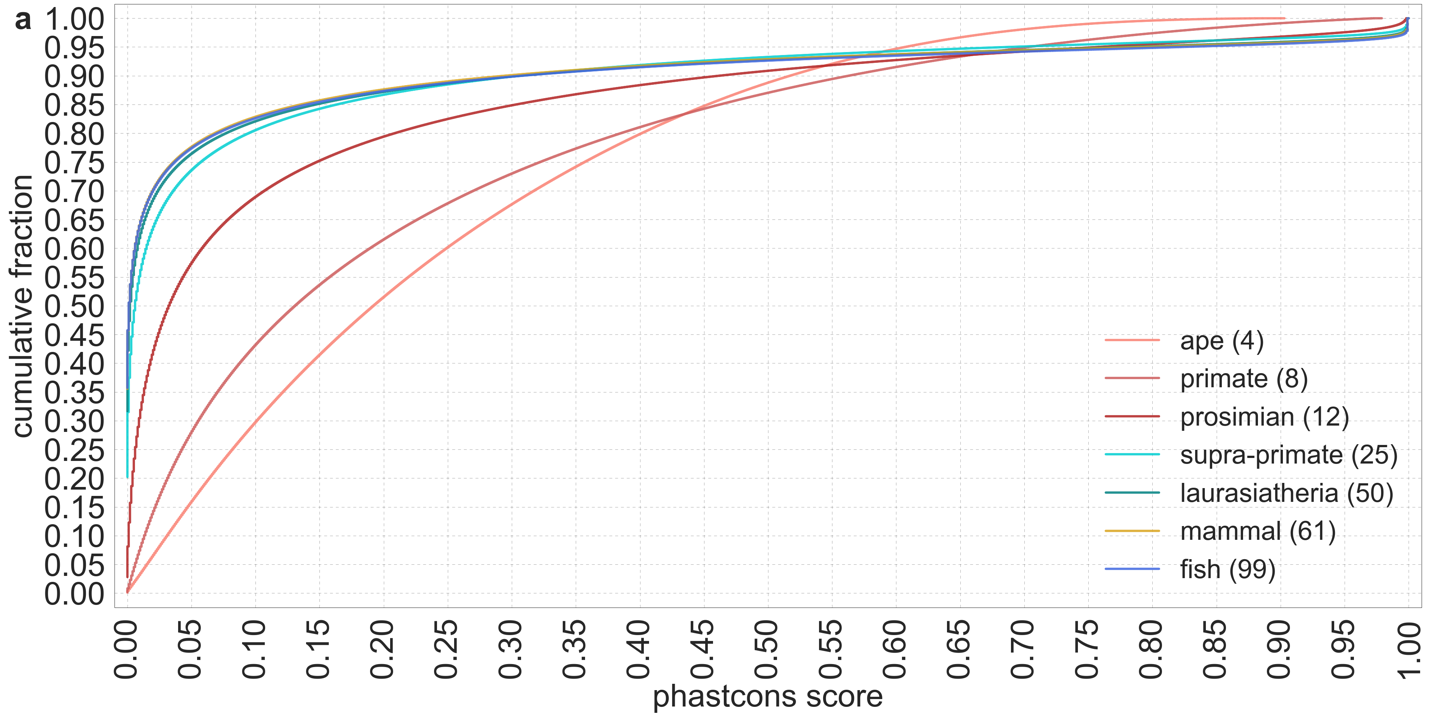

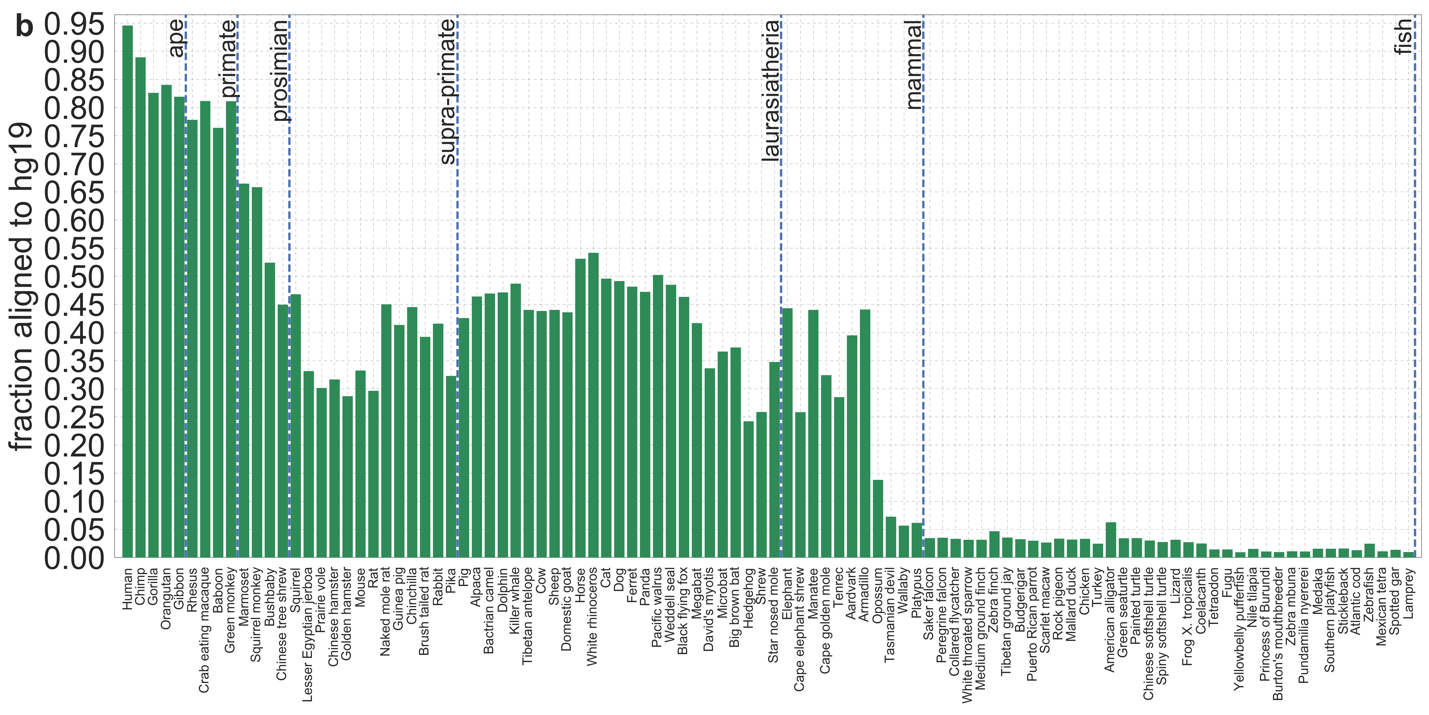

**Figure S10.** The distribution of phastCons scores across autosomes for varying phylogenetic distances from humans. a) The number of species included at each phylogenetic depth is noted in the caption. As the number of species in the alignment increases, the ability to distinguish between conserved and neutral sites increases. b) The proportion of a species’ sequenced genome that aligns to the human reference (hg19) decreases with their phylogenetic distance from humans. The decrease is not monotonic because of other factors, e.g., the quality of the sequencing. The proportion is not 1 for humans because of missing information in the reference genome.

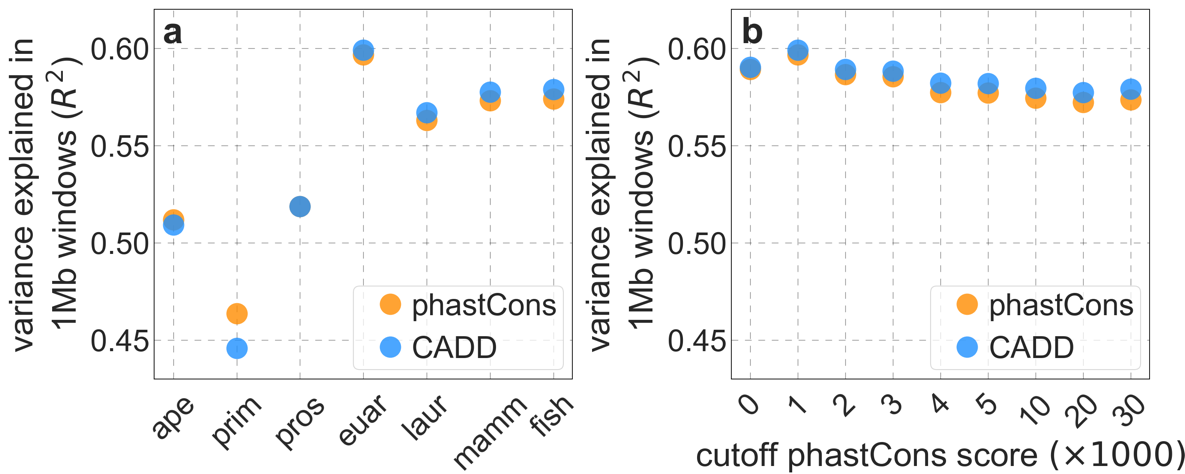

**Figure S11.** The variance in diversity levels explained by our two best-fitting models using different choices of putatively neutral sites. In (a) we vary the phylogenetic depth of the multi-species alignment (i.e., the maximal phylogenetic distance from humans to any/all of the other species) and in (b) we vary the cutoff phastCons score for the least conserved sites included in our set. The best fit corresponds to the least conserved 35% of sites (phastCons scores ≤ 0.001) in the supra-primate alignment (euar).

#### 3.2 Removing sites at the telomeric ends of chromosomes

The Hinch et al. genetic map (*24*) does not include recombination rate estimates for ~0.5-1 Mb at the 5’ and 3’ ends of autosomes. Consequently, we are unable to describe background selection effects of putatively selected regions that lie in these telomeric regions, and our inferences and predictions at putatively neutral sites near the telomeres are less accurate. We therefore exclude putatively neutral sites in telomeric regions not covered by the genetic map. Similar to our approach in the previous section, we choose the map size of the region to remove based on how the choice affects the model fit to diversity levels across autosomes (Fig. S12a). We find that filtering putatively neutral sites in 0.1 cM from the edge of the genetic map, which amounts to ~0.8% of neutral sites, largely removes this ‘edge effect’. This genetic distance makes sense, as it is roughly one at which background selection effects of deleterious mutations with $s={10}^{-3}$ – the strongest selection effects inferred to contribute substantially (Fig. S12b) – become negligible. Moreover, our estimates of model parameters are fairly insensitive to the removal of larger regions (Fig. S12b).

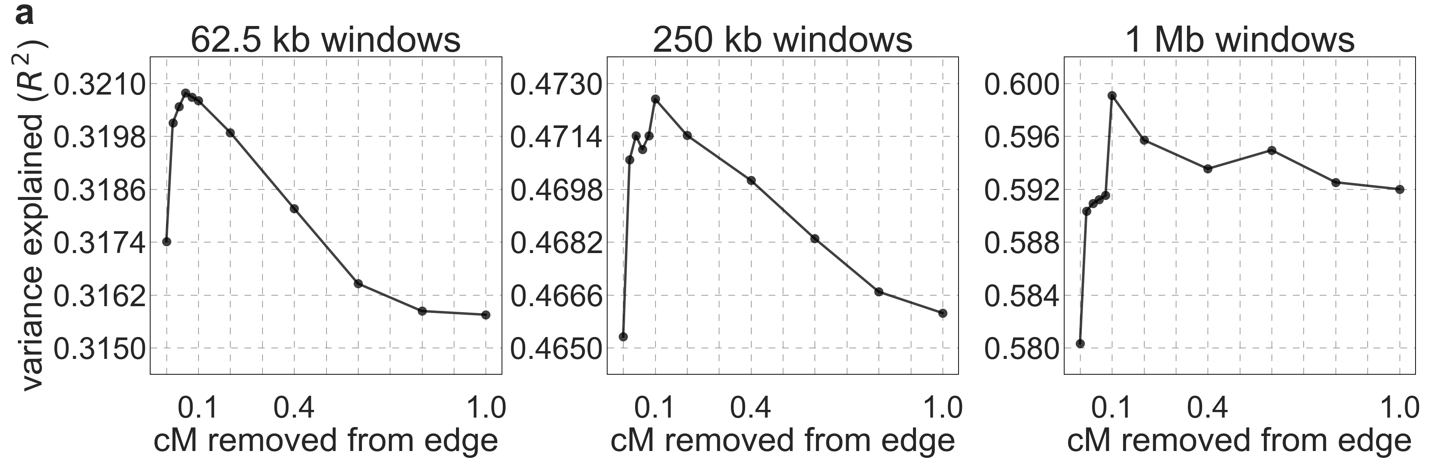

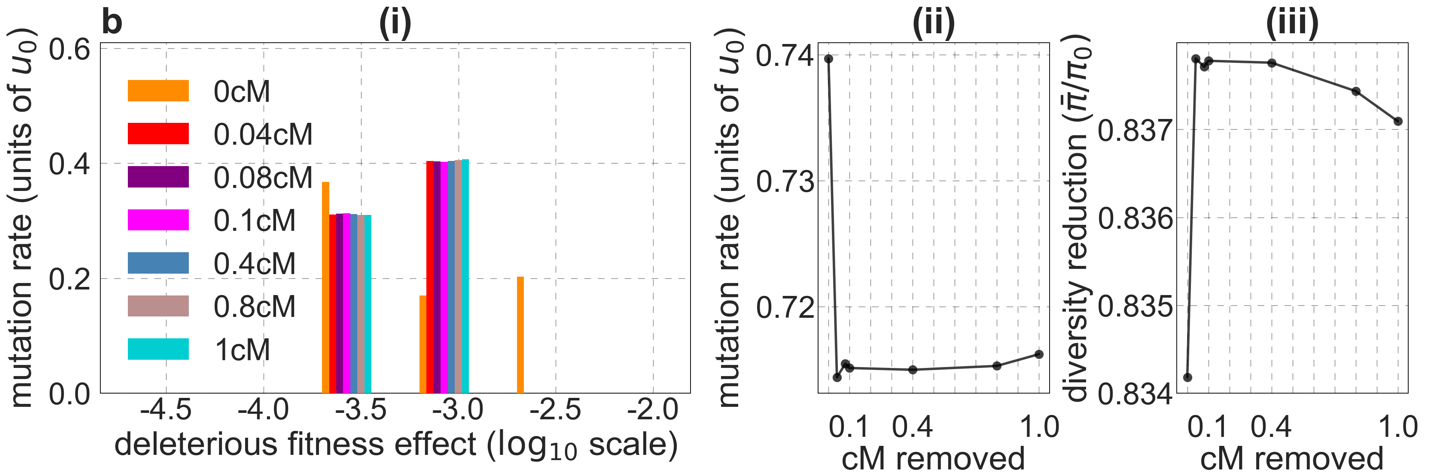

**Figure S12.** The effect of removing putatively neutral sites near telomeres on model fit and parameter estimates. We show the result for our best-fitting CADD-based model; results for phastCons scores are highly similar (not shown). a) The proportion of variance in diversity levels explained for different window sizes, as a function of the size of the removed region (in cM). b) (i-iii) Estimates of model parameters as a function of the size of the removed region (in cM).

#### 3.3 Estimating local variation in mutation rates

We rely on estimates of substitution rates at putatively neutral sites along the genome to control for the effect of variation in mutation rates on neutral diversity levels (see Eq. 1 in Section 1.1). To this end, we use phyloFit (*18*) to estimate the substitution rate in a phylogeny, in windows of putatively neutral sites across the genome. We choose the species to include in the phylogeny based on the following considerations. The number of substitutions in a given window can be approximated by a Poisson random variable with expectation $\lambda$, which is proportional to the total branch length of the phylogeny, $T$, and the number of putatively neutral sites in the window, $n$. Consequently, the precision of our estimates of the relative mutation rate increase with $\lambda\propto n\cdot T$. Including more species in the phylogeny increase $T$ but reduces $n$, because it reduces the fraction of putatively neutral sites that align to the human reference in all the species included. Fig. S13a shows the trade-off between the two factors, for all subsets of 9 primate species included in the 99-vertebrate alignment (see Section 2.2). We chose the subset that maximizes $n\cdot T$, which includes 8 of the 9 species (gibbon is removed) with an average of ~0.135 substitutions per putatively neutral site.

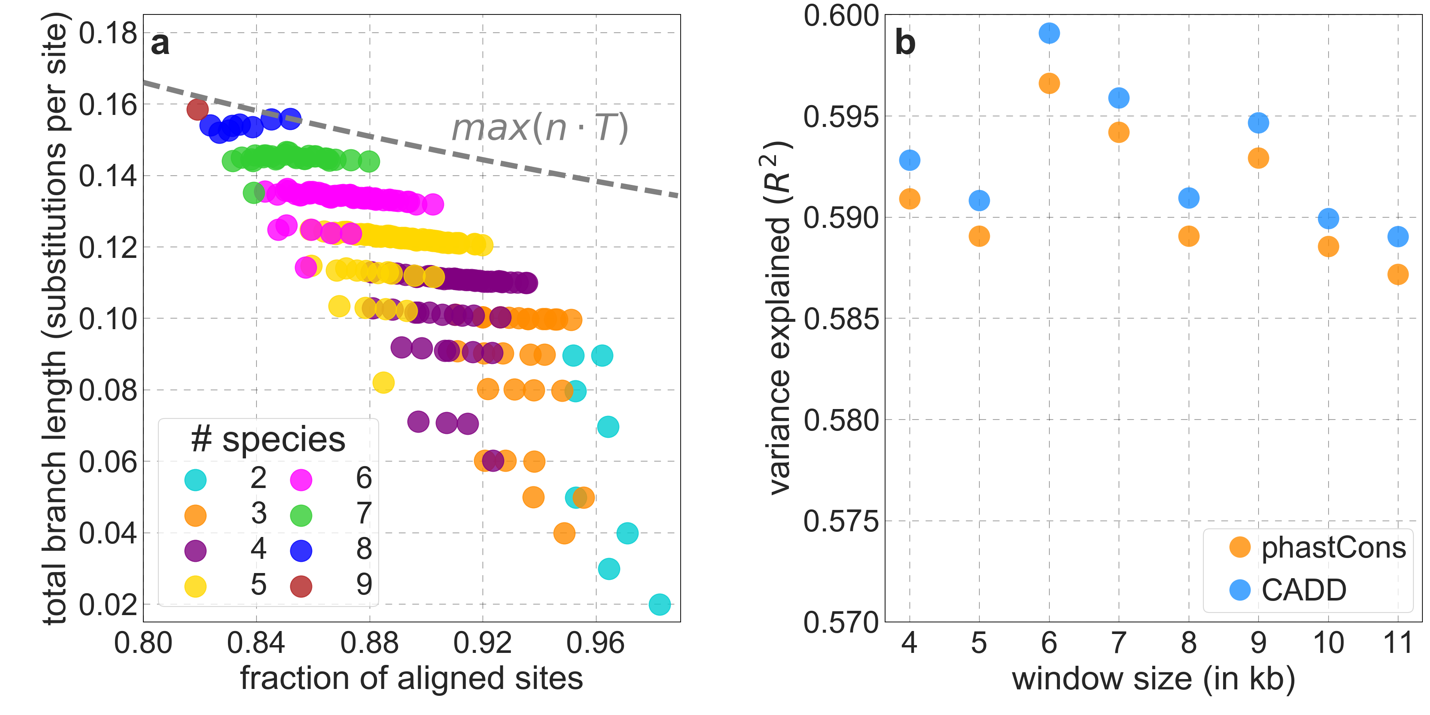

**Figure S13.** Choosing the parameters used in estimating the relative mutation rate at putatively neutral sites. a) The trade-off between the fraction of aligned sites and total branch length for subsets of the primate phylogeny. The fraction of aligned sites is estimated for our set of putatively neutral sites, and the total branch length is measured in terms of the average number of substitutions per site on the phylogeny, estimated by phyloFit. The maximum product of the fraction and branch length is attained by including all primates included in the 99-vertebrate alignment other than gibbon. b) The variance in diversity levels explained by our best-fitting models across 1 Mb windows, for different choices of window sizes (i.e., the number of putatively neutral sites) used to control for variation in mutation rates at putatively neutral sites.

We estimate relative mutation rates along the genome based on the estimated substitution rates in the 8-primate phylogeny in windows with a fixed number of contiguous putatively neutral sites. Using windows with a greater number of sites decreases the sampling error but reduces the spatial resolution of our estimates. We use the variance in diversity levels explained by our best-fitting models as a criterion for choosing the window size, finding that a window with 6000 putatively neutral sites performs best among the options we examined (Fig. S13b). This choice corresponds to mean physical window sizes of 26,454 bp (with a S.D. of 18,455 bp) and to a mean relative error of ~3.3% in our estimates of the relative mutation rate per window. We also examined other ways of estimating the relative mutation rate, including using windows of fixed physical length and sliding windows with varying degrees of overlap, but none of these approaches yielded better results.

In the analyses in which we bin neutral sites, either by their distance to genomic elements (e.g., Fig. 3) or by predicted $B$ (e.g., Fig. 5), we estimate the relative mutation rate in each bin. To this end, we use phyloFit (*18*) to estimate the substitution rate in the 8-primate phylogeny on all sites in that bin jointly and then normalize this estimate by the average across bins.

### 4. Fitting models with different targets of selection

Our framework allows us to fit models of background selection, selective sweeps, or both, based on different choices of putative targets of negative and/or positive selection. Here we detail the analysis of the models and choices that are described in the Main Text. We use several criteria to evaluate how well the models fit the data; these indicate that models of background selection alone in which the targets of selection are chosen based on constrained elements annotated by either phastCons or CADD scores are best supported by the data. We also compare the predictions of these models with those of McVicker et al. (*2*).

#### 4.1. Background selection model based on phylogenetic conservation

We first consider a model of background selection in which targets of selection are chosen based on phylogenetic conservation. We identify conserved genomic elements using phastCons scores (*17*) calculated on monophyletic subsets of the 99-vertebrate alignment to the human genome (*23*), all of which exclude the human genome itself (see Section 2.2). We vary the phylogenetic depth of the subset of species considered (i.e., the maximal distance from humans). For a given depth, we obtain targets of selection by specifying a proportion of selected sites (i.e., of the total autosomal length in hg19) and choosing those sites that have the highest phastCons scores in the alignment (after excluding some sites, e.g., from up to 5% of the four-ape alignment to less than 0.1% of the 99-vertebrate alignment, that are in our putatively neutral set). As we have done for previous choices (e.g., Section 3.1), we examine how our choices of phylogenetic depth and of proportion of selected sites affect the models’ fit to autosomal diversity levels.

We find the fit to be largely insensitive to the choice of phylogenetic depth, with models based on conservation in the full 99-vertebrate alignment fitting slightly better than other choices of depth (Fig. S14). Notably, the explained variance in diversity levels (in windows of different sizes) is similar across depths, increasing slightly with the number of species included, other than for the four-ape phylogeny (Fig. S14b and c). The fits of predicted diversity levels along the genome (e.g., Fig. S14d) and around genomic features (e.g., Fig. S14e) are similar, with none of the choices of depth clearly outperforming others. Moreover, for all choices, the predicted diversity levels are well calibrated (Fig. S14f), with the exception of regions in which background selection is predicted to be very weak, i.e., $B\approx1$ (see Section 8). When we restrict each annotation to the top 6% of scores in sites for which all phylogenetic depths include phastCons scores (~98% of sites satisfy this criterion), our results are unchanged.

Distantly related species, such as those added when we move from supra-primates (n=25) to vertebrates out to lamprey (n=99), have little effect on phastCons scores and thus on our models, because only a small proportion of their genomes align with humans (Fig. S10b). This can be seen in the high correlations between the number of conserved sites based on different depths across windows of different sizes (Fig. S15a). The spatial distribution of conserved sites is even fairly insensitive to varying the species included from four apes to 99 vertebrates (Fig. S15a). Interestingly, we later show that the improvement in fit across 1 Mb windows of the model based on conservation in 99 vertebrates compared with models based on conservation in shallower phylogenies is statistically significant, except for the model based on four-apes (Fig. S33), whereas the spatial distributions of conserved sites in the 99-vertebrate and four-ape models are the least correlated (Fig. S15). The v-shaped dependence on phylogenetic depth may reflect a tradeoff in which phastCons scores based on deeper alignments have greater power to identify long-lived selected regions (see, e.g., Fig. S10a), whereas those based on apes are better at identifying regions that are selected in humans but exhibited functional turnover in the deeper phylogeny (*36*) (see also Section 6.2).

The model fit is also fairly insensitive to the cutoff conservation score used in choosing selection targets, although choosing 5-7% of autosomal sites as targets does appear to yield slightly better fits than other choices (Fig. S16). Notably, the variance explained for different window sizes is maximized between 5-7% (Fig. S16b and c); at the higher end of the range of cutoffs from 2-9%, the fits of diversity levels along the genome (e.g., Fig. S16d) and around genomic features (e.g., Fig. S16e) appear to be slightly worse, and the stratification of observed values by predicted ones spans a smaller range (Fig. S16f). Among comparisons between models based on 6% and all other cutoffs in the range of 2-9%, only 8 and 9% lead to a statistically significant reduction of fit in windows of 1 Mb (Fig. S33). Based on these analyses, we use the model with the 6% of autosomal sites with the highest phastCons scores based on the 99-vertebrate alignment in many of our analyses, and refer to this as our *best-fitting phastCons-based model* in both the Main Text and throughout the Supplementary Online Materials.

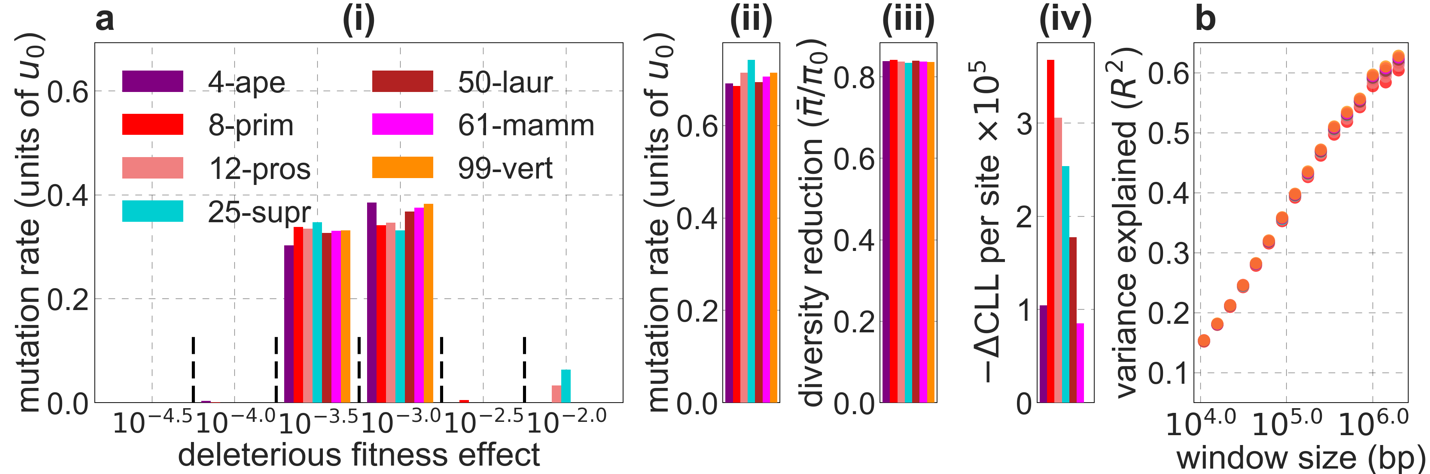

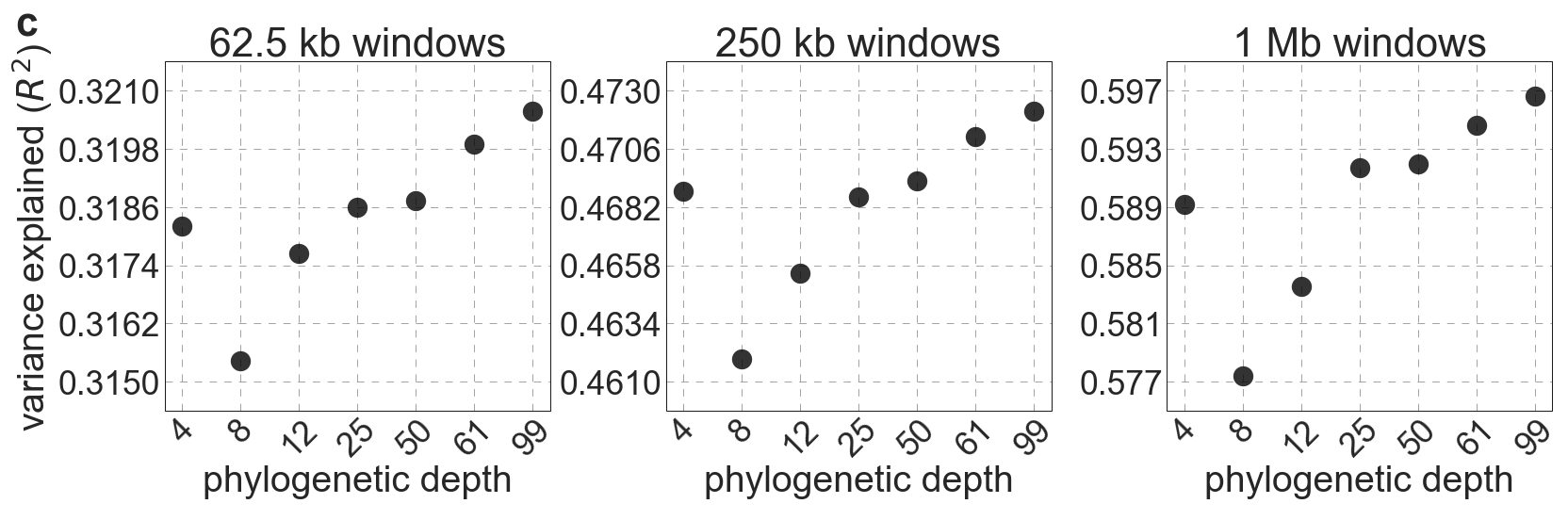

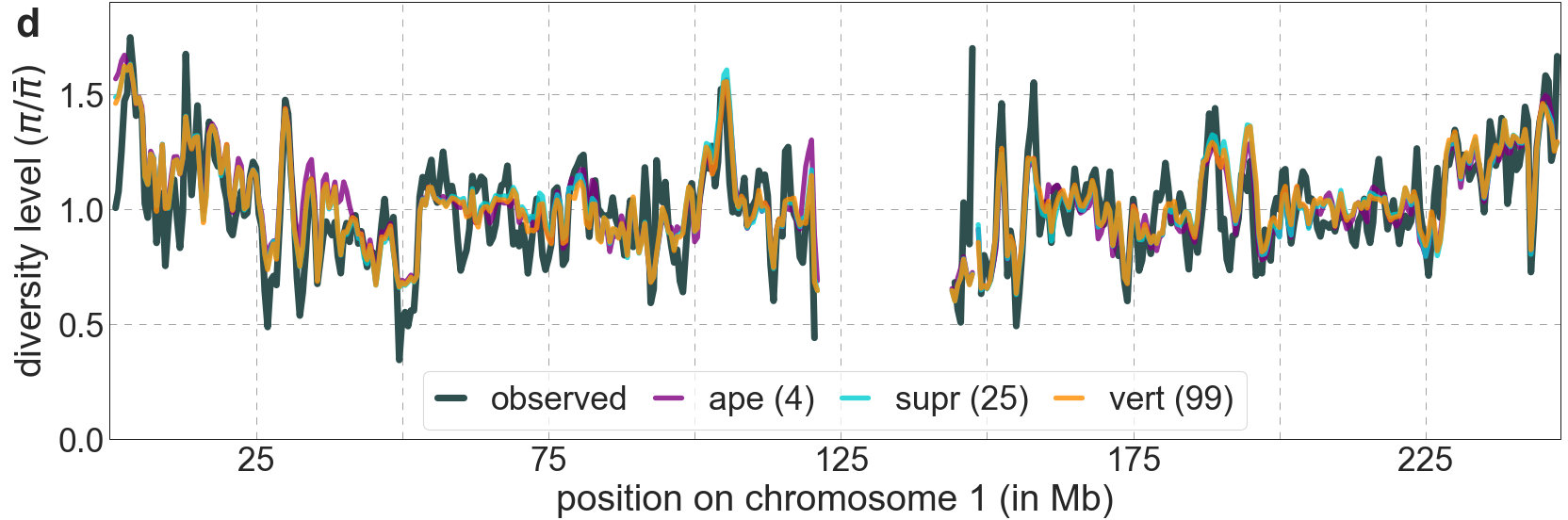

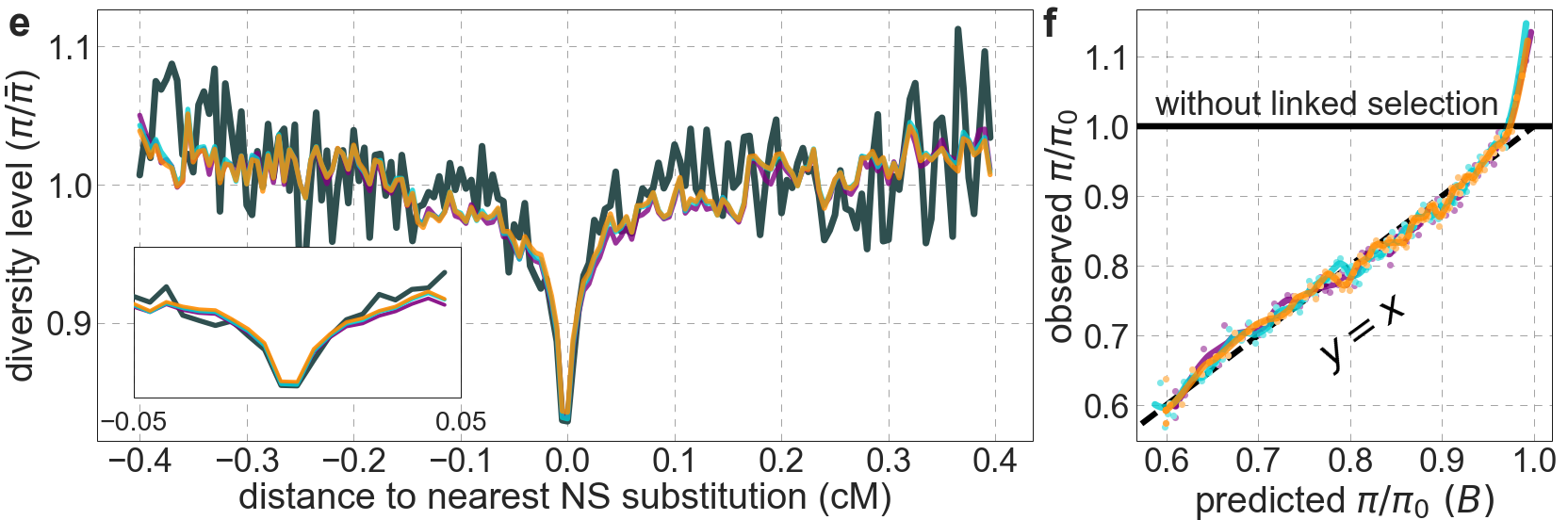

**Figure S14.** Comparison of background selection models based on phastCons conservation scores in phylogenies of difference depths. Shown are results of models based on conservation in four apes, eight primates, 12 prosimians, 25 supra-primates, 50 laurasiatherians, 61 mammals and 99 vertebrates extending out to lamprey. In all cases, we take the 6% of autosomal sites with the highest phastCons scores (excluding putatively neutral sites) as our targets of selection. Throughout the Supplementary Online Materials, with the exception of Section 7, we show results using data from YRI. ***The panels describe*:** a) Parameters and summaries of models (from left to right): i) Estimated distribution of fitness effects, described in terms of the rate of mutations with given selection coefficients. Mutation rates throughout are measured relative to the estimate of the estimated average mutation rate per bp per generation in humans, $u_{0}=1.4\cdot{10}^{-8}$ (see Section 5). As detailed in Section 1.5, the inferred distribution of selection coefficients should be interpreted with caution. ii) Estimated total deleterious mutation rate per selected site ($u_{d}$) measured in units of $u_{0}$. iii) Estimated autosomal average fold-reduction in neutral diversity levels due to selection at linked sites, i.e., the ratio of average predicted heterozygosity, $\bar{\pi}$, to average predicted heterozygosity in the absence of selection at linked sites, $\pi_{0}$. iv) The reduction in composite log-likelihood (CLL) per site relative to the model with the highest CLL. Differences in CLL should be interpreted with caution, as diversity levels at putatively neutral sites are not independent. b) The proportion of variance in diversity levels explained ($R^{2}$) on different spatial scales (measured in non-overlapping contiguous windows). c) Close-up on the variance explained for several window sizes. d) Predicted and observed diversity levels along chromosome 1. Diversity levels are measured in 1 Mb windows, with 0.5 Mb overlap, and are normalized by the mean level (as detailed in Fig. 2). The results here and in subsequent panels are shown for a subset of depths, including four apes, 25 supra-primates and 99 vertebrates. e) Predicted and observed diversity levels as a function of genetic distance to the nearest human-specific nonsynonymous (NS) substitutions. The plot was generated as detailed in Fig. 3. Inset shows closeup between -0.05 and 0.05 cM. f) Observed vs. predicted neutral diversity levels across the autosomes. The plot was generated as detailed in Fig. 5.

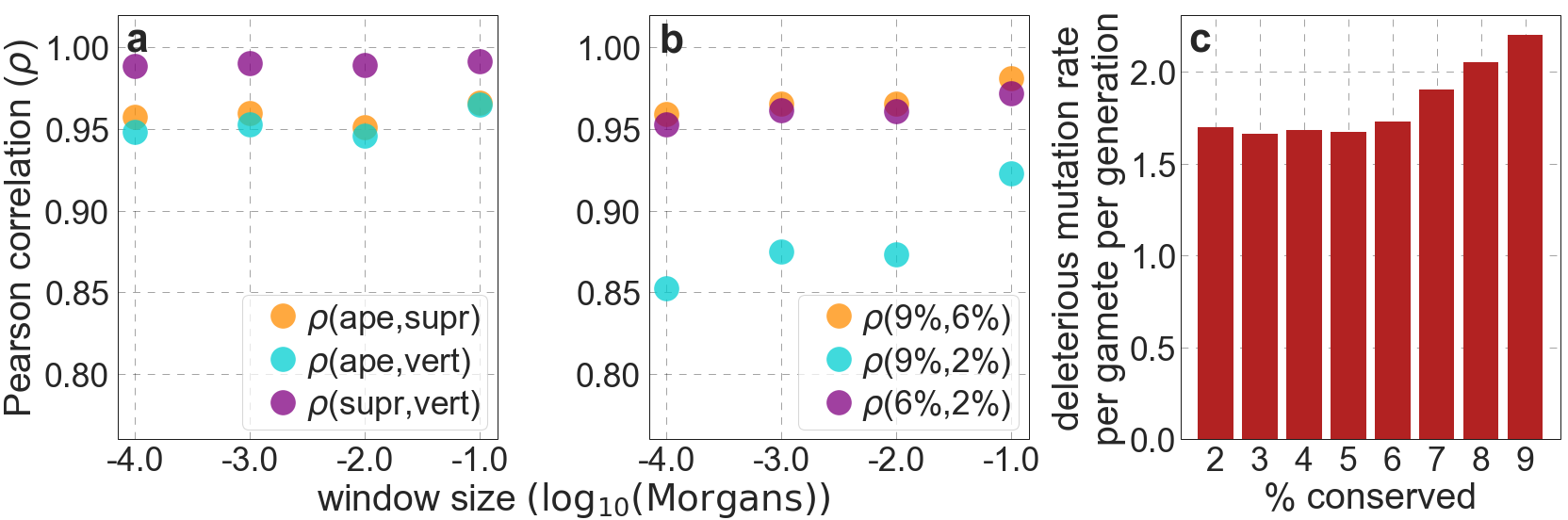
**Figure S15.** The spatial distribution of putatively selected sites targets remains similar when we vary the phylogenetic depth of the alignment used to infer conservation (shown in a), and the proportion of sites with the highest conservation scores included (in b). We compare two choices of selection targets at a time, and show the Pearson correlations ($\rho$) between the numbers of putatively selected sites among windows of different genetic lengths (measured in Morgans). The range of window sizes roughly corresponds to the spatial scales over which selection affects linked neutral diversity for the estimated range of selection effects. When we vary the phylogenetic depth, we use the 6% of autosomal sites with the highest phastCons scores, and when we vary the conservation cutoff, we use phastCons scores based on the 99-vertebrate alignment. c) The deleterious mutation rate per gamete per generation inferred as a function of assumed proportion of selected sites in autosomes.

The insensitivity of our fits to varying the conservation cutoff can be understood as follows. phastCons estimates the probability that runs of sites belong to conserved segments (*17*). When we reduce the conservation cutoff, shorter segments with high scores tend to expand to include adjacent, lower scoring sites. This results in a high spatial correlation between the conserved sites corresponding to different cutoffs (Fig. S15b). Given a lower conservation cutoff and longer ‘selected’ segments, we infer a lower deleterious mutation rate per site (Fig. S16a(ii)) but a similar deleterious mutation rate per segment (see, e.g., Fig. S15c), thereby producing similar troughs in diversity around such segments and similar fits overall.

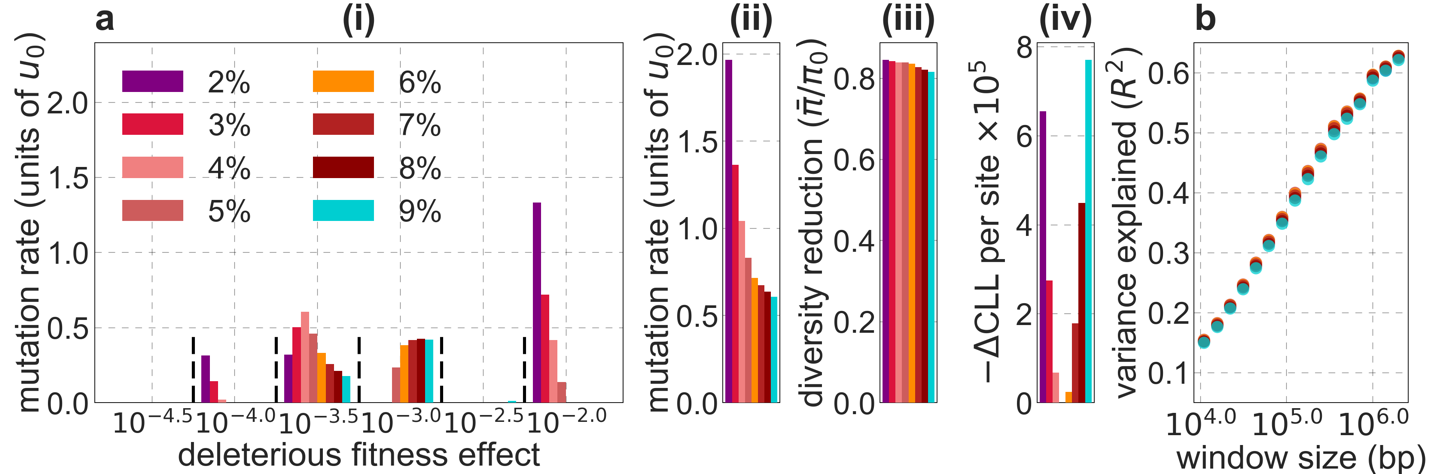

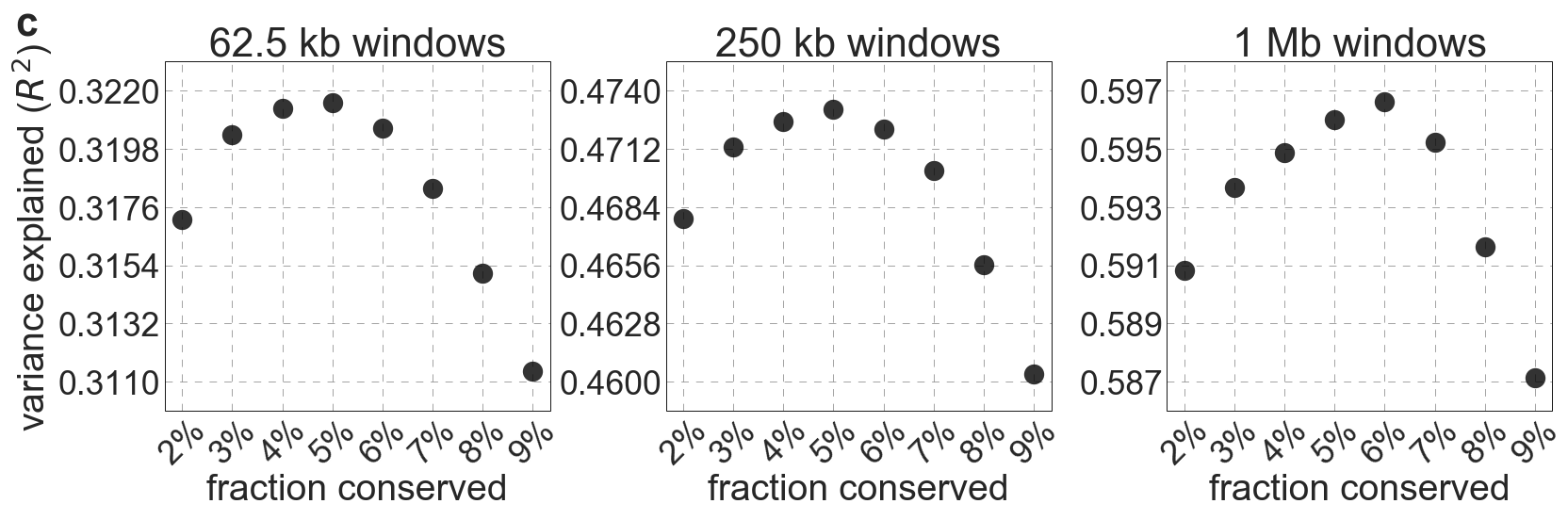

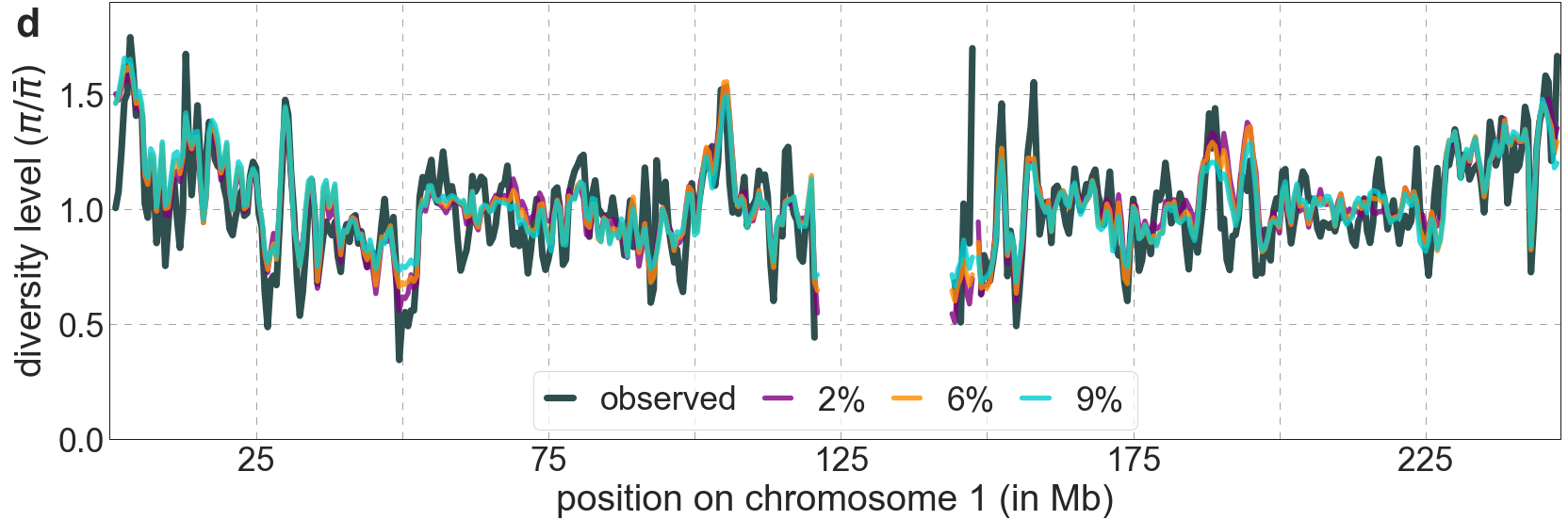

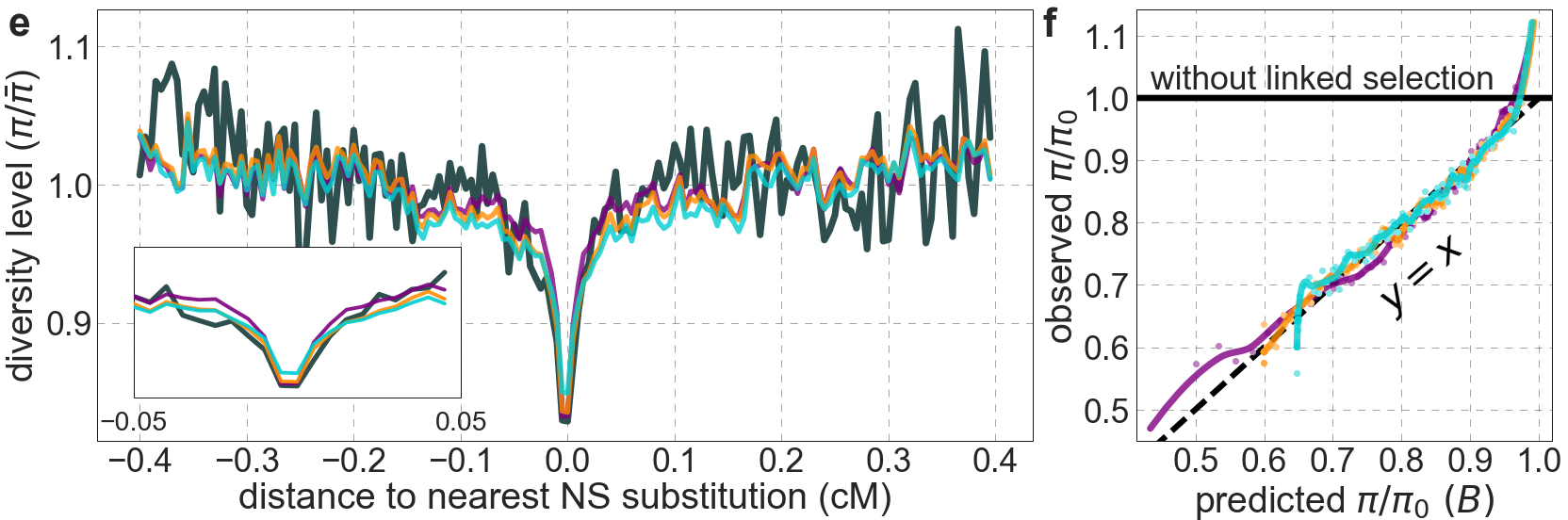

**Figure S16.** Comparison of background selection models based on phastCons scores using different proportions of autosomal sites as selection targets. In all cases considered, we rely on conservation in 99 vertebrates. Otherwise, all panels are as described in Fig. S14.

#### 4.2 Background selection model based on genic annotations

Next, we consider a model of background selection in which selection targets are chosen based on simple genic annotations, i.e., the exons divided into UTRs and protein coding sequences (CDSs), as well as regions in the immediate vicinity of these sequences controlling transcript regulation: regions 1 kb up- and downstream of transcript start/end, and splice regions 200 bp at the start and end of introns (*38, 39*) (see Section 2.4 for details). We allow selection parameters to vary among annotations, but find that in the best-fitting model only protein coding and splice regions have non-negligible deleterious mutation rates (for other annotations, ${u_{d}}/{u_{0}}<{10}^{-6}$).

We also find that this model fits much worse than our best-fitting phastCons-based model (Fig. S17): the variance in diversity levels it explains is substantially lower across different window sizes (Fig. S17b), its fit to diversity levels along the genome is discernably worse (e.g., Fig. S17c), and when observed diversity levels are stratified by the model’s predictions, they are less calibrated (Fig. S17e). The genic model does do reasonably well at predicting how diversity levels drop with genetic distance around nonsynonymous substitutions (e.g., Fig. S17d). The generally poorer fit as well as the reasonably good fit around nonsynonymous substitutions can be understood in terms of the overlap between our simple genic annotations and direct measures of constraint (Fig. S18). Namely, the genic annotations miss most constrained sites, which are intronic or intergenic (Fig. S18b), but most protein coding regions (CDSs) are constrained (Fig. S18a) explaining why models including them as an annotation perform well near them.

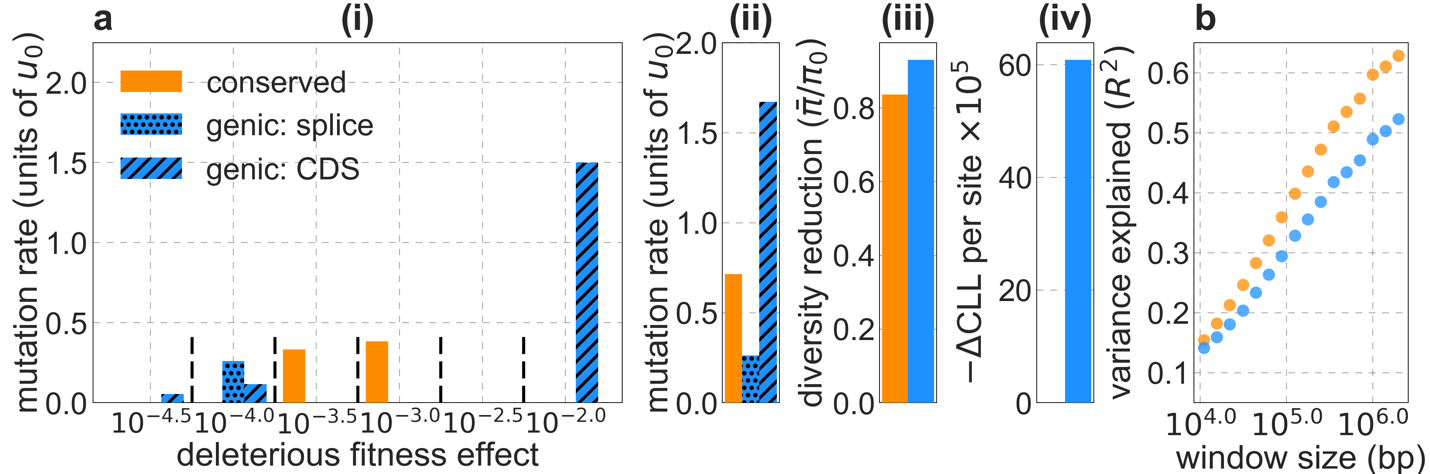

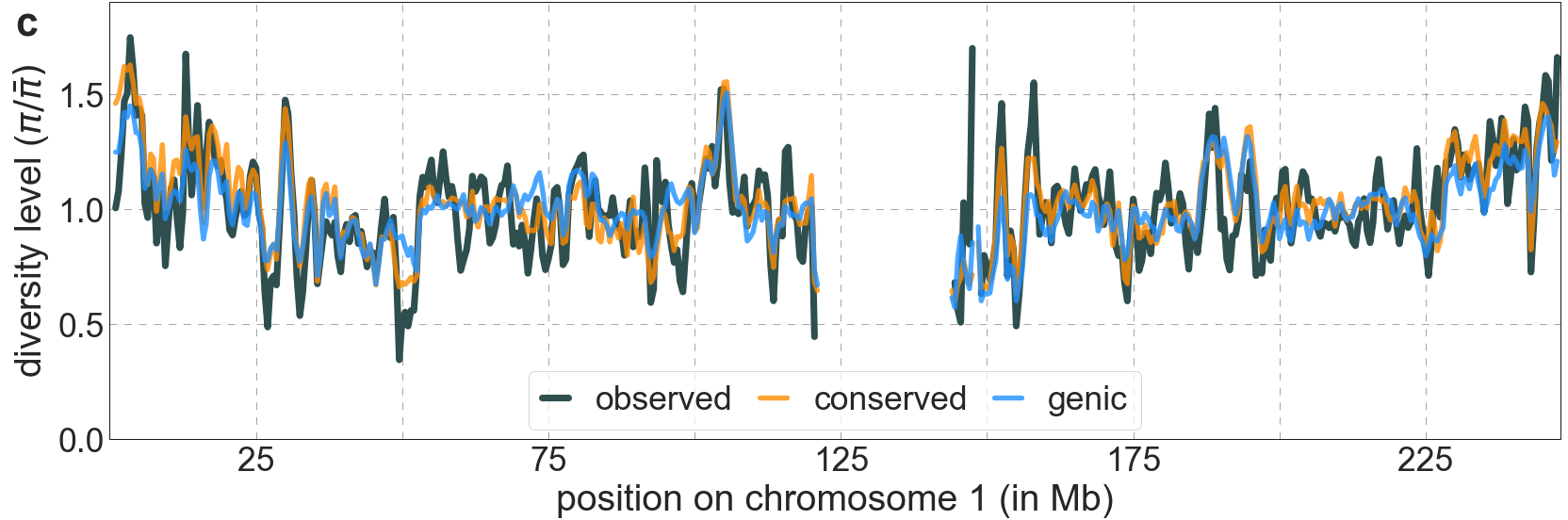

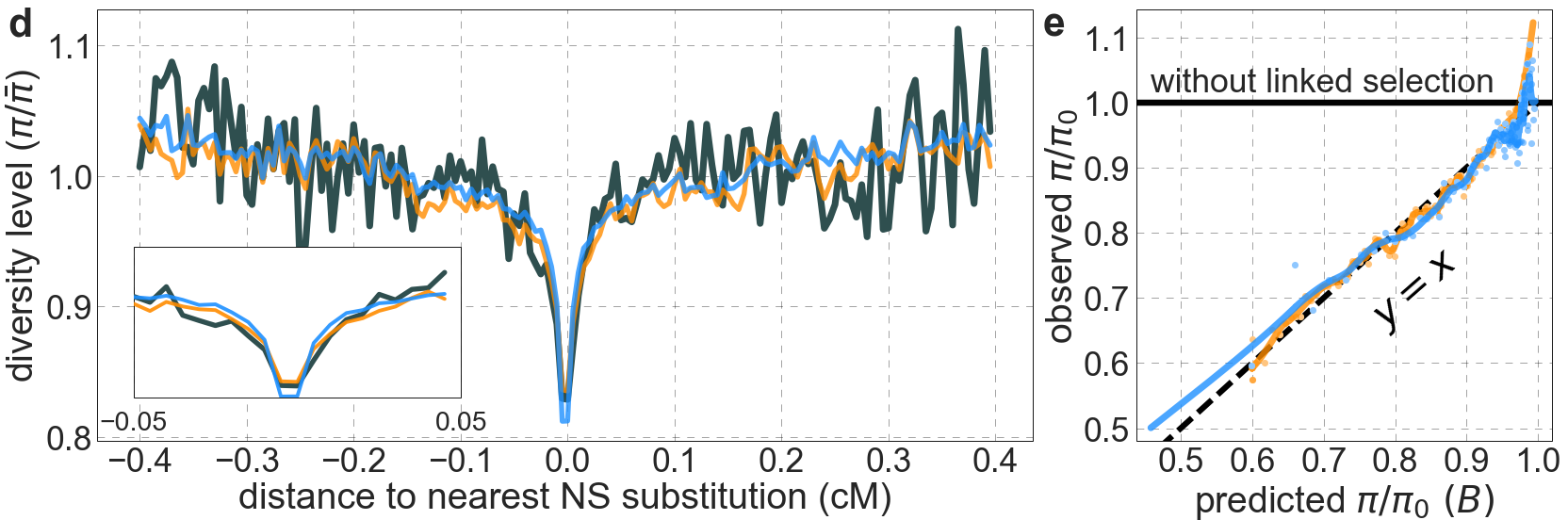

**Figure S17.** The background selection model based on simple genic annotations fits worse than our best-fitting phastCons-based model. All the panels are as described in Fig. S14 (but with the hatch-marked blue bars in a(i) and (ii) corresponding to different annotations of the genic model).

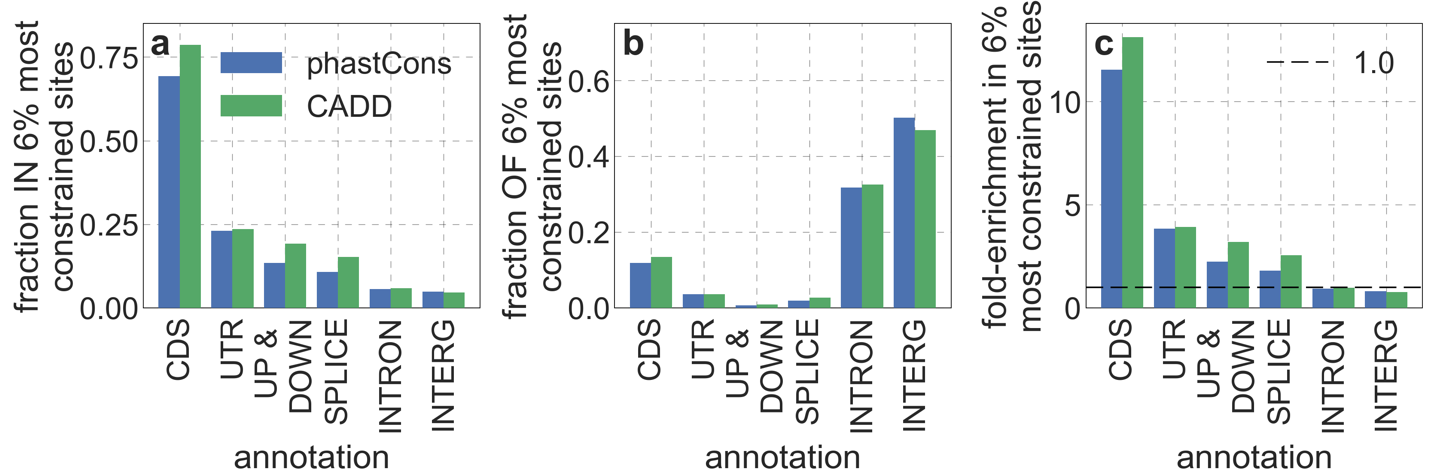

**Figure S18.** The relationship between simple genic annotations and our main measures of constraint. Specifically, we examine the overlap of the 6% of autosomal sites with the highest phastCons or CADD scores with the genic annotation detailed in the text; we added intronic (INTRON) and intergenic (INTERG) annotations for completeness. a) The fraction of each genic annotation within the 6% most constrained sites. b) The fraction of the 6% most constrained within each genic annotation. c) Enrichment of genic annotations in the 6% most constrained sites, i.e., the ratio of their proportion among constrained and all autosomal sites.

#### 4.3 Background selection models separating conserved exonic and non-exonic sites

While background selection models based on simple genic annotations do worse than those based on phylogenetic conservation, using such annotations in conjunction with conservation could allow for improved fits. Notably, it is often argued that purifying selection in protein coding regions is stronger than in functional non-coding regions (*36, 40*); if this were true, then allowing them to have different selection parameters could result in better fits. To examine this possibility, we fit a model with two types of selection target: exonic (i.e., segments combining CDSs and UTRs) and non-exonic conserved sites (see details in Section 2.4).

We infer a higher deleterious mutation rate and stronger selection in exonic compared to non-exonic sites (Fig. S19a), although we note that our estimates of selection parameters could be affected by thresholding (see Section 1.5). The total deleterious mutation rate per gamete is similar in models with and without the exonic/non-exonic division ($U=1.6$ and $U=1.73$ per gamete per generation, respectively), but the (weighted) average selection effect is greater in the model with the division ($\bar{s}=1.71\times{10}^{-3}$ vs. $\bar{s}=6.8 \times{10}^{-4}$ for the models with and without division, respectively), primarily due to stronger selection in conserved exonic sites. Overall, despite affording additional parameters, dividing conserved sites into exonic and non-exonic has little effect on our fits (Fig. S19b-e).

Regardless of whether we separate exonic and non-exonic conserved sites, most of the reduction in diversity levels is caused by selection in non-exonic regions. Weakly selected mutations cause a large reduction in neutral diversity levels over short genetic distances, whereas strongly selected mutations cause a weak reduction over long genetic distances; but the integral reduction in diversity levels due to weak and strong selection on a given set of deleterious mutations end up roughly equivalent (*41*). This property allows us to use estimates of the total deleterious mutation rates in conserved exonic and non-exonic regions as a rough measure of their proportional effects on neutral diversity levels, despite differences in selection effects in these regions. These estimates suggest that ~80% of deleterious mutations occur in non-exonic regions, indicating that they account for most of the reduction in linked neutral diversity (e.g., in the model with the top 6% of phastCons scores, ~84% of selected sites and ~76% of deleterious mutations are non-exonic; with the top 6% of CADD scores, ~83% of selected sites and ~85% of deleterious mutations are non-exonic; also see discussion in Section 4.6).

Given that the bulk of deleterious mutations exerting background selection occur in non-exonic regions, it is not surprising that a model including only conserved non-exonic sites fits the data only slightly worse than a model including all conserved sites as targets of selection (Fig. S20). By the same token, it is not surprising that a model including only conserved exonic sites fits the data substantially worse than models with either conserved non-exonic or all conserved sites as targets of selection (Fig. S20). Moreover, the estimate of the deleterious mutation rate per site in the exonic model is much higher than in the other two (Fig. S20a(ii)).

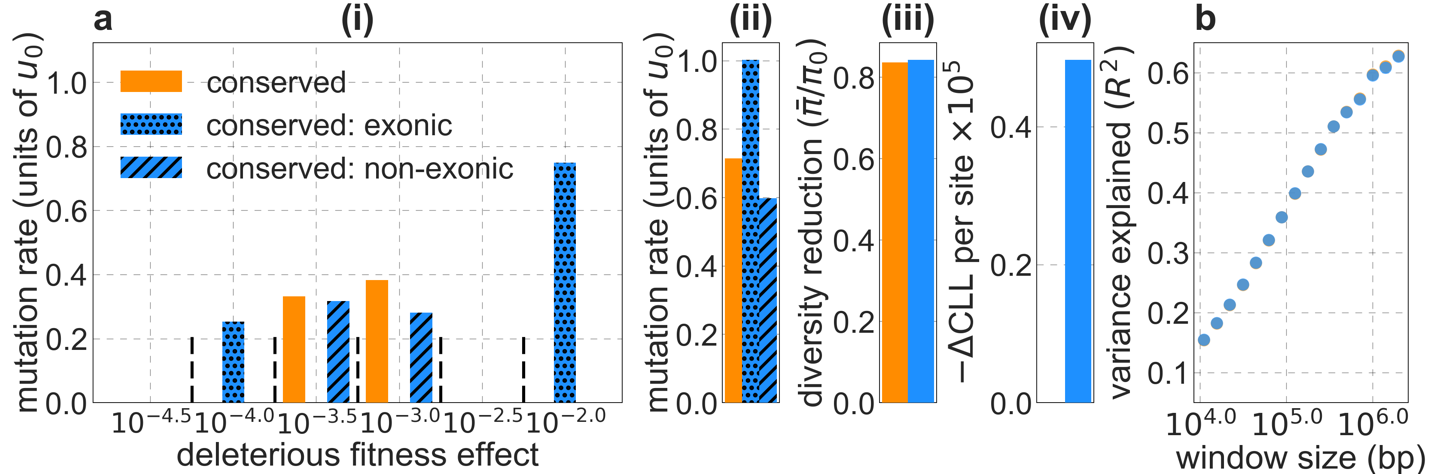

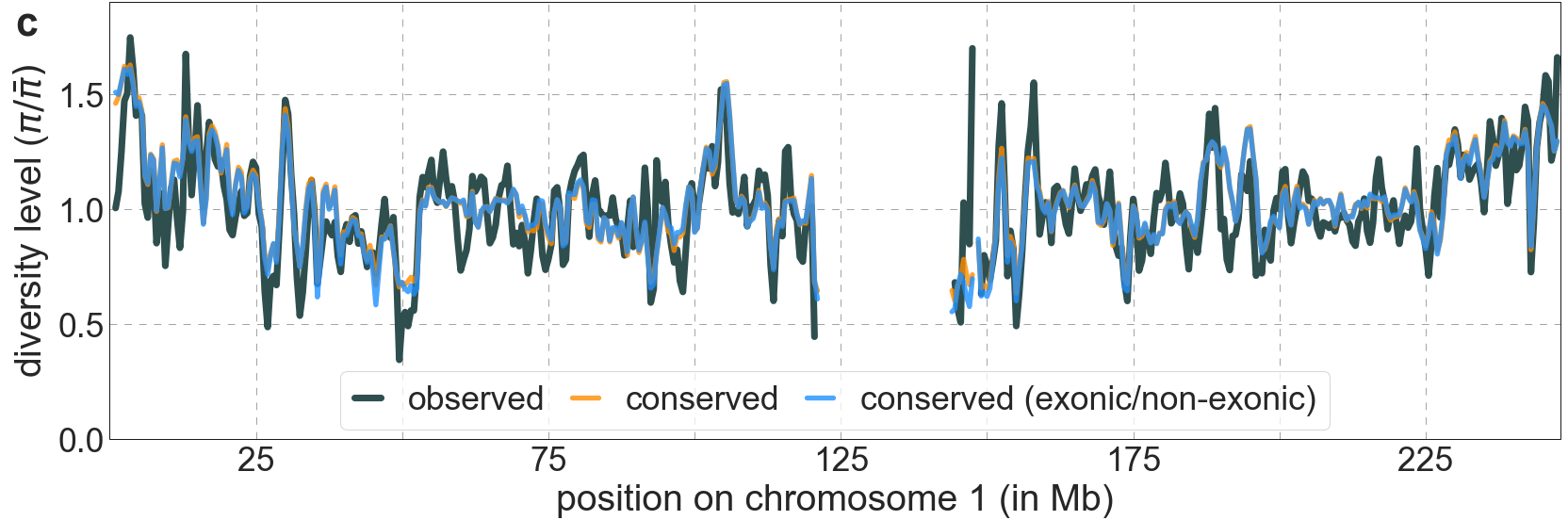

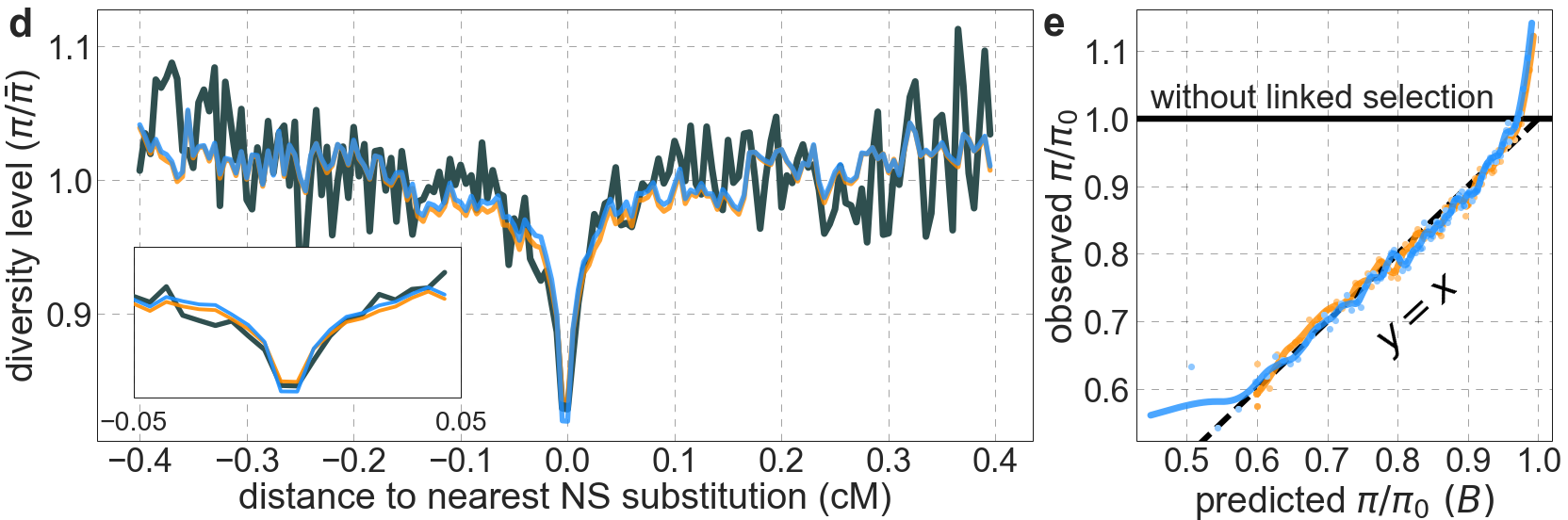

**Figure S19.** Dividing conserved sites into exonic and non-exonic sets leads to different estimates of selection parameters in each, but to little improvement in fit compared to the model based on conservation alone. Our set of conserved sites consists of the 6% of sites with the highest phastCons scores in the 99-vertebrate alignment (see Section 4.1). All panels are as described in Fig. S14. Because of thresholding (Section 1.5), the model based on conservation alone is not formally nested in the one with the division into exonic and non-exonic sets, explaining how its maximum composite-likelihood can be slightly greater.

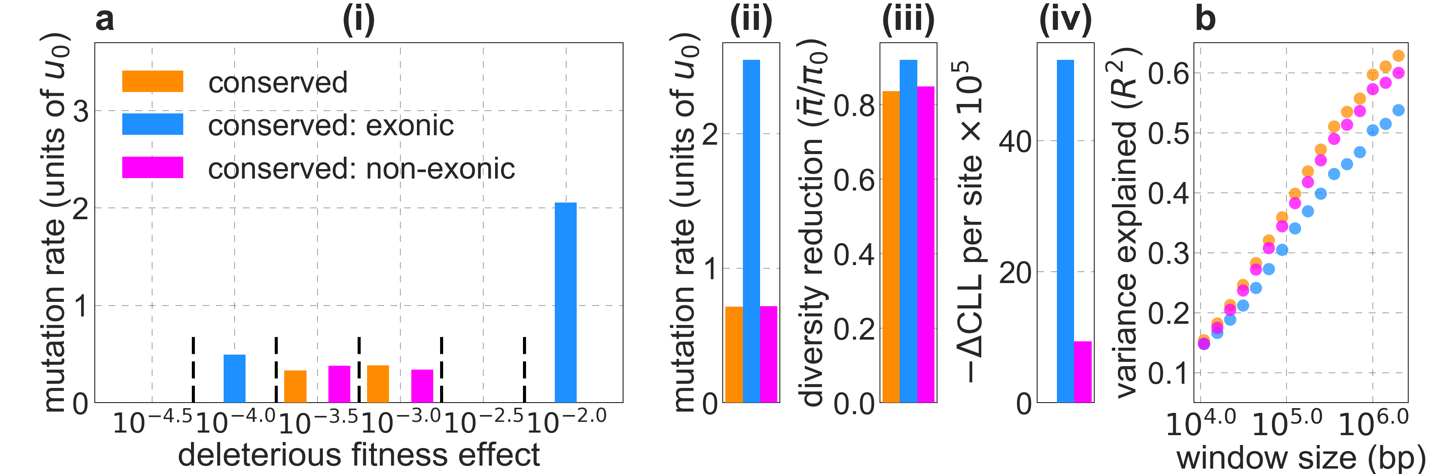

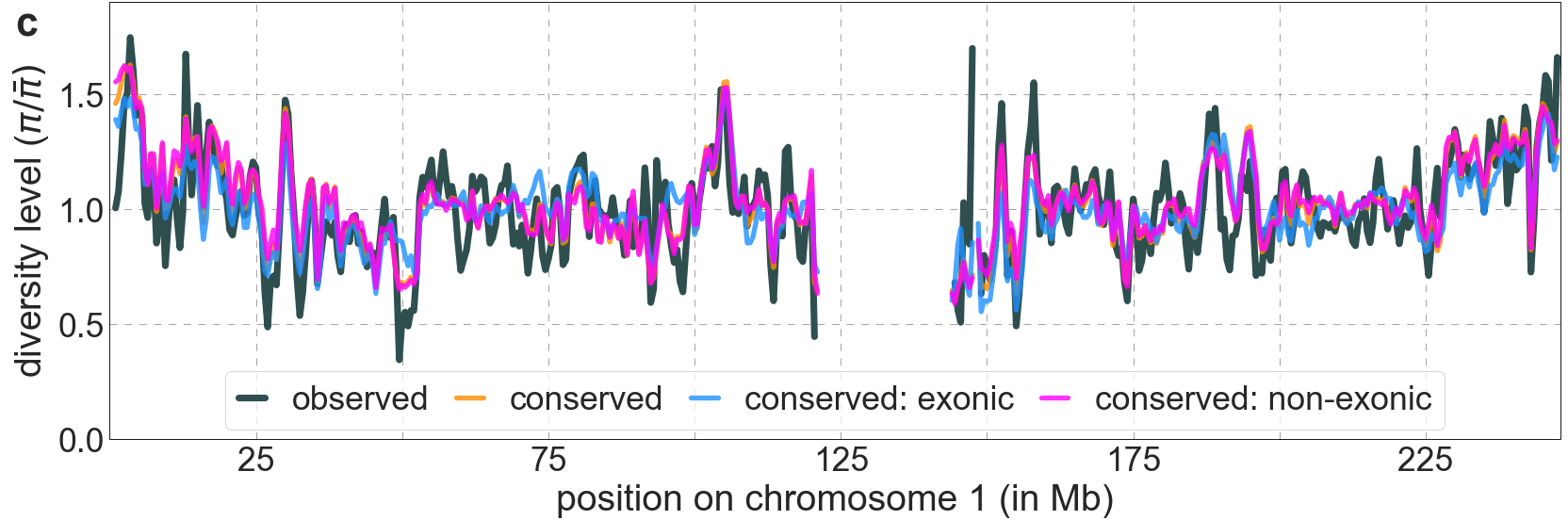

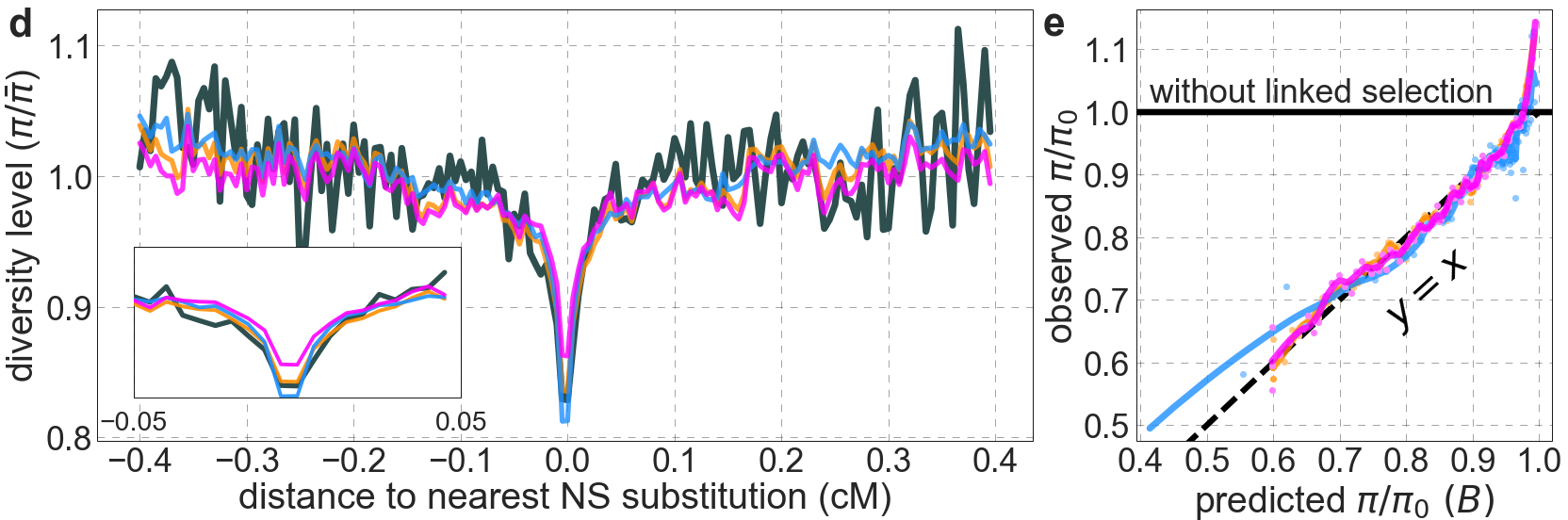
**Figure S20.** Comparison of background models using exonic, non-exonic and all conserved sites as targets of selection. Our set of conserved sites consists of the 6% of sites with the highest phastCons scores in the 99-vertebrate alignment (see Section 4.1). All panels are as described in Fig. S14.

It is somewhat surprising that the model based on conserved exonic sites alone fits the data as well as it does (Fig. S20b and c). This can be understood by noting that the spatial distribution of conserved exonic sites and of all conserved sites are fairly highly correlated (Fig. S21). Given similar spatial distributions of selected sites, the distribution of background selection effects in the model with all conserved sites can be approximated by having a higher deleterious mutation rate per site at the fewer selected sites in the exonic model. These considerations explain why we infer a similar (albeit lower) average reduction in diversity levels but a substantially higher deleterious mutation rate in the exonic model (Fig. S20a(ii) and (iii)). They also help to explain differences between our inferences and those of McVicker et al. (*2*), notably their implausibly high estimate of the deleterious mutation rate given that their main model assumes selection only at conserved exonic sites (see Main Text and Section 4.6).

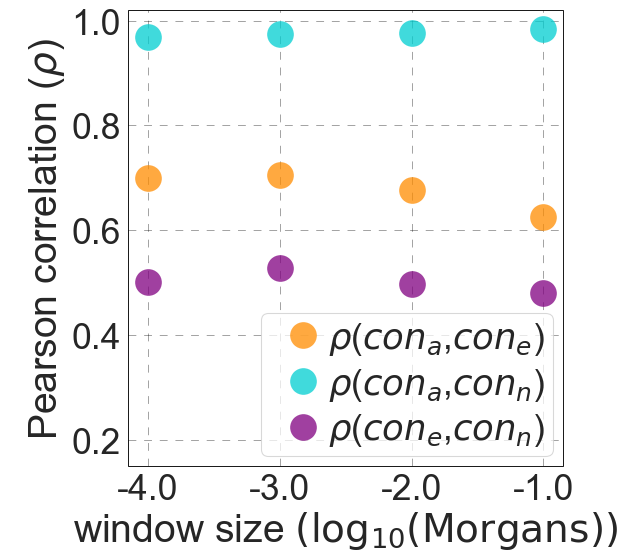

**Figure S21.** The spatial correlations of exonic, non-exonic and all conserved sites for varying window sizes (‘con_e_’, ‘con_n_’ and ‘con_a_‘, respectively).

#### 4.4 Background selection models based on other annotations

We consider two additional widely-used functional annotations as putative background selection targets. First, we rely on the expanded encyclopedias of DNA elements (ENCODE) annotations of candidate cis-regulatory elements (cCREs), including enhancer-like signatures (ELS), promoter-like signatures (PLS), CTCF-bound (CTCF) and poised/DNAse-hypersensitive (H3K4me3) assayed in 25 Tier 1a biosamples (*30*), alongside protein coding sequences (CDSs) (see Sections 2.4 and 2.6 for data sources and definition of elements). ENCODE cCREs attempt to capture the diverse repertoire of regulatory elements across cell types that control gene expression in different cellular and biological contexts. They are based on a large set of epigenomic assays, including ChIP-seq measuring the occupancy of histone marks associated with both activation and repression of gene expression, pulldown of DNA-bound transcription factors, and DNA accessibility measured in terms of DNAse sensitivity. Since we infer the majority of autosomal sites under purifying selection to be non-exonic (see Section 4.3), we reason that some combination of cCREs may substantially overlap these sites. Importantly, cCRE annotations may allow us to better partition non-exonic regions into sub-classes of sites experiencing different selection strengths. We define our choices of selection targets (other than CDSs) by grouping cCRE in two alternative ways. In one, we take the union of cCREs of a given type over all 25 biosamples. In the other, we divide cCREs of a given type into those identified in few ($\leq$ median number) or in many ($>$ median number) biosamples (in practice, most cCREs included in the first set are cell-type specific whereas most of those in the second are found in a few to all cell-types). The model in which cCREs of a given type are split performs slightly better, presumably because of the additional degrees of freedom. Both models, however, fit the data substantially worse than either of our best-fitting models (Fig. S22). The poor fit accords with the modest overlap between cCREs and our estimates of constraint sites (Fig. S23). Moreover, PLSs, the cCREs that are most highly enriched in constrained sites (Fig. S23a and c) are inferred to have a negligible deleterious mutation rate.

Next, we consider Combined Annotation-Dependent Depletion (CADD) scores (*28, 29*). CADD scores predict the ‘deleteriousness’ of every point mutation in the genome. They are generated by using machine learning to integrate information from a diverse set of annotations (122 annotations in version 1.6), such as measures of phylogenetic conservation (including phastCons scores based on the 99-vertebrate alignment), predictions of regulatory elements (including many of the assays used for constructing the

ENCODE cCREs), genic annotations (including those described in Sections 2.7 and 4.2) and predicted functional consequences of variants in protein coding sequences. The algorithm is trained using the depletion of 14.7 million high-frequency (>95%) derived alleles (based on 1000 Genomes Data) relative to 14.7 million simulated variants with the same genomic distribution as the criterion for ‘deleteriousness’. While the standard CADD scores (version 1.6) incorporate the McVicker et al. map^2^ of background selection effects as one of the annotations, we use a version in which this annotation was excluded in order to avoid circularity (see Section 2.5). We use the maximal score at each site (corresponding to the

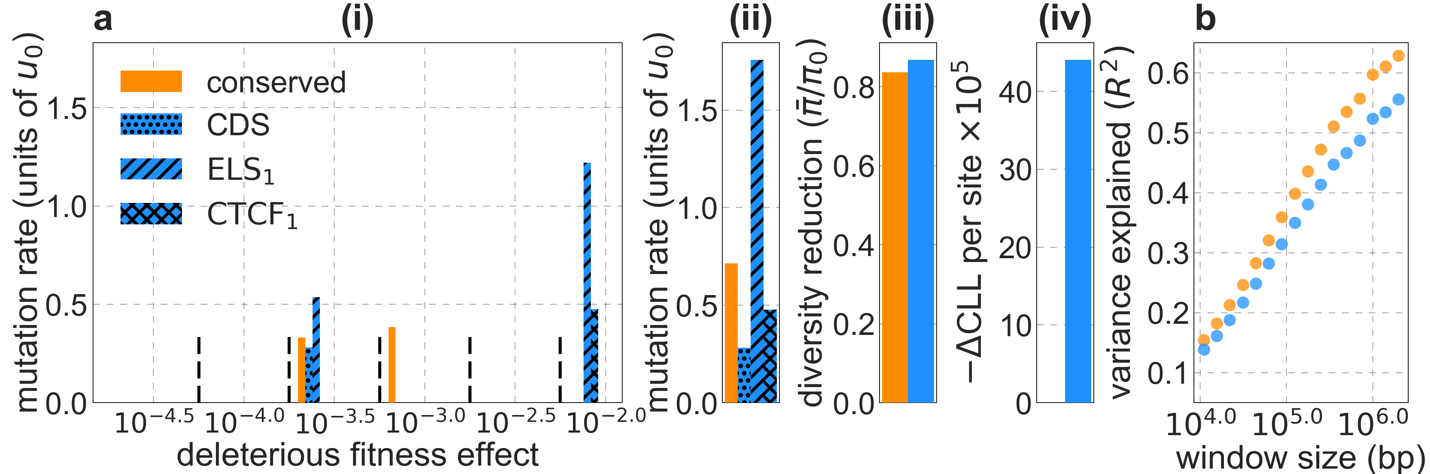

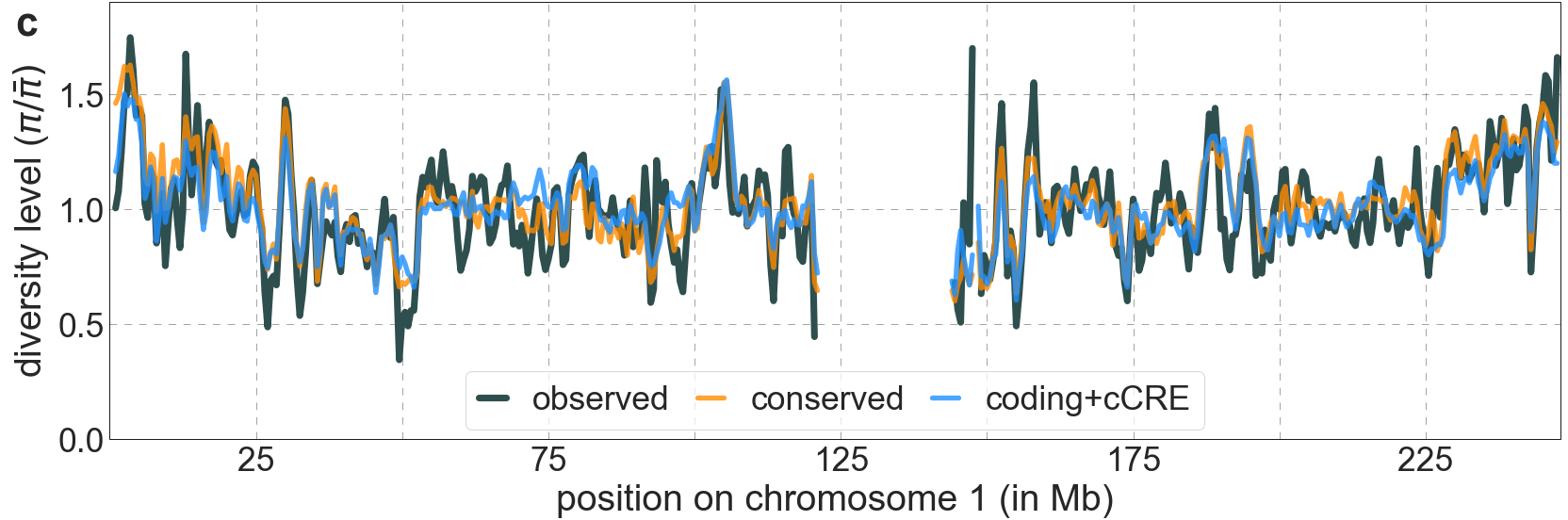

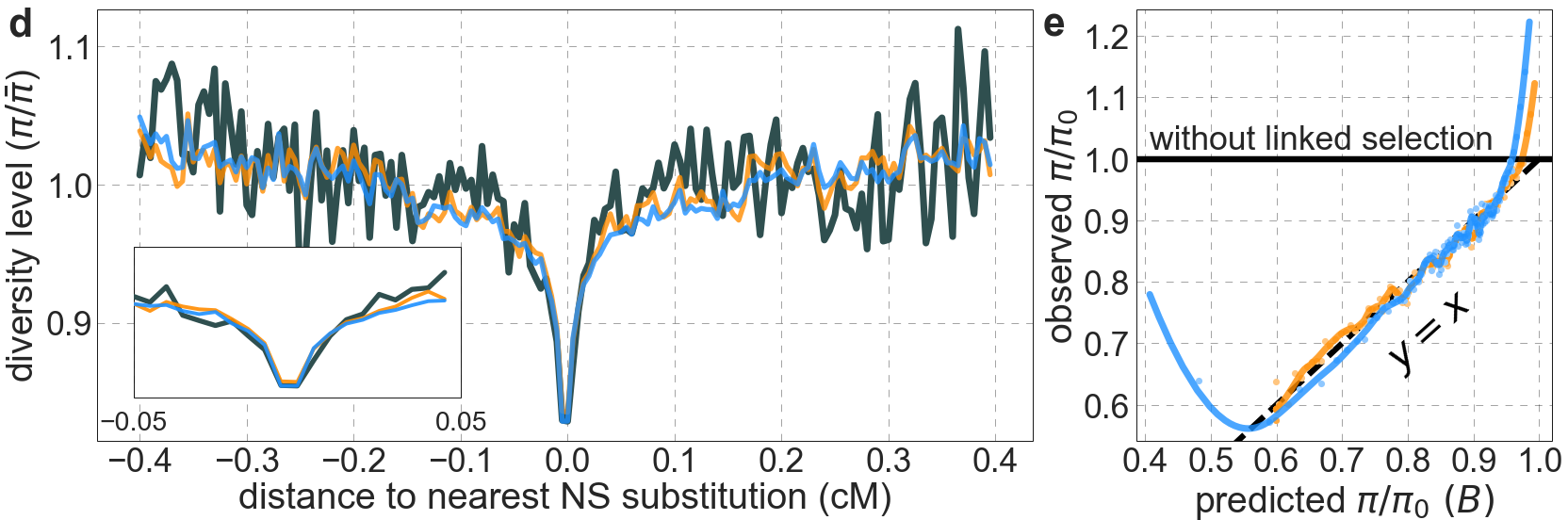

**Figure S22.** The model based on the ENCODE annotations of cCRE fit the data substantially worse than our best-fitting phastCons-based model using conservation in the 99-vertebrate alignment (conserved). The results shown correspond to the model in which we split each type of cCREs into those that occur in few (subscript 1) and many biosamples (subscript 2). We infer a non-negligible deleterious mutation rate (i.e, ${u_{d}}/{u_{0}}>0.01$) in 2 of the 8 cCRE-based putative selection targets: enhancer like sequences and CTCF binding sites identified in few biosamples, ELS_1_ and CTCF_1_ respectively, as well as in protein coding regions (CDS). All the panels are as described in Fig. S14.

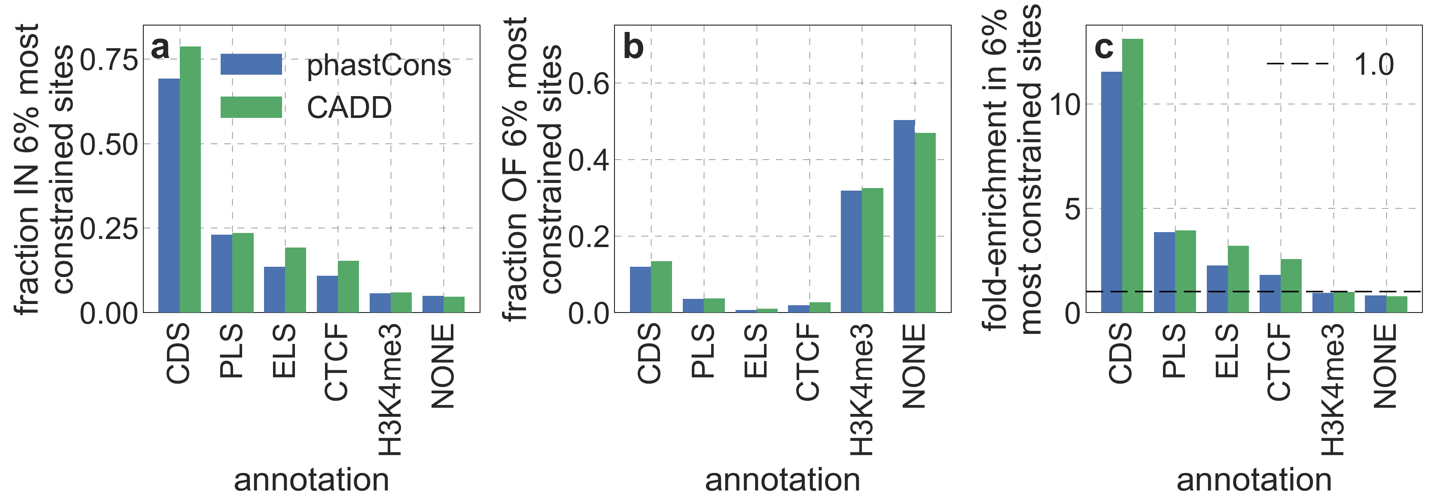

**Figure S23.** The relationship between ENCODE cCRE annotations and our main measures of constraint. Specifically, we examine the overlap of the 6% of autosomal sites with the highest phastCons or CADD scores with promoter like sequences (PLS), enhancer like sequences (ELS), CTCF-bound (CTCF), poised/DNAse-hypersensitive (H3K4me3), as well as sites that are not in any of these annotations (NONE). a) The fraction of each cCRE annotation within the 6% most constrained sites. b) The fraction of the 6% most constrained within each cCRE annotation. c) Enrichment of cCRE annotations in the 6% most constrained sites, i.e., the ratio of their proportion among constraint and all autosomal sites.

most deleterious of three possible point mutations), and, for comparison with our best-fitting phastCons-based model (Section 4.1), we begin by considering the 6% of autosomal sites with the highest CADD scores (excluding putatively neutral sites) as targets of selection.

Despite incorporating many sources of information beyond phylogenetic conservation, and doing better than phastCons scores at predicting functional consequences of variants at a single site resolution (*28*), the model based on CADD scores offers only a minor improvement over our best-fitting phastCons-based model (Fig. S24). For example, the model based on CADD scores explains 59.9% of the variance in diversity levels in 1 Mb windows compared to 59.7% for the model based on phastCons scores, although this difference and differences in other window sizes are not statistically significant (see Fig. S33 and Section 6.2). The little improvement is not that surprising, given that phylogenetic conservation is the annotation most correlated with CADD scores genome-wide (*28*), and that the spatial distributions of sites with top CADD and phastCons scores are highly correlated on the spatial scales that impact background selection effects (Fig. S25).

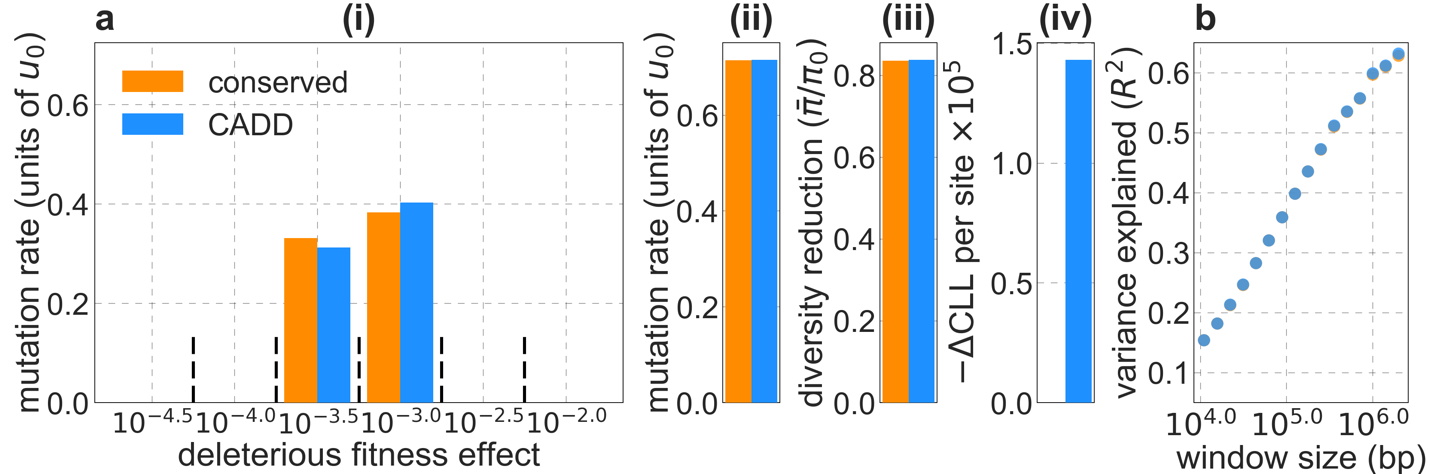

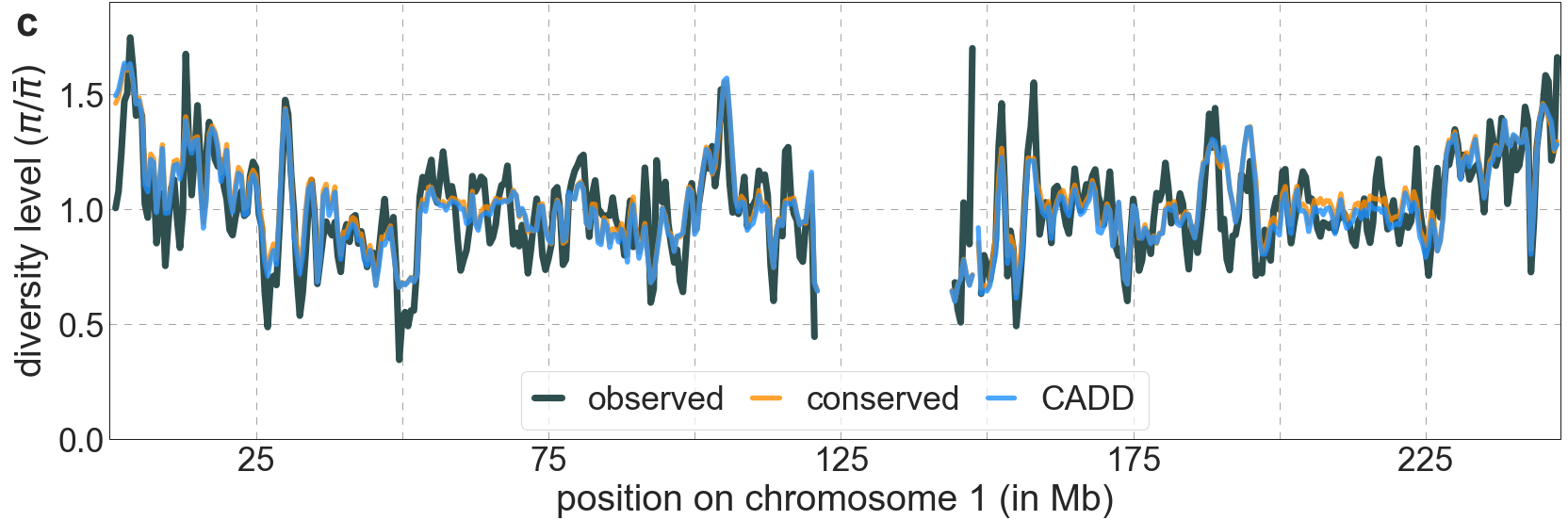

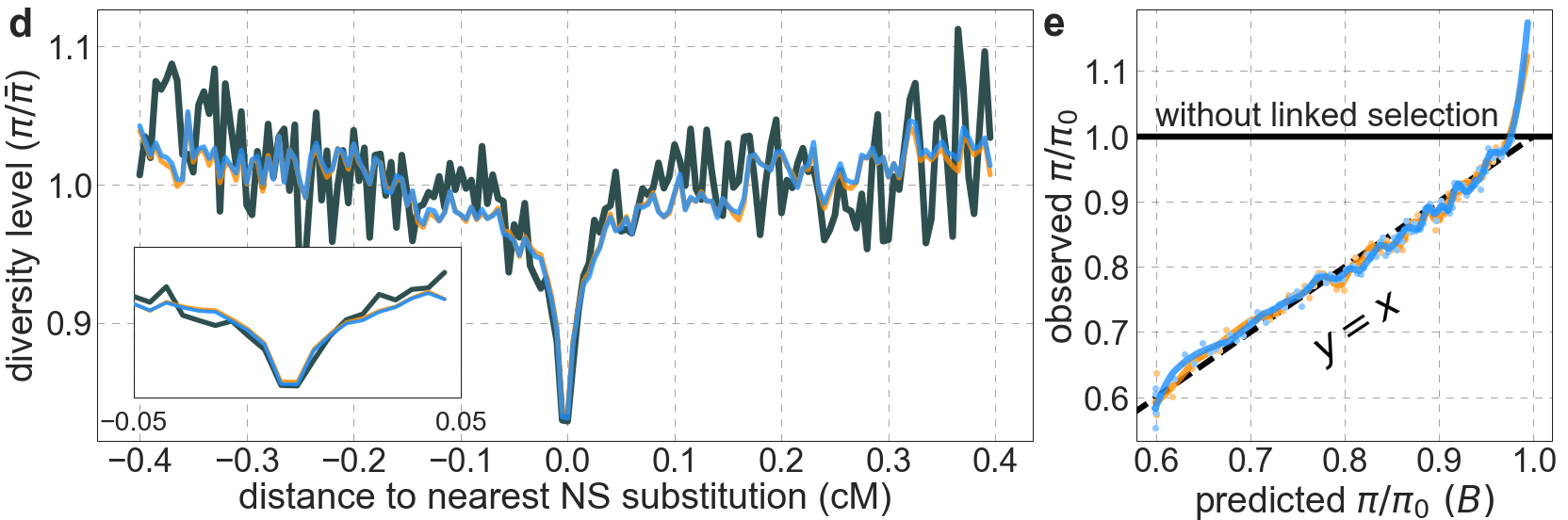

**Figure S24.** The model based on CADD scores offer little improvement over the model based on phastCons scores (based on the 99-vertebrate alignment). In both cases, we take the 6% of sites with the highest scores. All the panels are as described in Fig. S14.

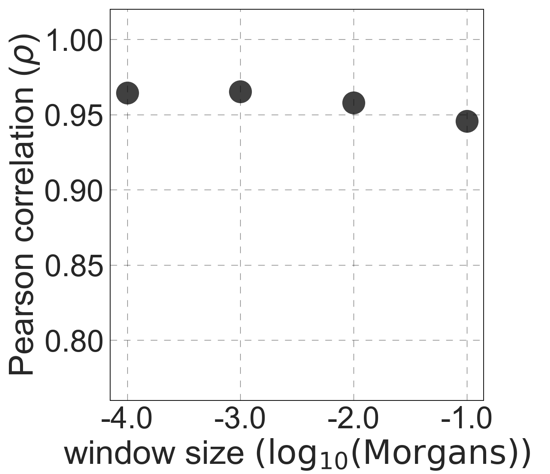

**Figure S25.** The spatial correlation between the 6% of sites with the highest CADD and phastCons scores.

The fit of models based on CADD scores is fairly insensitive to the proportion of sites included as selection targets, with proportions of 5-7% yielding slightly better fits than other choices (Fig. S26). This insensitivity and the increase in estimates of the deleterious mutation rate per site with decreasing proportion of sites used as selection targets (Fig. S26a(ii)) can be explained in the same way that we explained similar observations for models based on phastCons scores (Section 4.1).

Based on the analyses in Figs. S24 and S26, we refer to the model with the 6% of autosomal sites with the highest CADD scores as our *best-fitting CADD-based model*, and use it in most of our analyses here and in the Main Text. While the differences in fit of our best-fitting CADD-based and phastCons-based models are minor, the improved predictions of CADD compared to phastCons scores at the single site resolution substantially affects our estimates of the deleterious mutation rate based on evolutionary rates and thus their agreement with estimates based on the effects of background selection (see Main Text and Section 5).

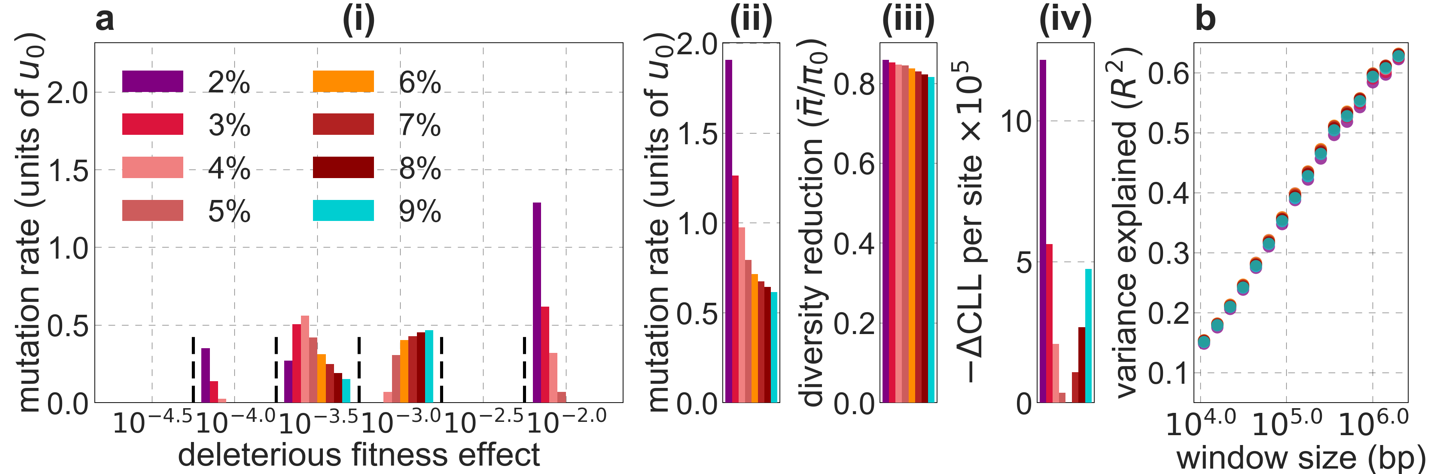

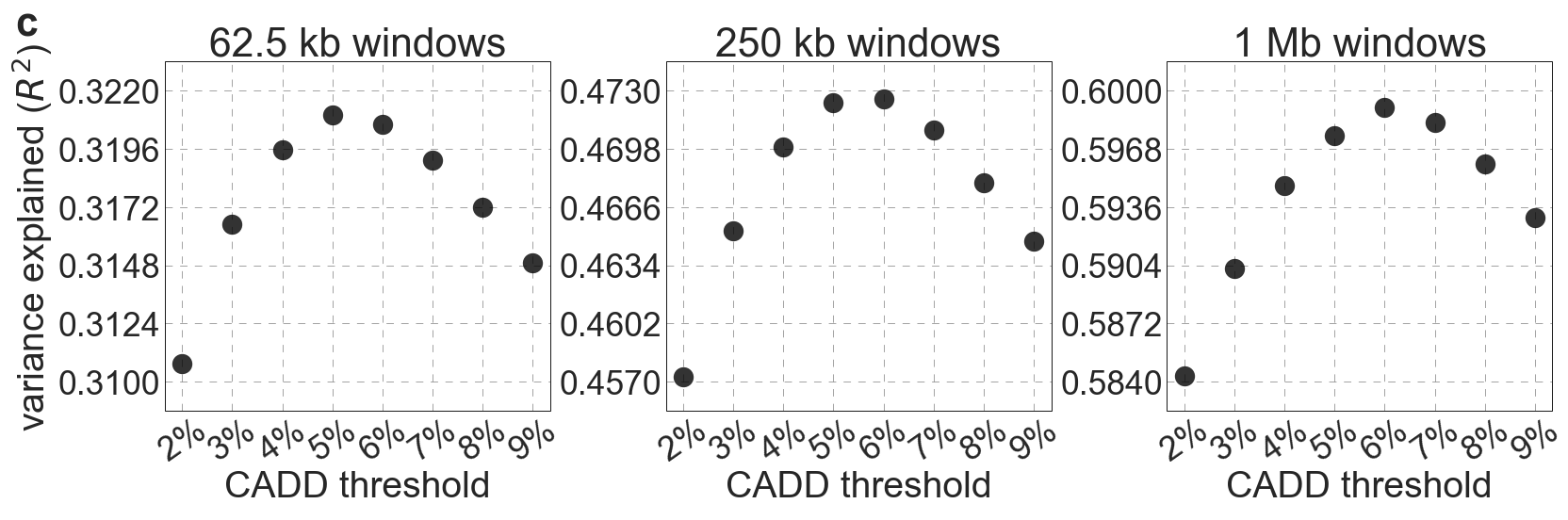

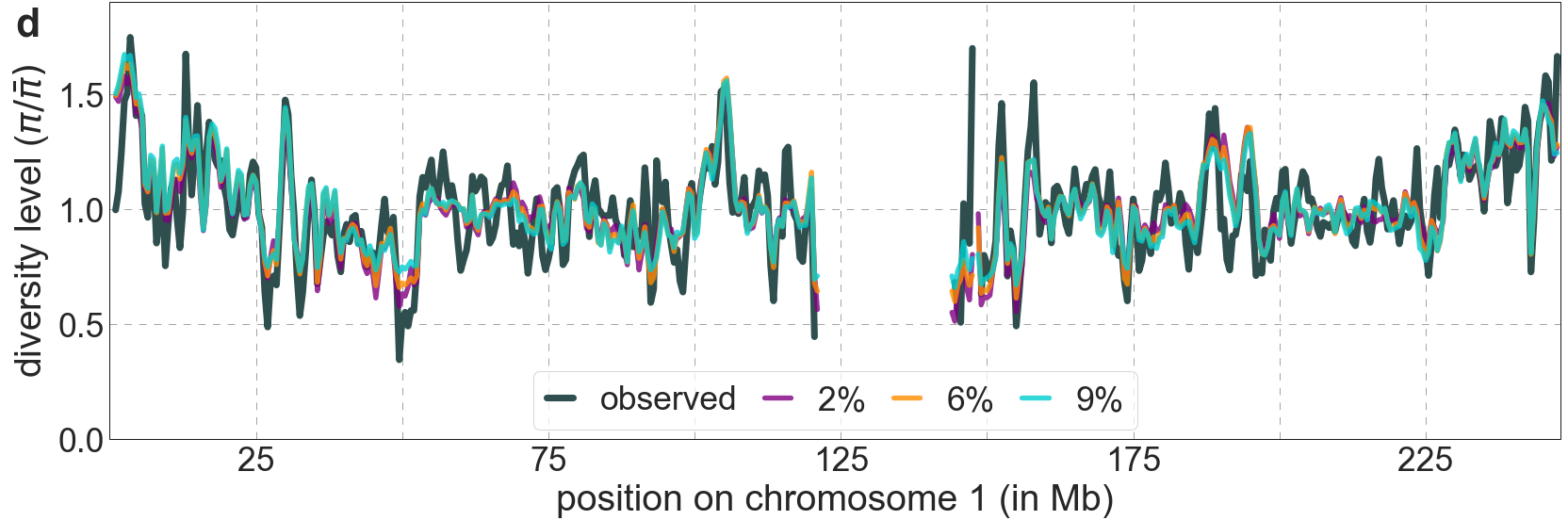

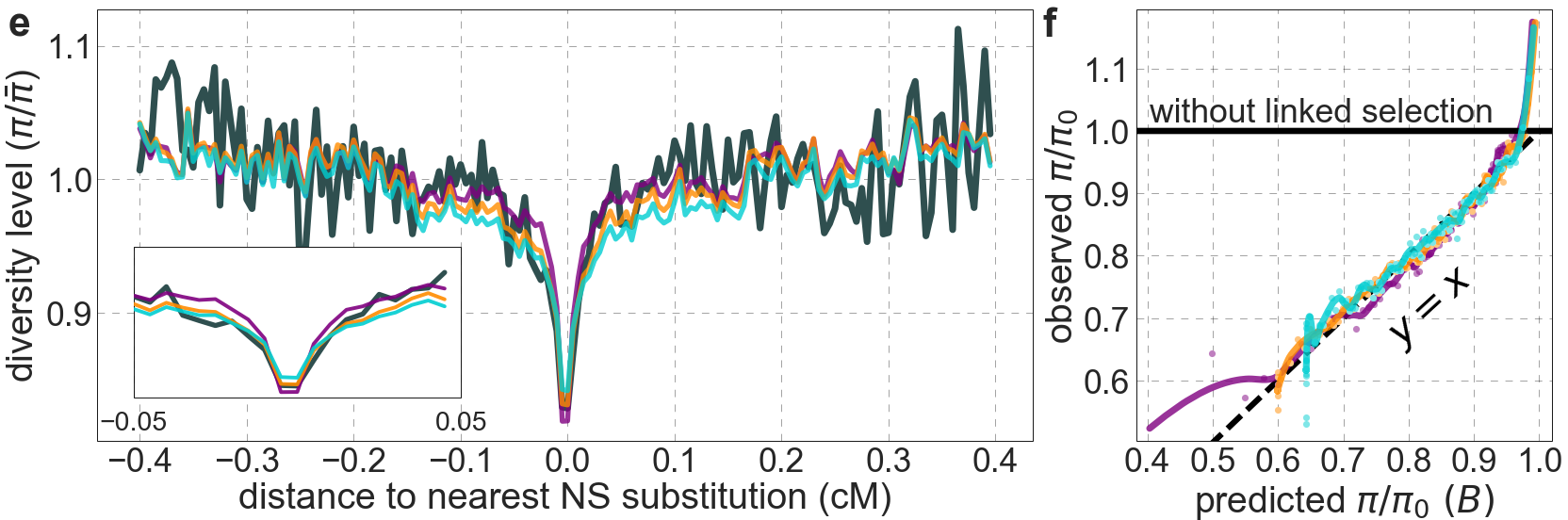

**Figure S26.** Comparison of background selection models based on CADD scores using different proportions of autosomal sites as selection targets. All panels are as described in Fig. S14.

#### 4.5 Models with selective sweeps

Next, we examine whether models that include both background selection and selective sweeps fit the data better than models with background selection alone. Our inference should be able to tease apart the effects of sweeps, primarily because these effects, unlike those of background selection, are centered around the locations of substitutions. This feature should hold true for hard, partial or soft sweeps (*8, 42-46*), so long as they result in substitutions and have a substantial effect on diversity levels (SOM Section D in Elyashiv et al. (*1*)). Indeed, previous work that applied a similar methodology to data from *Drosophila melanogaster* was able to identify and quantify distinct signatures of background selection and sweeps alongside one another (*1*).

We consider a variety of models characterized by different sets of putatively selected sites. For background selection, we consider the two sets used in our best-fitting models based on phastCons and CADD scores. We also consider several choices for targets of *positive* *selection*, i.e., for sweeps, corresponding to different kinds of substitutions that we infer to have occurred on the human lineage from the common ancestor with chimpanzees (see Section 2.7). Notably, we consider models that include the set of all nonsynonymous substitutions paired with either of the two sets for background selection. We also consider models with the substitutions that have occurred at sites with the top 2%, 3%, …, 9% of phastCons or CADD scores, where in each case we separate substitutions into sets of nonsynonymous and other, and pair that choice with the corresponding set for background selection (i.e., based on phastCons or CADD scores). For each of these choices, we infer the set of substitutions on the human lineage in two ways, either comparing the estimated human-chimpanzee ancestral genome (*31*) with the human reference genome (hg19) or with a population (YRI or CEU) sample of human genomes (see Section 2.7). We perform the inference for all of these models (18 in total) using the same grid of selection coefficients for each of the sets of selected sites, and data from either YRI or CEU. *In all cases, our estimate of the fraction of beneficial substitutions,* $\alpha$*, is essentially 0* ($<{10}^{-9}$). We do not show the results because they are indistinguishable from those for the corresponding models with background selection alone (i.e., see Figs. S16 and S26).

We also consider models with sweeps alone. Fig. S27 shows the results for a subset of these models, including the best-fitting one (e.g., based on variance explained). These models fit the data substantially worse than those with background selection alone, as seen by each of our measures (interestingly, even when considering the reduction in diversity levels around nonsynonymous substitutions; Fig. S27d). Sweep models do account for substantial variance in diversity levels, but given that they add nothing to a model of background selection alone yet fit much worse, this is plausibly because they approximate some of the effects of background selection. Notably, both background selection and sweeps cause reductions in diversity levels near selected sites, and the densities of sites that give rise to background selection and sweeps in the corresponding models are spatially correlated along the genome (Fig. S28). Moreover, the sweep models that fit the data best are those that rely on substitutions whose spatial distributions are the most highly correlated with the distributions of selection targets in our best-fitting background selection models (e.g., compare the fits and correlations for the models based on substitutions in the most conserved 2% and 9% of sites in Figs. S27 and S28). Taken together, the evidence presented here supports previous studies (*47, 48*) indicating that sweeps had little effect on current diversity levels and that background selection is the dominant mode of linked selection in humans.

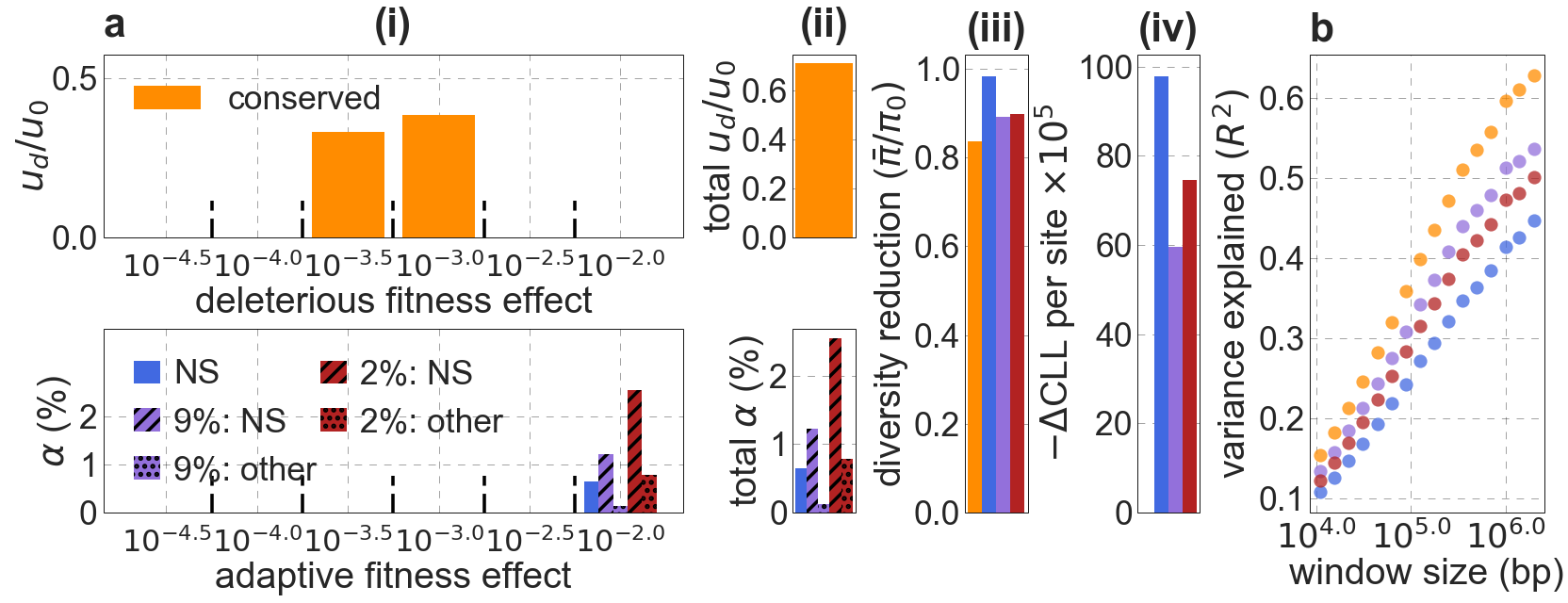

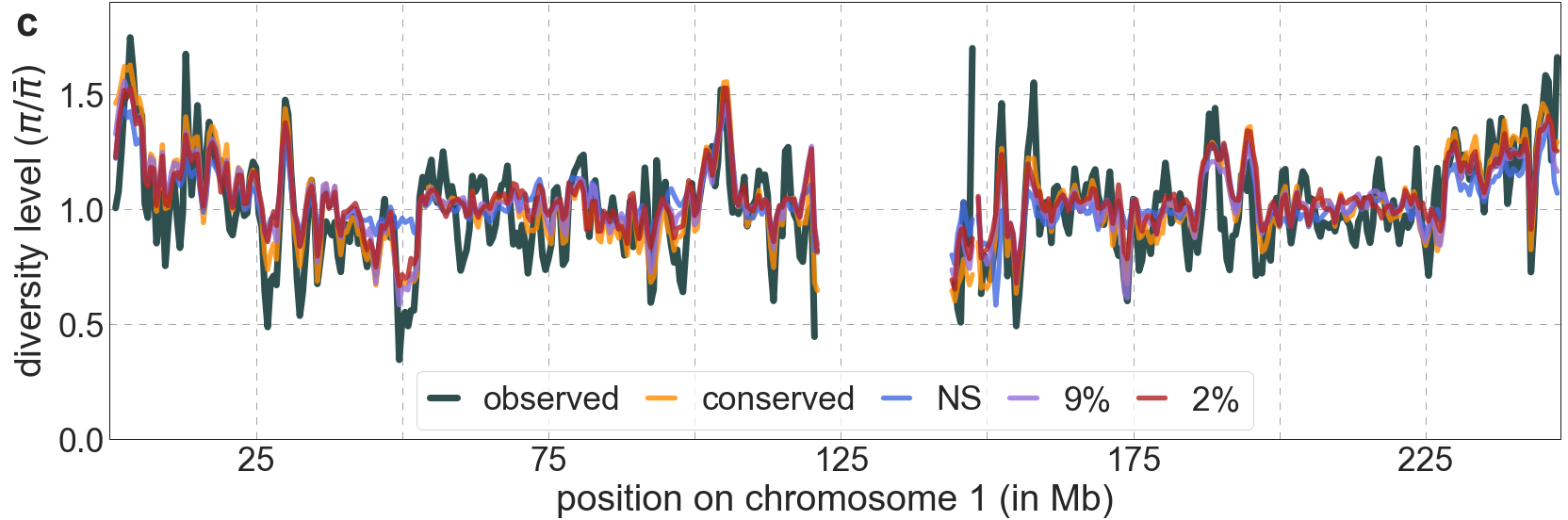

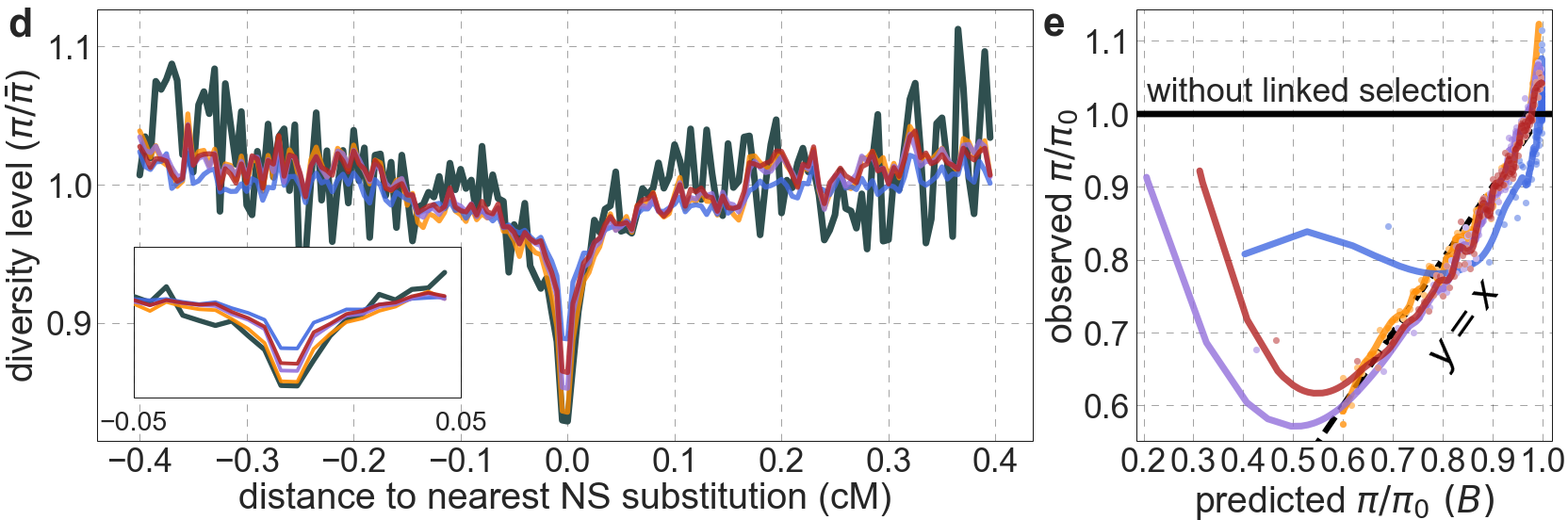

**Figure S27.** Models with sweeps alone fit substantially worse than models with background selection alone. Shown are the results for sweep models based on either: all nonsynonymous substitutions (NS); nonsynonymous and other substitutions at sites within the top 9% of phastCons scores (9%: NS/other); or nonsynonymous and other substitutions at sites within the top 2% of phastCons scores (2%: NS/other). For comparison, we also show the results of our best-fitting phastCons-based background selection model (conserved). The panels are as described in Fig. S14, with the exception of the bottom halves of panels a(i) and (ii), which show the proportions of substitutions that are estimated to be adaptive for a given selection coefficient (i) or in total (ii), for different annotations and sweeps models.

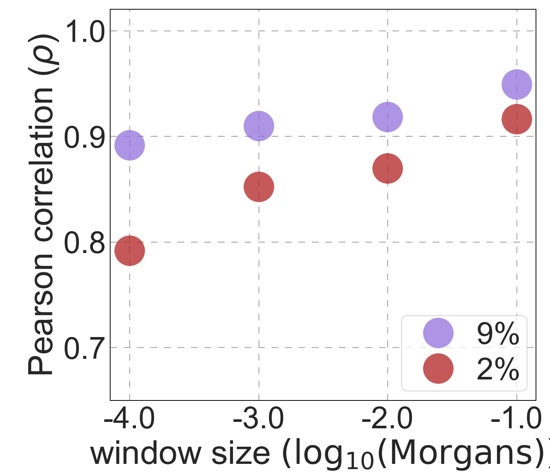

**Figure S28.** The spatial correlation between targets of selection in sweep models and in our best-fitting phastCons-based background selection model. Results shown for sweep models based on human-specific substitutions at sites within the top 2% and 9% of phastCons scores (see text for details).

#### 4.6 Comparison with previous work by McVicker et al.

For completeness, we conclude by comparing our inferences about the effects of background selection with those of McVicker et al. (*2*). The McVicker et al. study was done more than a decade ago, before genome-wide resequencing polymorphism data were available. Instead, they ingeniously used a five-primate alignment of ~4.7 million putatively neutral sites, relying on incomplete lineage sorting between human, chimpanzee and gorilla in order to learn about variation in the effective population size along the genome of the common ancestor of humans and chimpanzees. We rely on diversity levels in samples of 108 individuals at ~653 million putatively neutral sites (Section 2.1). Similar to this study, they relied on conservation scores and estimates of neutral substitution rates based on multiple sequence alignments, but they based themselves on the genomes of 15 placental mammals when we have 99 aligned vertebrate genomes at our disposal (Section 2.2). Lastly, they used a genetic map based on LD patterns (*49*), whereas we rely on genetic maps based on ancestry switches in African Americans (*24*). The McVicker study also differed in several aspects of the methodology. Notably, McVicker et al. did not incorporate selective sweeps into their models, and were therefore unable to exclude the possibility

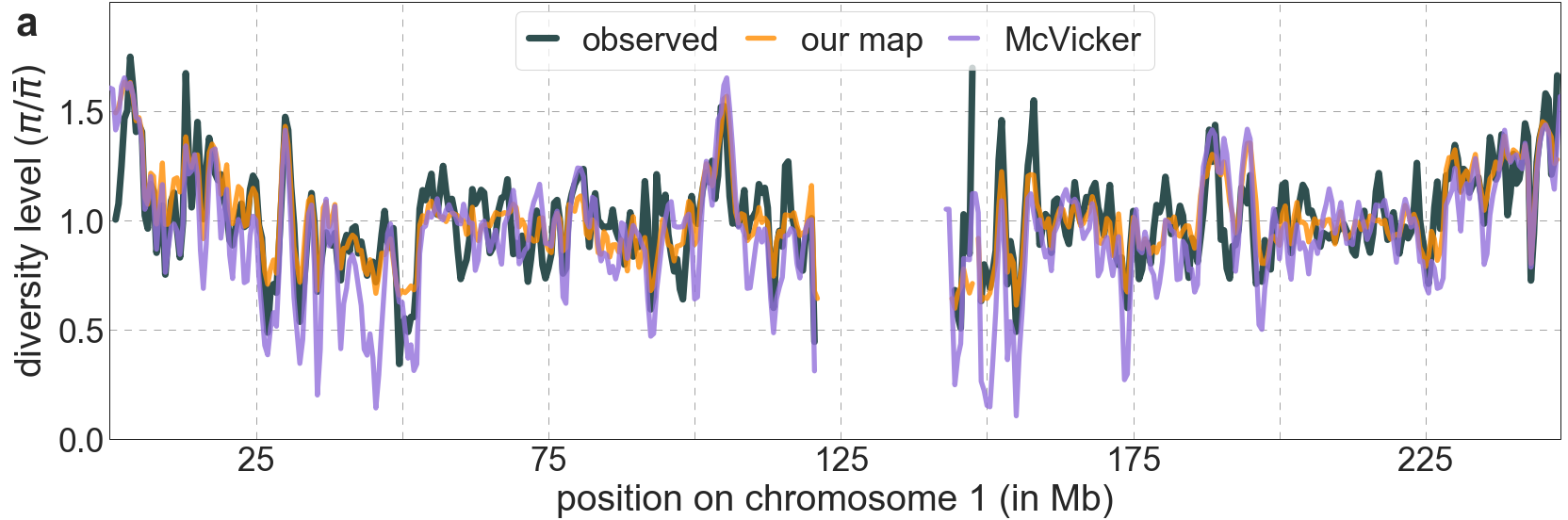

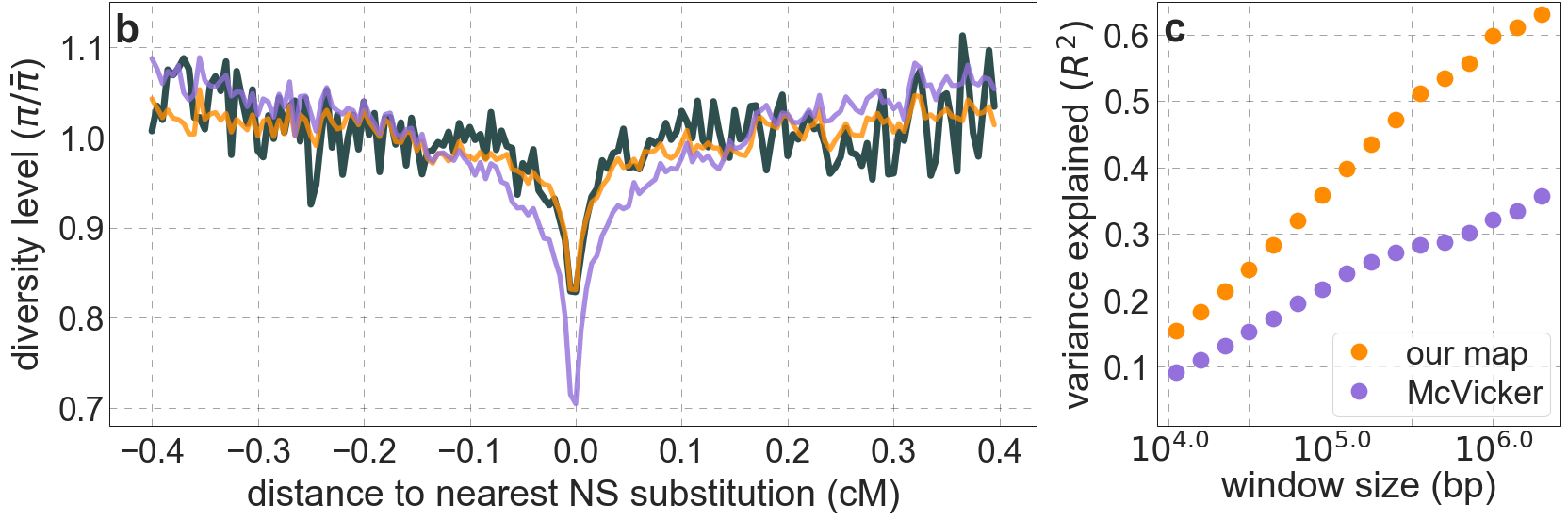

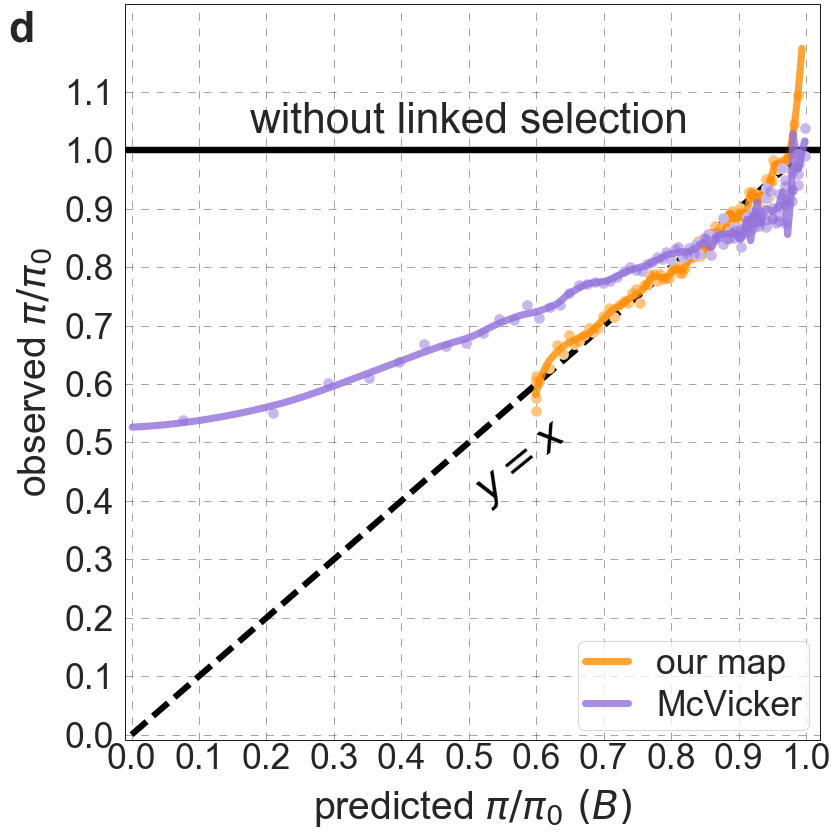

**Figure S29.** Our maps of the effects of background selection fit the data much better than the maps from McVicker et al. (*2*). Shown are the results for our best-fitting CADD-based model. All panels are as described for the corresponding ones in Fig. S14.

that sweeps had made a substantial contribution to their inferred effects of background selection (*2*). Also, McVicker et al. assumed that selection coefficients are distributed exponentially, whereas we assumed a more flexible (non-parametric) distribution on a grid. Despite limitations, the McVicker et al. maps of the effects of background selection capture substantial variation in diversity levels along the human genome (Fig. S29a and Fig. 7 in McVicker et al. (*2*)).

Nonetheless, our maps of the effects of background selection fit the data substantially better than the map from McVicker et al., both quantitatively and qualitatively (Fig. S29). They explain considerably greater proportions of the variance in diversity levels across window sizes (Fig. S29c); for example, they explain ~60% compared to ~32% of the variance on the 1 Mb scale. Our predictions are well calibrated, whereas those of McVicker et al. are not (Fig. S29d). Our predictions also do substantially better at capturing diversity patterns near specific genomic features, as illustrated by the fit to diversity levels around nonsynonymous substitutions (Fig. S29b). The relatively poor quantitative fit of the McVicker et al. predictions around synonymous and nonsynonymous substitutions (*47*) was used to argue that the effects of background selection could be more pronounced around synonymous than nonsynonymous substitutions, thereby masking the effects of selective sweeps (*50*). In this regard, the close fit of our predictions helps to refute one of two arguments for a residual, important role of selective sweeps.

We turn to the second argument, regarding estimates of the deleterious mutation rate, next. Our work and that of McVicker et al. differ markedly in our inferences about the rate and genomic distribution of deleterious mutations causing background selection in humans. In fact, the main problem in interpreting the McVicker et al. findings in terms of background selection alone is that they are based on an estimated deleterious mutation rate of $7.4\times{10}^{-8}$ per generation at their ‘conserved exonic’ sites (defined as sites within the top 5.3% of conservation scores in segments that overlap exons, accounting for ~1.1% of euchromatic autosomal sites) – more than fivefold higher than current estimates of the total mutation rate per site (see next Section). In contrast, as we detail in the next section, our estimates of the deleterious mutation rate per selected site are quite plausible ($1.00\times{10}^{-8}$ per generation for both of our best-fitting models based on phastCons and CADD scores; Fig. 4 in Main Text). The results of McVicker et al. further suggest that background selection arises predominantly from deleterious mutations in the ‘conserved exonic’ regions covering ~1.1% of euchromatic autosomal sites (i.e., they estimate ~2.3 mutations per gamete per generation in such regions in exons compared to ~0.1 elsewhere). In contrast, our results suggest that background selection arises mostly from deleterious mutations at non-exonic sites (i.e., from ~1.22 and ~1.27 mutations per gamete per generation in non-exonic compared to ~0.38 and ~0.23 mutations in exonic sites in the models based on phastCons and CADD scores, respectively). Notably, in our best-fitting models, these deleterious mutations occur in 6% of autosomal sites as opposed to only ~1% in the McVicker et al. model. Having the effects of background selection arise from deleterious mutations in a substantially greater fraction of the genome largely explains why our estimates of the deleterious mutation rate are much lower and much more plausible (Fig. 4 and S30).

### 5. Assessing estimates of the deleterious mutation rate

Here, we consider the plausibility of the deleterious mutation rate that we estimated by fitting models of background selection. First, we consider the total mutation rate per site in humans, which provides an upper bound on the deleterious mutation rate. Second, we rely on the reduction in substitution rates at our selection targets relative to putative neutral sites to obtain estimates of the proportion of mutations at selected sites that are deleterious. These estimates should be largely independent of those that we obtained by fitting background selection models, and can therefore be used to evaluate the plausibility of the latter. Lastly, we briefly consider to what extent we should expect the two kinds of estimates to line up.

#### 5.1 Estimates of the total mutation rate per site

The total mutation rate per site includes contributions from point mutations, indels, mobile element insertion (MEIs) and copy variants such as inversions. Current estimates of mutation rates per site per generation in humans are $1.2\times{10}^{-8}-1.29\times{10}^{-8}$ for point mutations (*13, 51*), $8.79\times{10}^{-10}-9.82\times{10}^{-10}$ for indels (*51*), whereas the rate for MEIs and other structural variants (including inversions and duplications) are more than two orders of magnitude lower than the point mutation rate (*52-54*), making their contribution to our calculations below negligible. Adding up point mutation and indel rates results in a per site per generation estimate of $1.29\times{10}^{-8}-1.38\times{10}^{-8}$. In estimating an upper bound on the rate of deleterious mutations at selected sites, we may consider weighting deletions by their length. For instance, we would like to count a deletion that begins at a neutral site but includes a selected site yet avoid counting one that includes multiple selected sites more than once. Counting deletions, which account for ~0.725 of indels, between once and up to their mean size of ~2.88bp (*51*), yields estimates of the total mutation rate in the range of $1.29\times{10}^{-8}-1.51\times{10}^{-8}$ per site per generation. Throughout the paper, we use the middle of this range, i.e., $1.4\times{10}^{-8}$ per site per generation, as our estimate for the total mutation rate ($u_{0}$). The estimates of the deleterious rate per putatively selected site for our best-fitting models fall well below the estimated total mutation rate (Fig. 4 and Sections 4.1 and 4.4), as one would hope.

#### 5.2 Estimating the proportion of deleterious mutations at putatively selected sites

Next, we estimate the proportional reduction of the substitution rate at selection targets relative to that at putatively neutral sites. We apply phyloFit (*18*) to the human-chimp-gorilla-orangutan (HCGO) alignment (based on the HCGO sequences from the 99-vertebrate alignment described in Section 2.2) in order to estimate the substitution rate per site on the human lineage from the ancestor with chimpanzee, for sets of selected and neutral sites (Section 3.1). To control for differences in base composition between the two sets, we estimate the reduction in substitution rates separately for each type of ancestral nucleotide (e.g., substitutions from G>X), and weight the proportional reductions by the proportions of

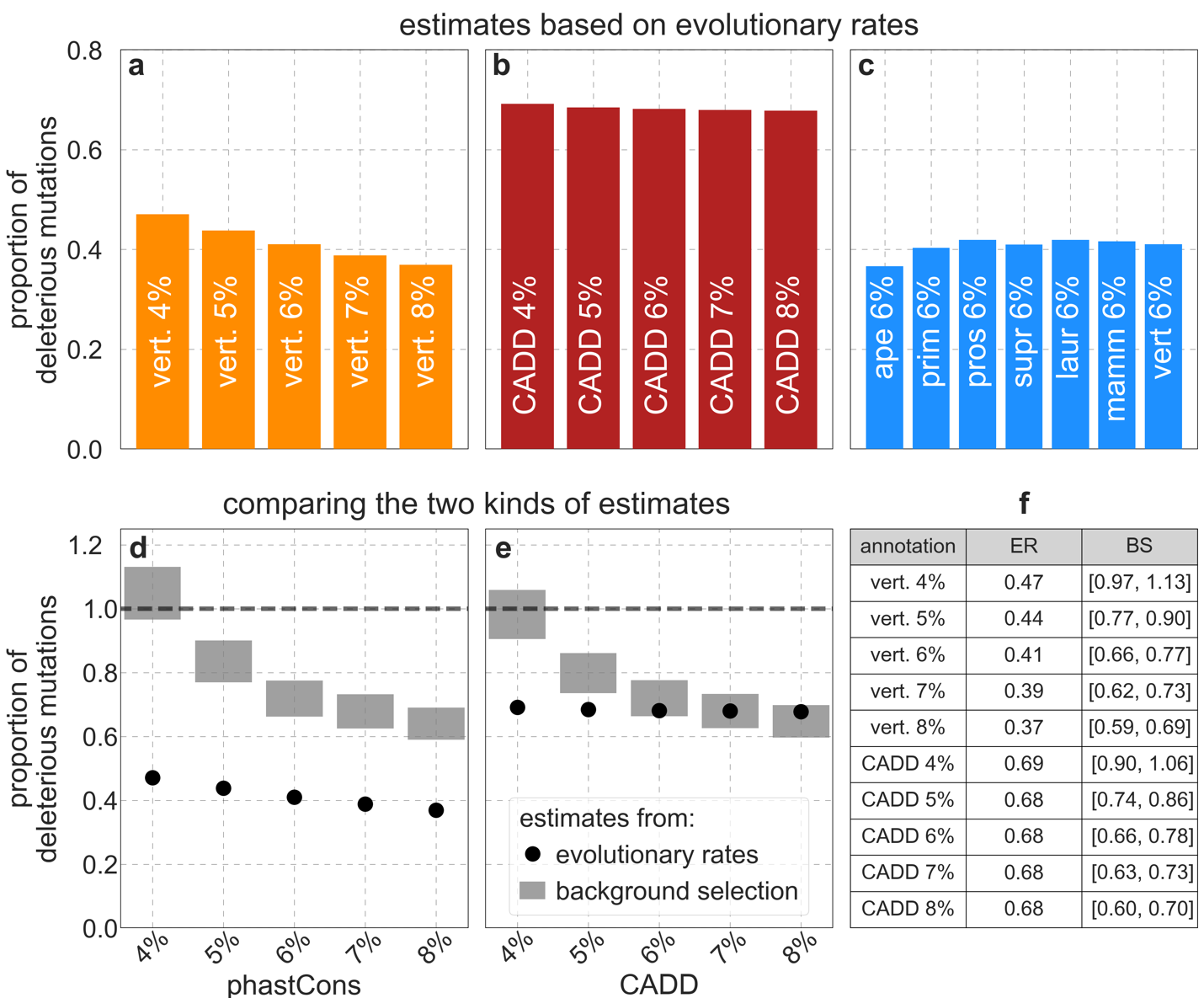

**Figure S30.** Different estimates of the deleterious mutation rate at putatively selected sites, measured relative to total mutation rates per site ($u_{0}$). Estimates based on evolutionary rates are shown for sets of selected sites chosen based on either: (a) the top 4-8% of phastCons scores for the 99-vertebrate alignment, (b) the top 4-8% of CADD scores, or (c) the top 6% of phastCons scores for alignments of varying phylogenetic depths. As expected, (see Section 4.1), the estimates are fairly insensitive to the phylogenetic depth (c). d and e) Estimates based on evolutionary rates vs. those based on the effects of background selection, for sets of putatively selected sites based on phastCons scores for the 99-vertebrate alignment (d) and on CADD scores (e). f) Estimates for different sets of putatively selected sites based on evolutionary rates (ER) and background selection effects (BS). The range of estimates based on background selection effects (in d-f) is due to the uncertainty about the total mutation rate per site (Section 5.1).

each nucleotide in the set of selected sites. Controlling for the composition of triplets rather than single nucleotides produces similar estimates. Note that in choosing our sets of neutral and selected sites based on phylogenetic conservation (Sections 3.1 and 4.1), we excluded the human genome from the alignments and therefore our estimates of the reduction in substitution rates on the human branch should be minimally confounded with the choice of sites. Similarly, the conservation scores that serve as input for calculating CADD scores are based on the same 99-vertebrate alignment excluding the human reference genome (see Supplementary Table 1 in Kircher et al. (*28*)).

The estimates of the proportion of mutations that are deleterious are shown in Fig. S30 (and Fig. 4), along with their comparison to estimates from background selection models. Expectedly, estimates based on substitution rates decline slightly as the cutoff phastCons or CADD score decreases (i.e., as the percentage of sites included in the selected set increases) (Fig. S30a and b). Importantly, estimates based on substitution rates are substantially greater for the sets chosen based on CADD than on phastCons scores (Fig. S30a and b), whereas estimates based on background selection effects are similar in both cases (Fig. S30d and e). We interpret this finding as reflecting the greater ability of CADD scores to identify selection on a single site resolution (*28*), plausibly because CADD scores incorporate measures of phylogenetic conservation based on one site at a time (e.g., phyloP, GERP (*55, 56*)) in addition to measures that rely on runs of sites, such as phastCons scores. Consequently, the two estimates of the deleterious mutation rate are within a factor of 2 for our best-fitting phastCons-based model whereas they overlap for our best-fitting CADD-based model, while the estimates based on background selection effects are similar in both cases (Fig. S30d and e).

#### 5.3 Interpreting the relationship between the two estimates

We would expect the two estimators of the deleterious mutation rate to yield similar but not identical answers. For one, the range of selection coefficients that cause a substantial reduction in evolutionary rates, e.g., $4N_{e}s\gtrsim3$(*57*), is greater than the range of effects that cause a substantial reduction in diversity levels via classic background selection, e.g., $4N_{e}s\gtrsim10$(*58-60*). This consideration suggests that estimates based on evolutionary rates should be greater than those based on the effects of background selection (although non-equilibrium demographic history, notably changes in population size, might complicate quantitative expectations). On the other hand, we cannot expect to identify all selected sites and only those by our criteria. Estimates based on the effects of background selection plausibly soak up much of the contribution of missing selected sites, because their spatial distribution is likely to be highly correlated with sites that are included in our sets (see Section 4.1). In contrast, estimates based on evolutionary rates are affected only by the sites in our sets and would be biased downwards by the accidental inclusion of effectively neutral sites. For these reasons, we do not expect the two estimates of the deleterious mutation rate to align perfectly. Nonetheless, it is encouraging that when we rely on selected sites that amount to current estimates of the proportion the human genome under selection, i.e., ~5-9% (*36, 40*), our two estimates of the deleterious mutation rate are quite similar. Moreover, the similarity is highest when we use CADD scores, which are better than phastCons scores at identifying selection on a single site resolution (*28*). Thus, our results resolve the issues raised by the substantial overestimation of the deleterious mutation rate in past work (*2*).

### 6. Statistics

#### 6.1 Estimates of explained variance

Our main quantitative measure for the fit of our models is the variance in diversity levels explained by our predictions, $R^{2}$, for different window sizes. A concern in using $R^{2}$ as a measure of fit is that it not be inflated by overfitting. To avoid this problem, we exclude the data in a given window from the inference used to predict diversity levels in that window. Specifically, we divide the autosomal polymorphism data into contiguous non-overlapping blocks of 2 Mb, and repeat the inference using the data excluding one block at a time. As the autosomes cover just over 2.88 Gb, this amounts to repeating our inference 1,440 times. When we calculate the contribution of a given window (of size $\leq$ 2 Mb) to $R^{2}$, we use the prediction based on excluding the 2 Mb block containing that window. In Section 6.3, we use the same datasets and inferences to calculate jackknife estimates for the sampling error of our estimates of model parameters.

Fig. S31 shows the relative difference between our $R^{2}$ estimates using all the data and this exclusion approach for our two best-fitting models. These estimates suggest that overfitting has a tiny effect, which is not surprising given the large amounts of data used in our inferences. Given the negligible effect and computational burden of this analysis, we do not repeat it for each of the models we examine, and use the $R^{2}$ estimates based on predictions using all the data instead.

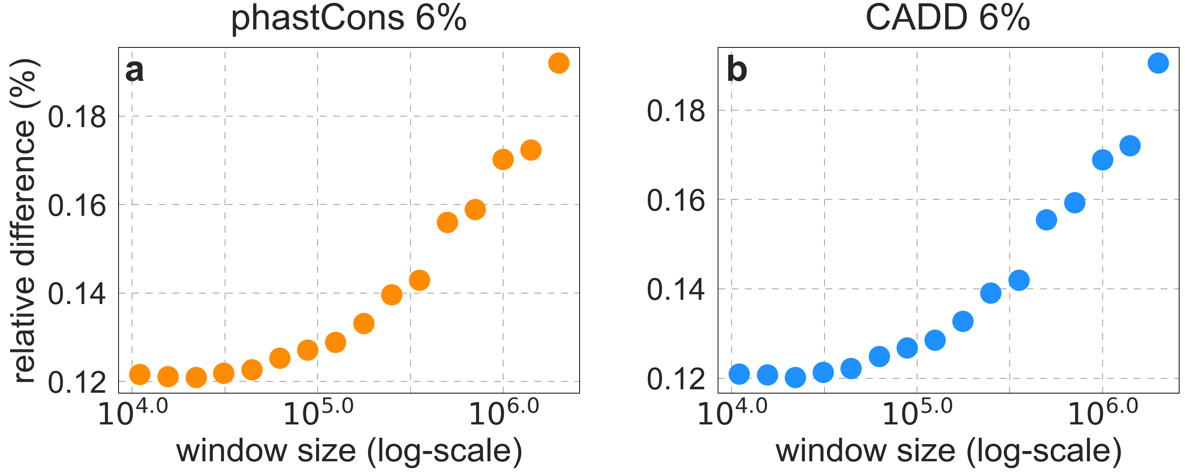

**Figure S31.** The relative difference between $R^{2}$ estimates using all the data and the exclusion approach described in the text, for our two best-fitting models.

#### 6.2 Comparing the fit of different maps

We use permutations of paired maps to test whether differences in $R^{2}$ between two maps are statistically significant. Assume without loss of generality that $R^{2}$ for a given window size is greater for map I than for map II. We divide autosomes into $2881$ contiguous non-overlapping blocks of 1 Mb and generate a new map (map A) by picking each 1 Mb block from map I or II at random; we generate the complementary map (map B) by picking the alternative 1 Mb blocks throughout. This way, we generate $n$ paired maps and calculate the difference in $R^{2}$ between each pair, $\Delta R^{2}$, to obtain a distribution for the expected differences in $R^{2}$ between maps I and II under the null hypothesis that their fit to

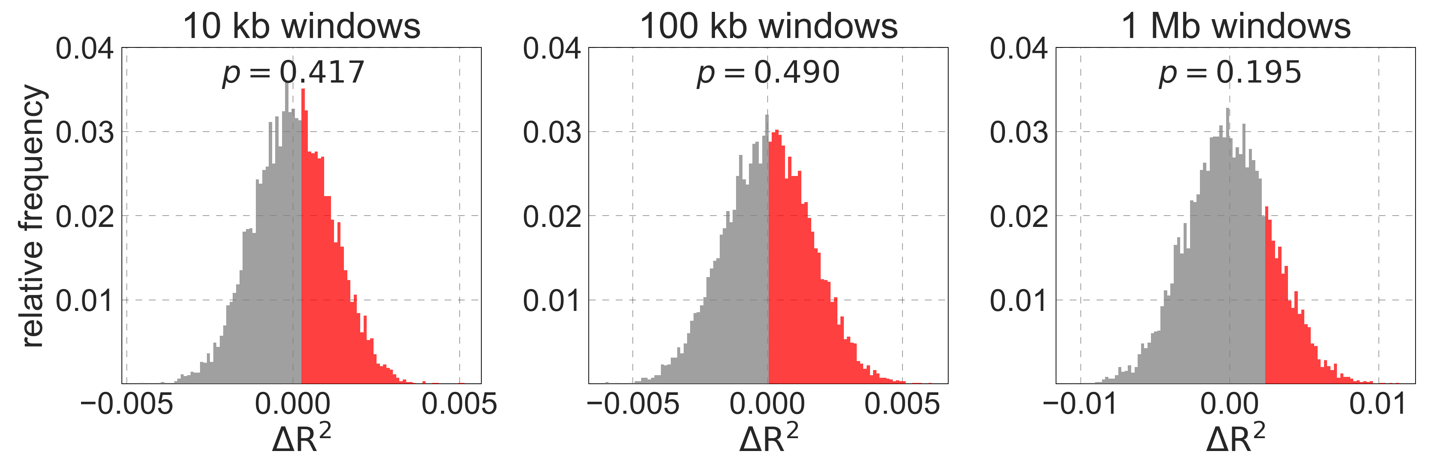

**Figure S32.** Assessing the differences in fit between our best-fitting models on three spatial scales. We show the distribution of differences in explained variance $({\Delta R}^{2})$ for 10,000 paired permutations of the best-fitting CADD and phastCons based maps. The part of the distributions with ${\Delta R}^{2}\geq{\Delta R}_{O}^{2}$ is in red, and the corresponding p-value is shown above.

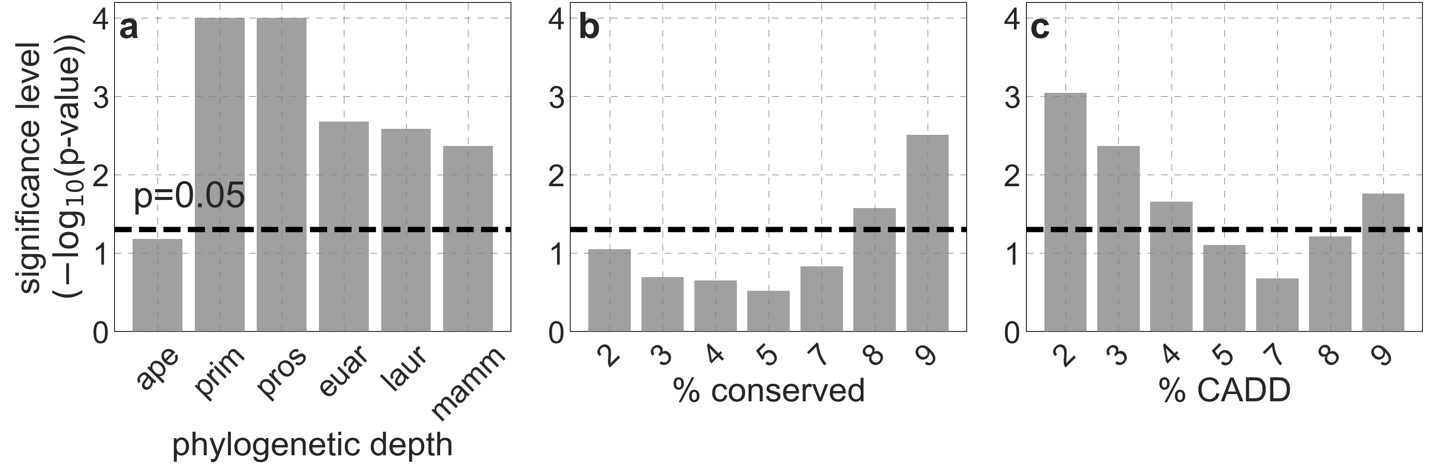

**Figure S33.** Significance level of differences in fit between our best-fitting models and variations on these models, on the 1 Mb spatial scale. a and b) Comparison of our best-fitting phastCons-based model with phastCons-based models with alternative phylogenetic depths (a) and conservation thresholds (b). c) Comparisons of our best-fitting CADD-based model with CADD-based models with alternative thresholds

polymorphism data is roughly equivalent. Having $r$ denote the number of permutations with $\Delta R^{2}$ greater than or equal to the observed difference $\Delta R_{O}^{2}$, we estimate the p-value for $\Delta R_{O}^{2}$ under the null by $p=(r+1)/(n+1)$.

We illustrate this procedure by comparing our two best-fitting models (Fig. S32). The fit of these models is very similar, with, e.g., $R^{2}=0.599$ and 0.597 at the 1 Mb scale for the models based on CADD and phastCons scores respectively. We find that the difference

between the fits is not statistically significant, supporting our claim that the functional annotations incorporated in CADD offer little or no improvement in predictive power (see Main Text and Section 4.4). Using this procedure to compare our best-fitting phastCons-based model with phastCons-based models with alternative phylogenetic depths or conservation thresholds, we only find significant differences at the 1 Mb scale in a small subset of cases (Fig. S33a and b). The same is true for comparisons of our best-fitting CADD-based model with CADD-based models with alternative thresholds (Fig. S33c). We note that even when the fits are significantly worse, they are still far closer to the fits of our best-fitting models than any of the models based on other choices of selection targets discussed in Section 4.

Interestingly, the difference in fit between the model based on conservation in 99-vertebrates and four-apes is the only non-significant comparison across phylogenetic depths (Fig. S33a), despite the fact that other phylogenetic depths have $R^{2}$ values closer to the 99-vertebrate result across various window sizes (Fig. S14c). We believe this is due to the fact that the four-ape alignment may actually better capture some recent targets of selection, but with greater noise, reflecting a tradeoff in the power to detect conservation vs. functional turnover. As a result, a non-negligible subset of 1 Mb windows in the four-ape yield better fits to the data than the 99-vertebrate map. In contrast, maps from deeper in the phylogeny are essentially highly correlated to the 99-vertebrate map, and the small differences in fit are uniformly biased in favor of the 99-vertebrate map across 1 Mb windows due to its greater power to resolve the boundaries of conserved elements.

#### 6.3 Sampling error in parameter estimates

We use a jackknife resampling approach to estimate the sampling errors of our parameter estimates (see, e.g., Patterson et al. (*61*)). To this end, we perform the inference on datasets excluding 2 Mb blocks as described in Section 6.1. Specifically, denoting the parameter of interest by $\theta$, and the estimate based on the data set excluding block $i=1,\ldots,n$ by $\hat{\theta}_{i}$, our jackknife estimates of the mean and variance are $\bar{\theta}=\frac{1}{n}\sum_{i=1}^{n} \hat{\theta}_{i}$ and $V\left( \bar{\theta} \right)= \frac{n-1}{n}\sum_{i=1}^{n} {(\hat{\theta}_{i}-\bar{\theta})}^{2}$ respectively. We use the standard deviation $\sqrt{V\left( \bar{\theta} \right)}$ as our measure of sampling error (SE). Fig. S34 shows the SEs for the parameter estimates of our two best-fitting models. As these examples illustrate, these errors are quite small and do not affect the conclusions of our analyses. Consequently, and given the computational cost of obtaining them, we do not calculate these SEs for most models.

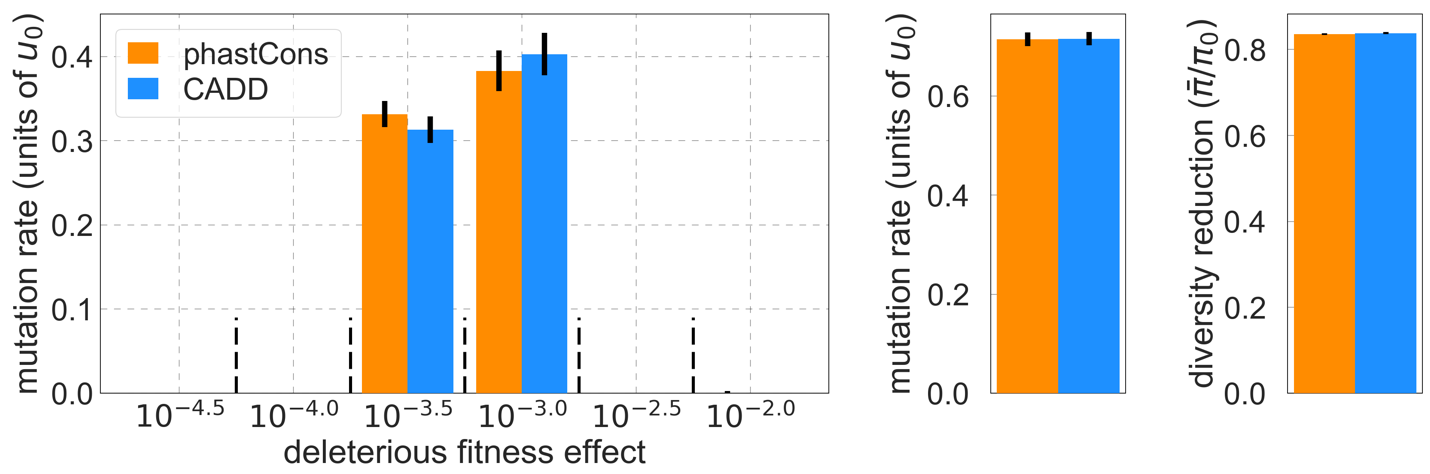

**Figure S34.** Estimates of the sampling errors of parameter estimates for our two best-fitting models. The bars denote $\pm SE$ estimated using jackknife as described in the text.

### 7. Results for other human populations

Here, we examine whether the maps of the effects of background selection that we infer and evaluate using polymorphism data from YRI provide a good fit to data from other populations. To this end, we use data from each of the other 25 populations collected in Phase III of the 1000 genomes project (*19*), which span a wide geographic range and have had different demographic histories (*19*), to infer the maps corresponding to our best-fitting models. The population-specific maps can be found at github.com/sellalab/HumanLinkedSelectionMaps.

Overall, we find that the maps and main parameters inferred in different populations are remarkably similar (Fig. S35 and S49-S51). When we compare the predictions of relative diversity levels along autosomes (i.e., relative to the mean in each population) we find nearly perfect correlations across window sizes (Fig. S35a and b). The distributions of selection effects of deleterious mutations, estimates of the total deleterious mutation rate per selected site, and the mean reduction in diversity levels, are all quite close among populations (Fig. S35c). The similarity among maps implies that we can use the maps of relative diversity levels inferred in YRI to predict diversity levels in other populations without loss of accuracy. Specifically, we multiply predictions based on YRI by a constant, chosen such that the predicted and observed mean diversity level in the focal population match. Fig. S36 illustrates that the adjusted YRI maps predict diversity levels as well as the population specific ones.

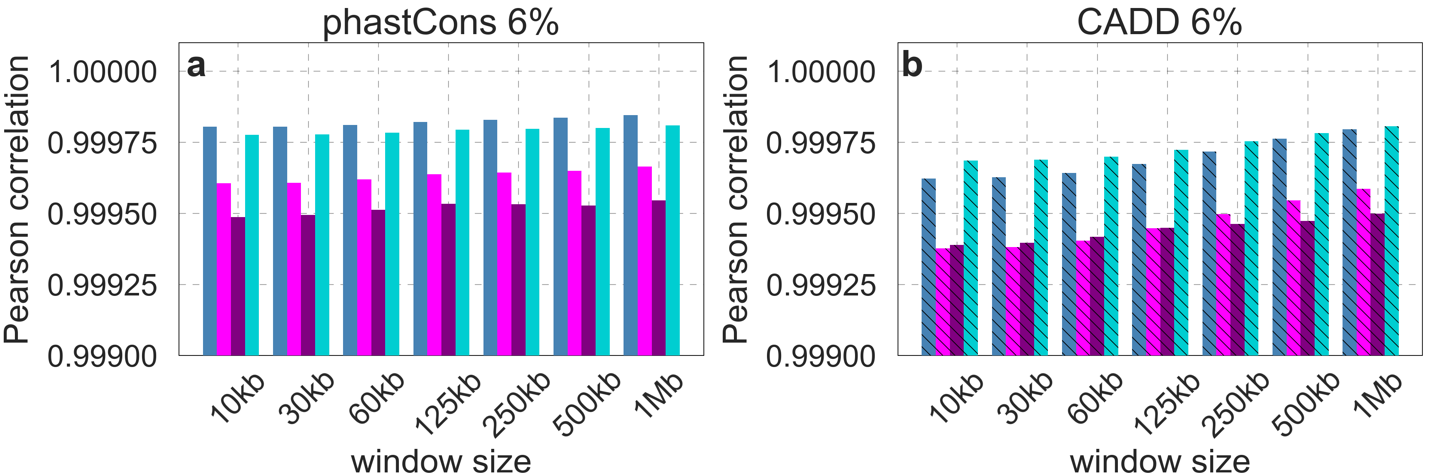

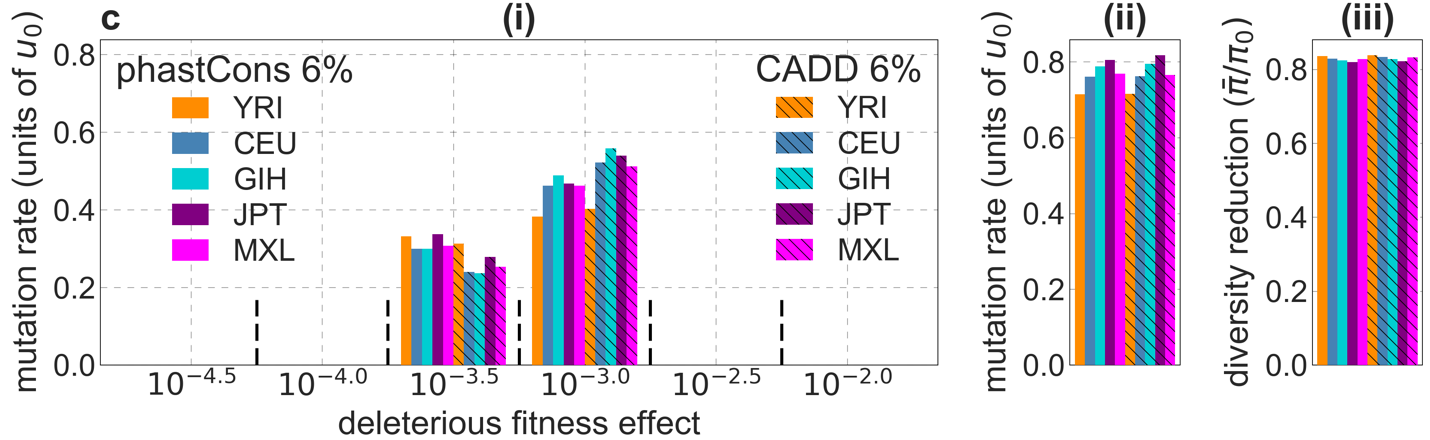

**Figure S35.** The maps and parameter estimates for different populations are remarkably similar. Shown are the results for our best-fitting models using data for one of 1000 Genomes Project populations from each continental group: Africa – Yoruba (YRI), Europe – North-Western European (CEU), South Asia – Gujrati Indian (GIH), East Asia – Japanese (JPT), and Americas – Mexican (MXL). a and b) The Pearson correlations between predictions of relative diversity levels (compared to the population mean) in YRI vs. the other populations for the models based on phastCons (a) and CADD (b) scores. c) Comparison of parameter estimates using data from these populations (panels c (i-iii) as described in panels a (i-iii) in Fig. S14).

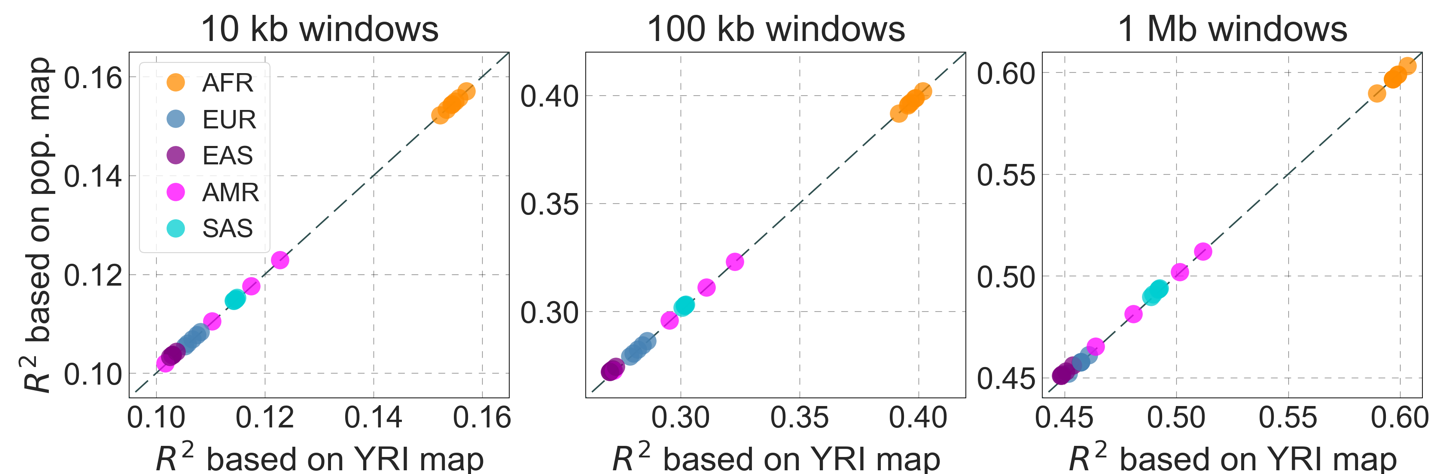

**Figure S36.** The proportion of variance in diversity levels explained ($R^{2}$) in different populations, using the population specific map vs. the YRI map. We show the results for three window sizes (10 kb, 100 kb and 1 Mb) based on our best-fitting CADD-based model. Each point corresponds to one of the 26 populations sampled in the 1000 genomes project and is colored based on continental origin, i.e., African (AFR), European (EUR), South Asian (SAS), East Asian (EAS) and American (AMR).

While the maps inferred in different populations are highly similar, the proportion of variance explained differs substantially among populations (Fig. S36). These differences can be explained by the effects of different demographic histories (e.g., historical changes in effective population sizes) on variation in diversity levels across the genome. To make this more concrete, we consider a simple model for the variance in neutral diversity levels in non-overlapping windows of a given size; for simplicity, we ignore variation in mutation rates across windows. We denote the relative (average) diversity level in window $i$ by $y_{i}={\pi_{i}}/\bar{\pi}$, the predicted relative diversity level in that window by $f_{i}={B_{i}}/\bar{B}$, and the corresponding residual by $e_{i}=y_{i}-f_{i}$, where $\bar{y}=\bar{f}=1$ and $\bar{e}=0$. We can now decompose the variance in relative diversity levels across windows, as

| $V\left( y \right)=V\left( f \right)+V\left( e \right)+2Cov(e,f)$, | $(15)$ |
| --- | --- |

where $V\left( f \right)$ corresponds to the variance due background selection; $V\left( e \right)$, the variance of residuals, can be thought of as reflecting the effects of drift and demographic history; and the covariance, $Cov(e,f)$, can be thought of as reflecting the interaction between background selection and demographic history. Recasting the proportion of variance explained in these terms, we find that

| $R^{2}\equiv1-\frac{V\left( e \right)}{V\left( y \right)}=\frac{V(f)}{V(y)}\left( 1+\beta\right)$, | $(16)$ |
| --- | --- |

where $\beta={2\cdot Cov(e,f)}/{V\left( f \right)}$ is the slope of the linear regression of the residuals against the predictions, which reflects the effects of interactions between background selection and demographic history on diversity levels.

Given that we found the predicted effects of background selection to be highly similar across populations, this modeling exercise sets up a testable prediction: if the interaction terms were nil, the difference in $R^{2}$ among populations should come from the total variance in the denominator, $V\left( y \right)=V\left( f \right)+V(e)$, and specifically from the contribution of demographic history to this variance, $V\left( e \right)$. Fig. S37a-c suggest that while most of the differences in $R^{2}$ among populations are indeed explained by differences in total variance due to demographic history, the interaction terms (the $\beta$s) are non-zero. To examine these interactions further, we look at the relationship between residuals, $e_{i}$, and predictions, $f_{i}$, in several populations (Fig. S37d-h). We find a strong apparent dependency at the low and high ends of our predicted range, presumably reflecting artifacts due to thresholding at the low end (see Section 1.5) and possibly the effects of ancient introgression at the high end (see Section 8); removing 1.5% of the windows at each of these extremes appears to largely remove these effects. In the rest of the range, we find a weak negative correlation between residuals and predictions (which becomes somewhat stronger when we remove the ends). We also find that this correlation varies substantially among populations, e.g., -0.06 to -0.1 for 1 Mb windows, which is what we would expect given differences in demographic history. Thus, our analysis suggests that interactions between demographic history and background selection also contribute to the differences in $R^{2}$ among populations.

In summary, our findings suggest that the effects of background selection are similar across human populations, and that differences among populations in the proportion of variance in diversity levels that our predictions explain are likely due to differences in population demographic history. Interestingly, there appears to be an interaction between the effects of background selection and demography on diversity levels, which varies among populations, as recently suggested by several studies (*62-65*). Our maps of the effects of background selection have enabled us to identify evidence for these interaction effects and should facilitate a better understanding of these effects in the future.

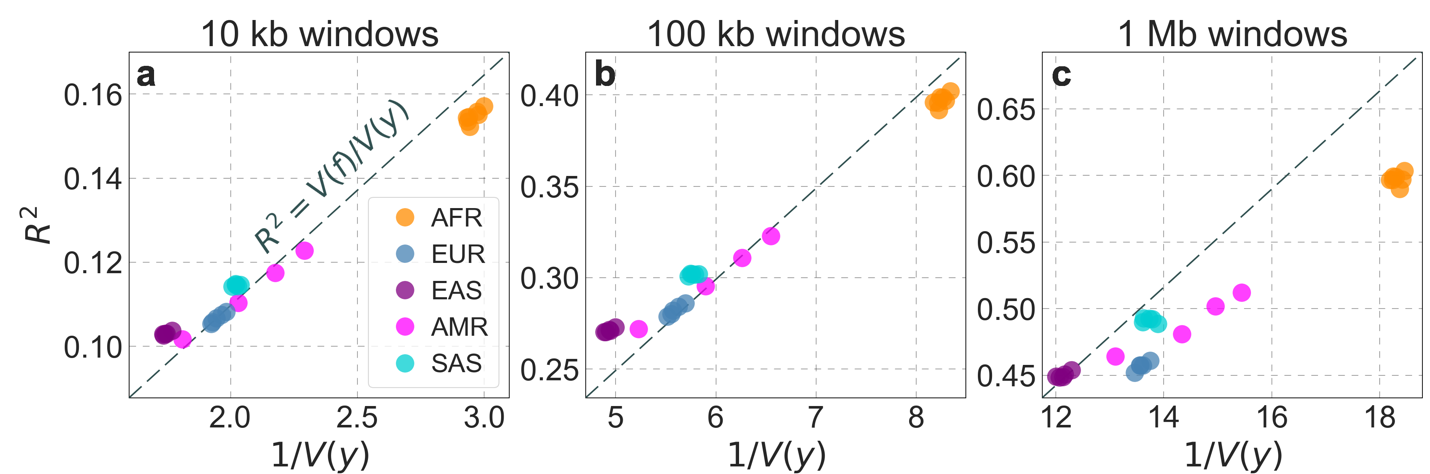

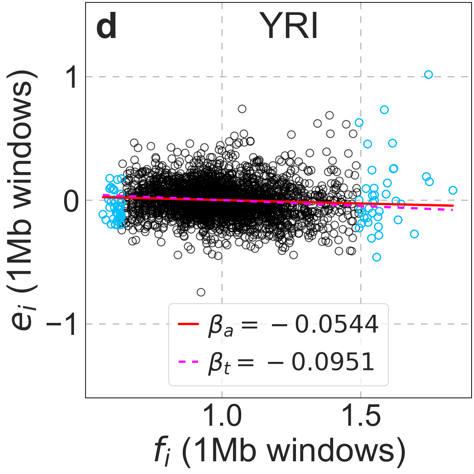

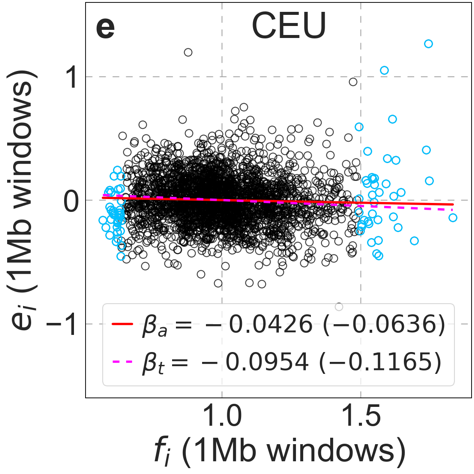

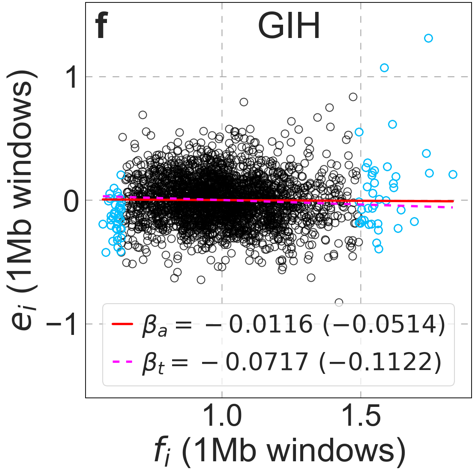

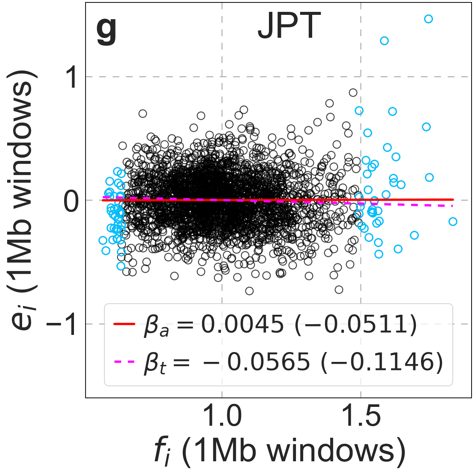

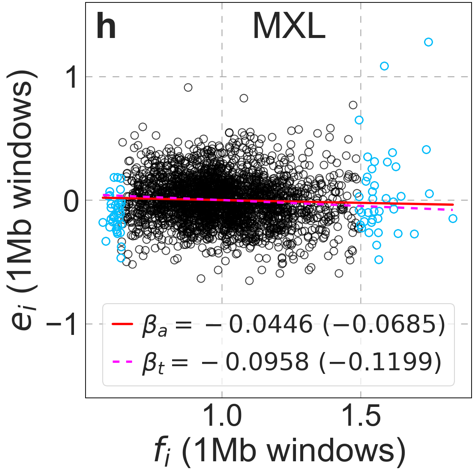

**Figure S37.** Differences among populations in the variance in diversity levels explained by our map of the effects background selection. (a-c) The variance explained ($R^{2}$) as a function of $1/{V\left( y \right)}$ (where $V\left( y \right)$ is the total variance), for three choices of window size. If the interaction terms ($\beta$) were 0, we would expect populations to fall on the dashed line $R^{2}=V(f)\cdot1/{V(y)}$, with slope $V\left( f \right)$ and differences in $V(y)$ due to demographic history. The distances from the dashed line reflect the interaction terms (specifically, $R^{2}-{V\left( e \right)}/{V\left( y \right)}=\beta\cdot{V\left( e \right)}/{V\left( y \right)}$; Eq. 15). The points correspond to the 26 populations sampled in the 1000 genomes project (*19*) and are colored by continental origin as described in Fig. S36. Here we base our predictions on our best-fitting CADD-based map in YRI, but using other population-specific maps yields almost identical results. (d-h) The relationship between the residuals ($e_{i}$) and predictions ($f_{i}$) in representative populations (same as in Fig. S36) on the 1 Mb scale. The 1.5% of windows with lowest and highest predicted values, where our predictions are likely off for various reasons (see Sections 1.5 and 8), are marked in blue. The $\beta$s for each population, with and without extreme points, are shown on the graph (denoted ‘a’ for ‘all’ and ‘t’ for ‘trimmed’, respectively). As above, we use the predictions in YRI, but other population maps yield qualitatively similar results ($\beta$s obtained using the corresponding population specific maps are shown in parenthesis).

### 8. Diversity levels where background selection is weakest ($B\approx1$)

Our maps of background selection effects are well calibrated throughout the range of predicted effects, with two exceptions. One is in the ~4% of sites in which background selection is predicted to be strongest, where predictions are imprecise; this arises from the thresholding approximation we apply in fitting, and is discussed in Section 1.5. The other exception is for sites in which background selection is predicted to be the weakest, where observed diversity levels are markedly greater than expected (Figs. 5 and S38). A close up on this region shows that observed values depart from predictions in the ~2% of sites where $B\gtrsim0.98$ (Fig. S38b). Similar behavior is seen in all 26 populations sampled in the 1000 Genomes Project (Fig. S51). We consider possible explanations for it here.

First, we characterize the main covariates of the strength of background selection (quantified by *B*), such as recombination rate, base composition, and chromosomal position. Beyond the inherent interest in these covariates, they point toward processes that may explain the departure from our predictions near $B=1$. Second, we investigate whether differences in rates of different kinds of mutations and of biased gene conversion associated with the covariates of $B$ can explain the departure near $B=1$; our analysis suggests that they cannot. Third, we argue that the residual effect of ancient introgression between archaic humans and ancestors of extant humans may contribute to this departure.

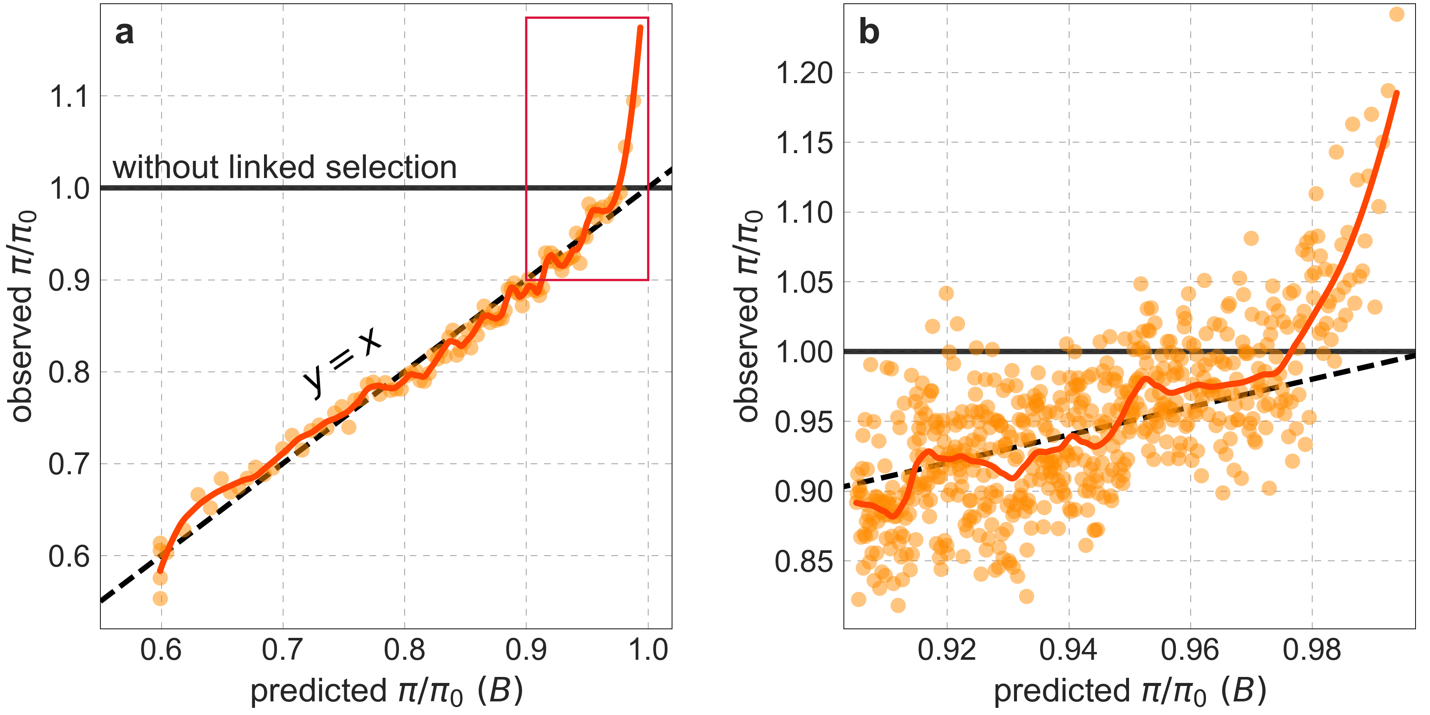

**Figure S38.** Observed vs. predicted neutral diversity levels across the autosomes (similar to Fig. 5). a) Light orange scatter plot: We divide putatively neutral sites into 100 equally sized bins based on the predicted effect of background selection, *B*, from the best-fitting CADD-based model. For predicted values (x-axis), we average the predicted *B* in each bin. For observed values (y-axis), we divide the average diversity level in each bin by the average predicted diversity level in the absence of background selection, $\pi_{0}$, after scaling each in bin by its estimated local (relative) mutation rate ($u(x)/\bar{u}$ in Eq. 8; Section 3.3). Dark orange curve: the LOESS curve for a similarly defined scatter plot but with 2000 rather than 100 bins (with span=0.1). b) A close-up near $B=1$ corresponding to the boxed region in (a). Here, the LOESS curve has span=0.033 and the scatter plot corresponds to 2000 bins (showing the top 30%).

#### 8.1 Covariates of *B*

We expect the effects of background selection to be strongest in regions with low rates of recombination and high densities of functional sites, because neutral variation in such regions will be linked to more deleterious mutations. In line with these expectations, recombination rates increase with greater predicted *B* (Fig. S39a); in particular, they increase sharply between the 99^th^ and 100^th^ percentile of predicted *B* to >10 cM/Mb, which is tenfold the autosomal average. In addition, as expected, the average level of conservation around neutral sites decreases as *B* increases (Fig. S39b and c).

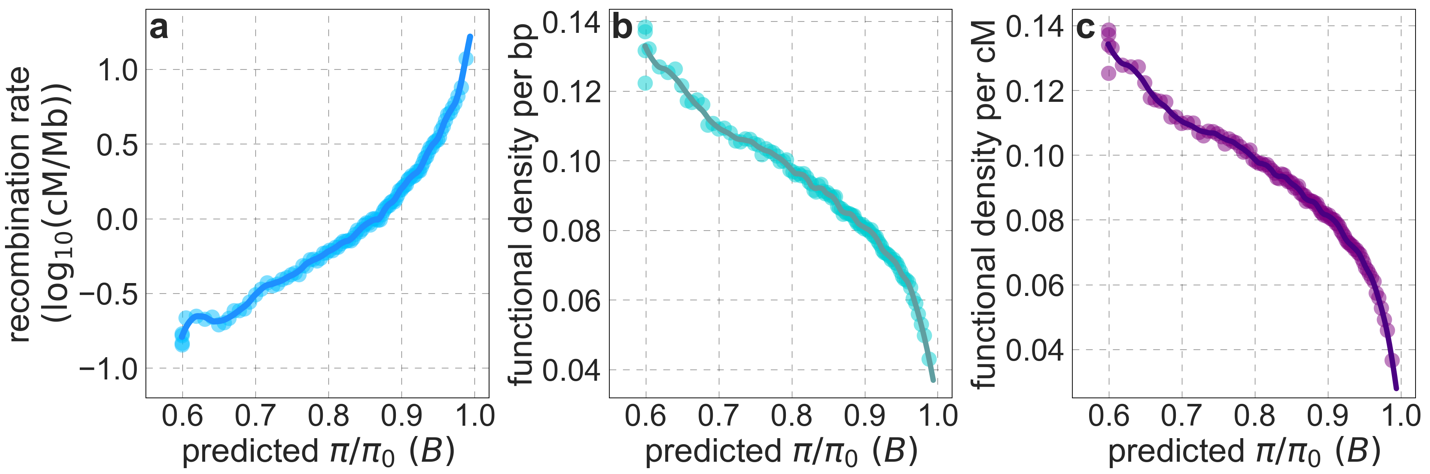

**Figure S39** Average recombination rate (a) and functional density per bp (b) and per cM (c) as a function of predicted *B*. The predicted *B*, binning of putatively neutral sites and LOESS fitting are as described in Fig. S38a. We calculate the average recombination rate in each bin based on the African-American admixture map from Hinch et al. (*24*) (Section 2.3). We measure functional density by calculating the mean phastCons score (based on the 99-vertebrate alignment) in a radius of 50 kb (b) or 0.05 cM (c) around each putatitively neutral site, and averaging these means over sites in each bin.

Next, we consider base composition and other factors that are known to affect rates of mutation and biased gene conversion (BGC). GC content has a J-shaped dependence on predicted *B­* (Fig. S40a). The greater peak in GC content, in regions under weak background selection (*B* near 1), is plausibly largely driven by the long-term effects of BGC due to higher rates of recombination in these regions (*66, 67*) (Fig. S39a). Both peaks (for low and high *B*) are associated with an increase in the proportion of GC sites in CpG islands but this proportion is small throughout (<1%) (Fig. S40b), suggesting that it has little effect on GC content and of mutation rates. In contrast, methylated CpG content increase sharply as predicted *B* approaches 1 (Fig. S40c), suggesting a corresponding increase in the rate of C>T transitions. The proportion of sites in C>G hypermutable regions also increases with predicted *B* (Fig. S40d).

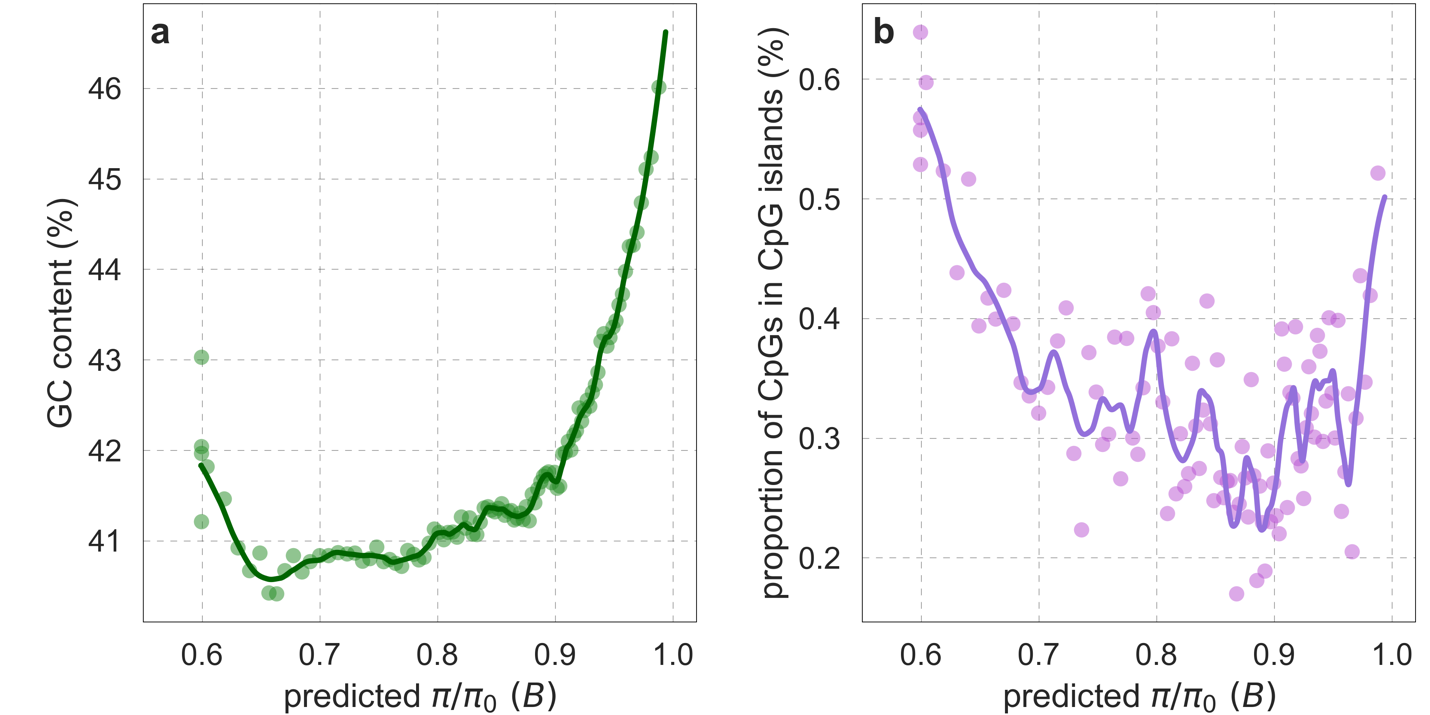

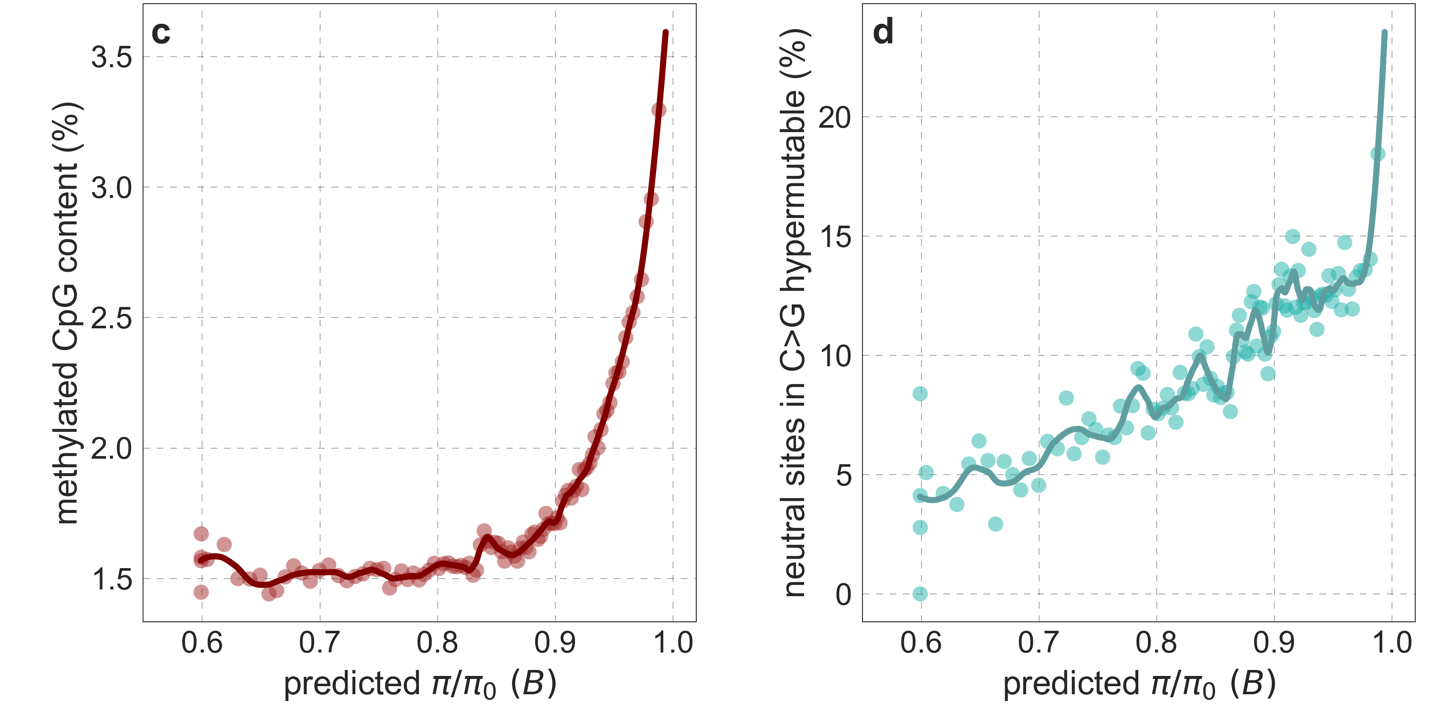

**Figure S40.** GC content (a), CpG sites in CpG islands (b), proportion of methylated CpGs in neutral sites (c), and proportion of neutral sites in C>G hypermutable regions (d) as a function of predicted *B*. Proportions and other quantities are measured for putatively neutral sites, whose type (i.e., GC and CpG) is defined based on the inferred state in the human-chimpanzee ancestor (Section 2.7). Data sources are detailed in Section 2.8. The predicted *B*, binning of putatively neutral sites and LOESS fitting are as described for Fig. S38a.

Lastly, predicted *B* is associated with chromosomal position, with regions under weak background selection (*B* near 1) clustered near telomeres (Fig. S41). In turn, regions near telomeres are early replicating, and replication timing is known to affect mutational patterns (*68*).

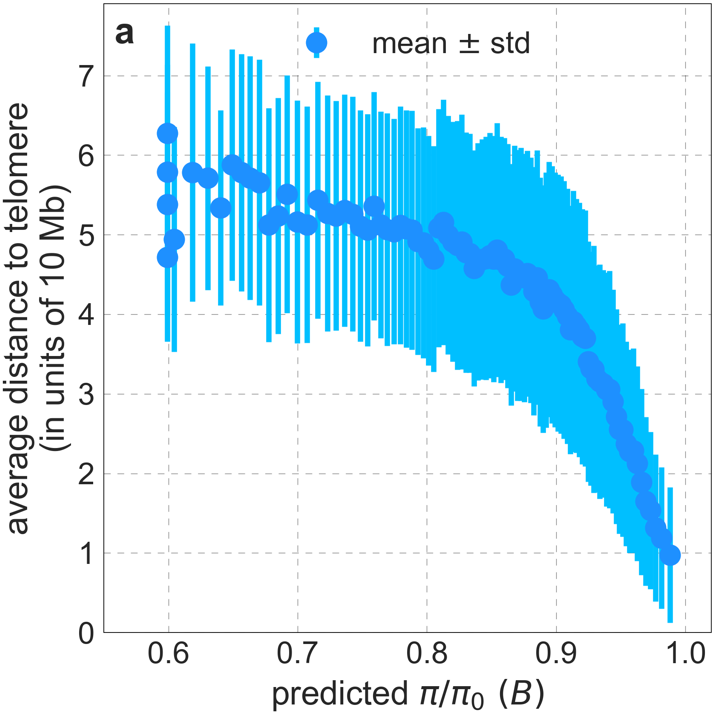

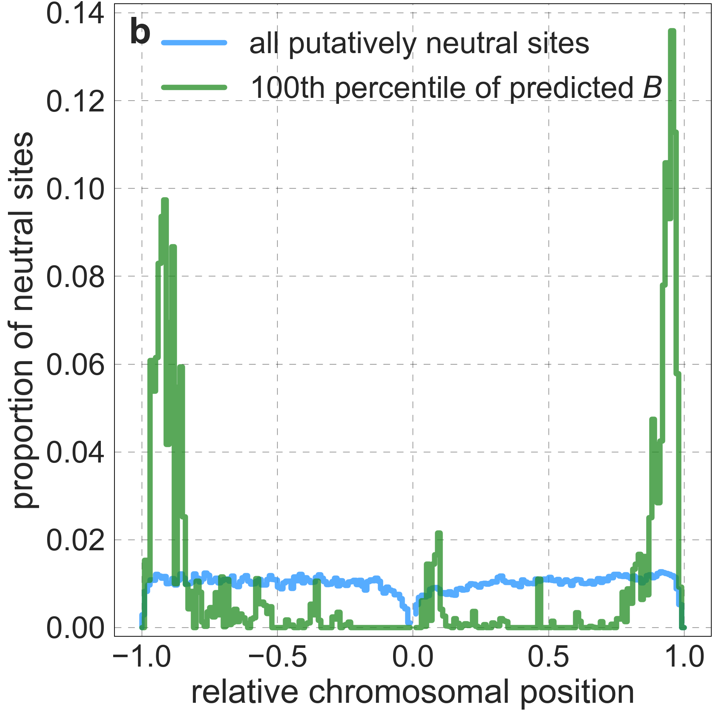

**Figure S41.** The relationship between chromosomal position and predicted *B.* a) The the distance to telomeres is measured on the same chromosome (see Section 2.8). b) The distribution of putatively neutral sites by relative chromosomal position, defined as the ratio of a site’s distance to the centromere and the distance between centromere and telomeres on that chromosomal arm (see Section 2.8); relative distances corresponding to the shorter chromosome arm are shown on the left (in [-1, 0]) and to the longer arm on the right (in [0, 1]). The binning of putatively neutral sites by predicted *B* is as described for Fig. S38a.

#### 8.2 Mutational spectrum and biased gene conversion

As we noted, the covariates of *B* are associated with mutational processes and with biased gene conversion that affect diversity and divergence levels (Fig. S42). Specifically, we see the footprints of the following processes:

- Increased rates of **A>C/T>G** and **A>G/T>C** substitutions near $B=1$ (Fig. S42a), due to higher rates of biased gene conversion that are associated with the higher rates of recombination (*66, 67*) (Fig. S39a). Biased gene conversion also reduces the rates of **C>A/G>T** and **C>T/G>A** substitutions, but this is not clearly visible (Fig. S42b), presumably because of other processes affecting these substitutions (see below).
- Increased rate of **C>T mutations** near $B=1$ (Fig. S42b), associated with the higher methylated CpG content (*33*) (Fig. S40c).
- Reduced rates of **C>A/G>T** and **A>T/T>A** mutations near $B=1$ (Fig. S42 a and b), associated with improved repair of these types of mutations near origins of replication, which tend to be near telomeres (*68*) (Fig. S41).
- Increased rates of **C>G/G>C** mutations near $B=1$ (Fig. S42b), due to the enrichment of C>G hypermutable regions (*34*) (Fig. S40d).

Consequently, the rates of substitutions between any two bases covary with predicted *B* (Fig. S42). However, the different types of substitutions have markedly different contributions to levels of diversity and divergence (Fig. S43), and different dependencies on predicted *B* (Fig. S42).

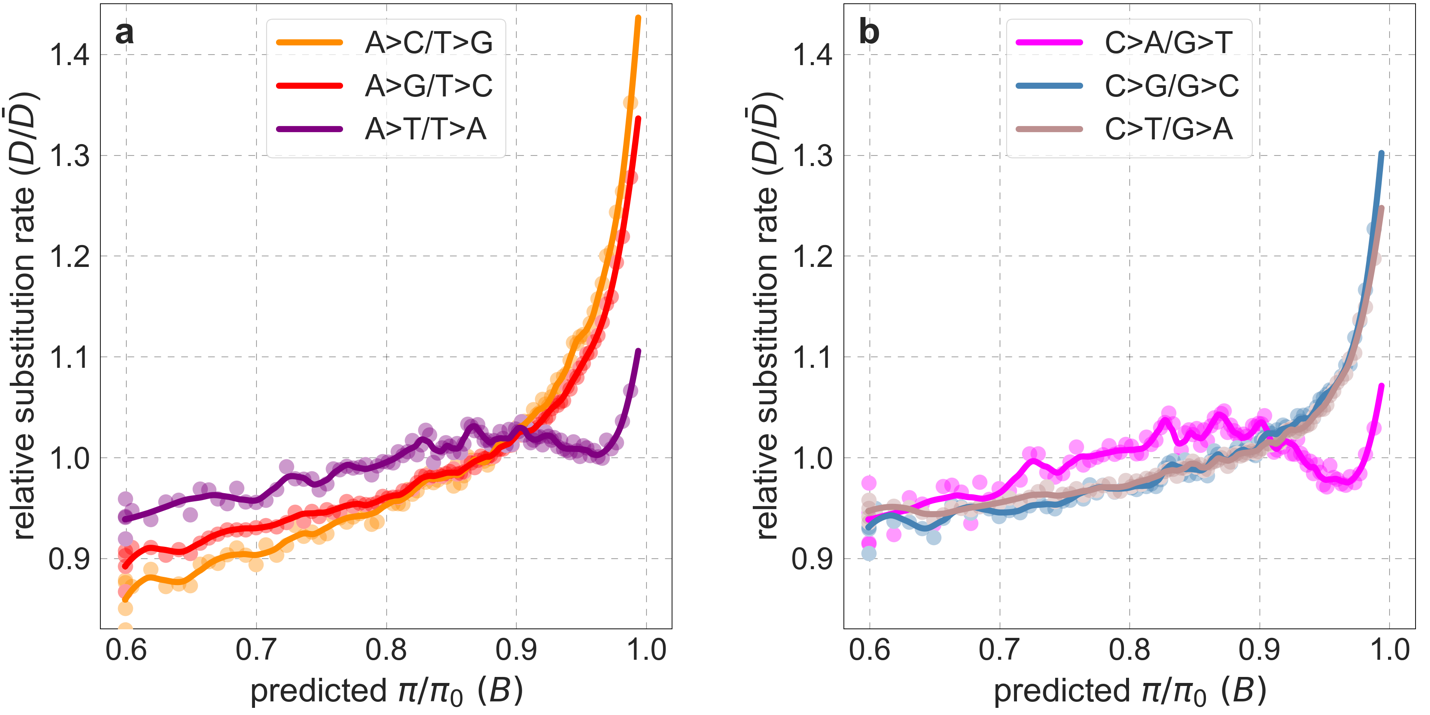

**Figure S42.** Rates of different types of substitutions as a function of predicted *B*. We bin putatively neutral sites by predicted *B* as described in Fig. S38a. We calculate the rate of X>Y substitutions in a bin by dividing the estimate of the number of X>Y substitutions at its sites in an 8-primate phylogeny by the estimated number of its sites with state X in the ancestor of that phylogeny (see Section 3.3). We obtain the relative rate by dividing the rate in a bin by the average rate across bins. We show the rates of substitutions with ancestral state AT in (a) and GC in (b).

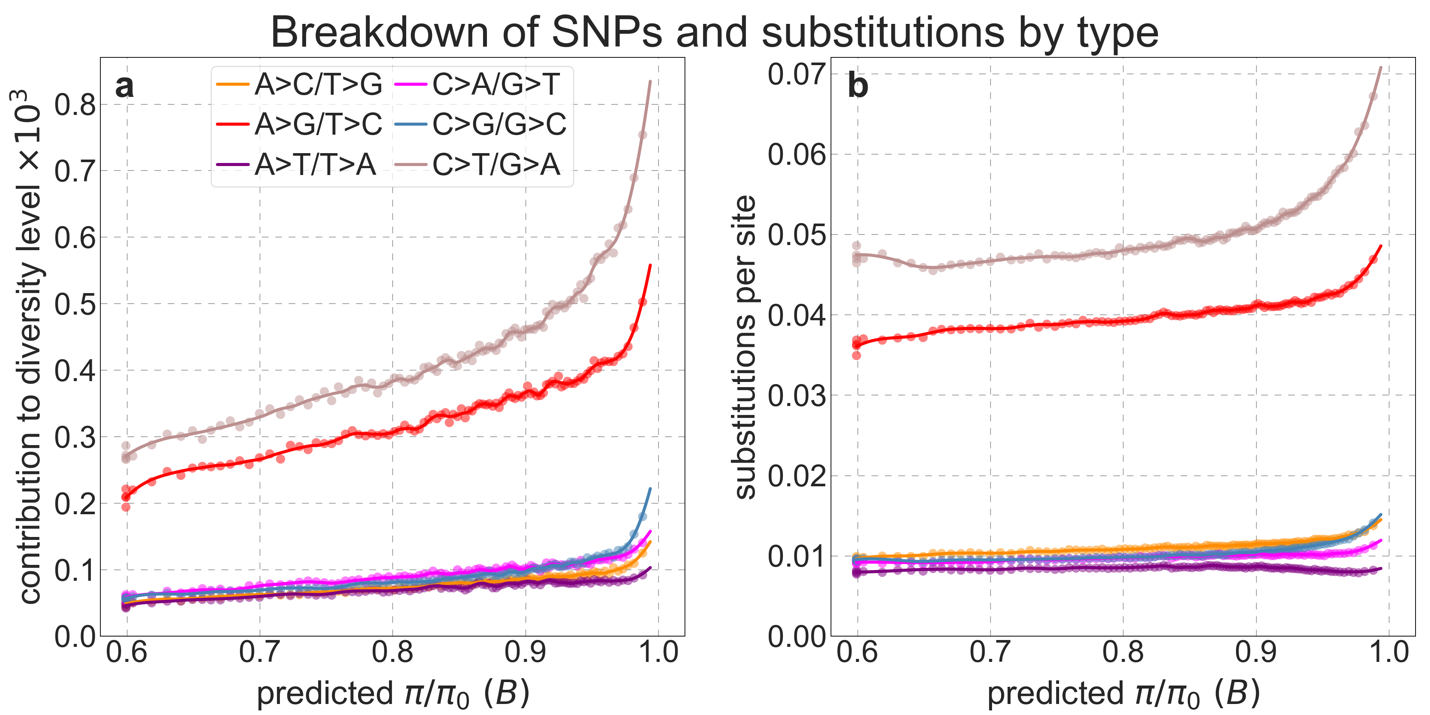

**Figure S43.** Contribution of different types of substitutions to diversity levels (a) and number of substitutions per site in the 8-primate phylogeny (b) as a function of predicted *B*. We bin putatively neutral sites by predicted *B* as described in Fig. S38a. a) We define the ancestral state, i.e., AT or GC, as the inferred state in the human-chimpanzee ancestor (see Section 2.7). We define the contribution of each type of substitution to the diversity level in a bin as the ratio of the number of pairwise differences of that type and the total number of pairwise comparisons in a bin; this way, the sum over types equals the observed diversity level in a bin ($\hat{\pi})$. b) We calculate the number of substitutions of each type in a bin as described in Fig. S42, but here we normalize it by the number of sites, such that the sum over types equals the observed number of substitutions per site in the 8-primate phylogeny.

To investigate whether these processes can explain the departure from our predictions near $B=1$, we break up the observed diversity levels by types of substitutions (Fig. S44). We reason that if all types behave similarly near $B=1$, the differential processes affecting them cannot explain the departure from predictions (at least not fully). Note that, up to a multiplicative constant, our observations are ratios of diversity levels and substitution rates (on the 8-primate phylogeny), implying that the signatures of processes that affect diversity and divergence similarly should cancel out. Conversely, for a process to affect our observations it must have noticeably different effects on diversity and divergence. We find that the observations associated with different types of substitutions largely align with each other and with the observations that include all types jointly (Figs. S38 and S44). Specifically, they align with predictions throughout nearly the entire range of predicted *B*, and are markedly higher than predictions near$B=1$.

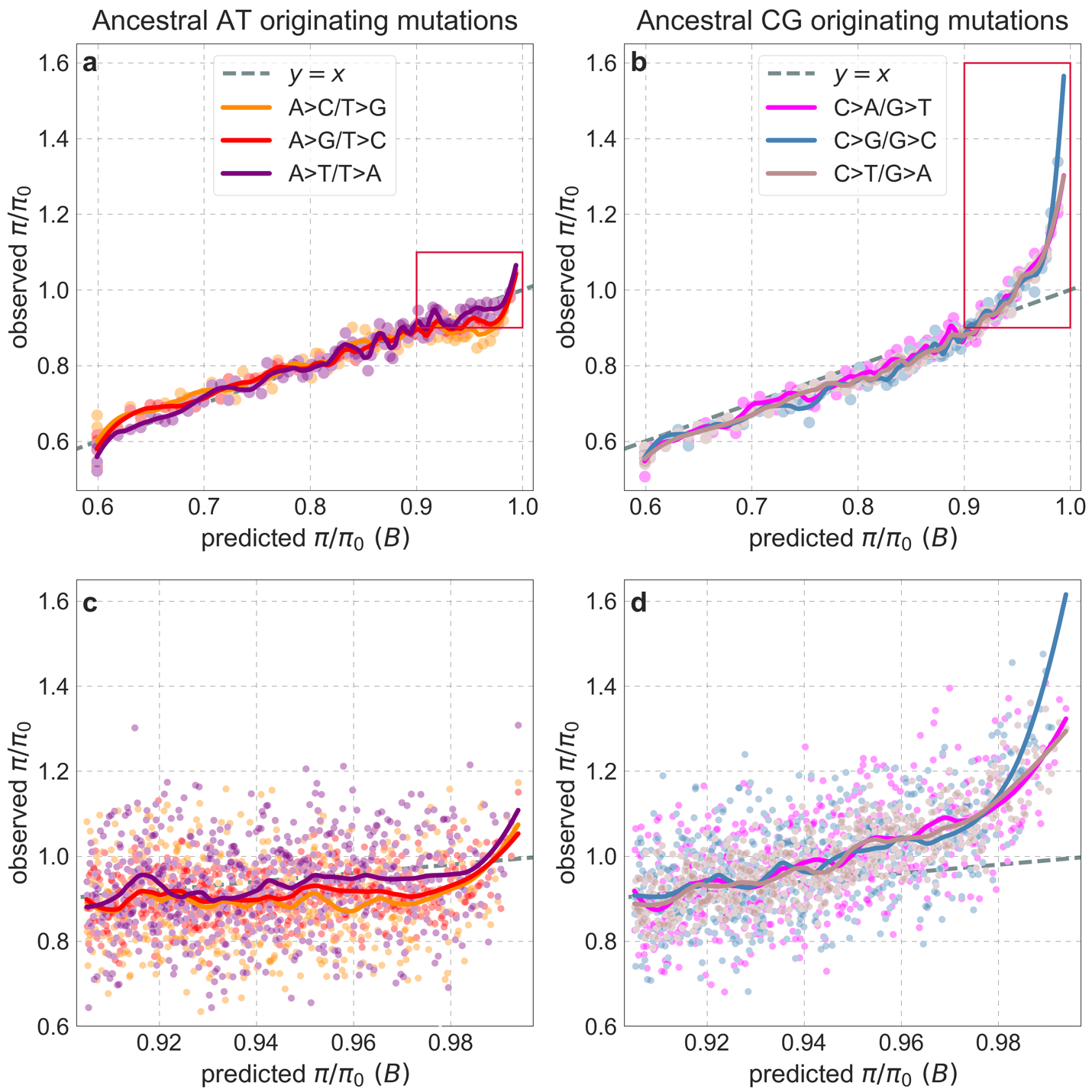

**Figure S44.** Observed vs. predicted neutral diversity levels for different types of substitutions. Diversity levels are presented as in Fig. S38, but here, we calculate diversity levels and substitution rates for sites with a given ancestral state, i.e., AT (a and c) or CG (b and d), and for each of the three types of substitutions from the ancestral state, as in Fig. S43.

Nonetheless, the observed levels associated with C>G mutations near $B=1$ are markedly higher than for other types of substitutions (Fig. S44b and d). This effect contributes negligibly to the total observed levels near$B=1$, however, because C>G substitutions have a minor contribution to total diversity and divergence levels (Fig. S43). The higher levels of C>G substitutions are associated with the enrichment of C>G hypermutable regions near $B=1$ (Fig. S40d). Notably, when we remove these regions, the observed levels associated with C>G substitutions are no longer higher than for other types of substitutions (Fig. S45). Removing C>G hypermutable regions also affects the magnitude of the departure from predictions for other types of substitutions (compare Fig. S45c and d with Fig. S44c and d), because C>G is not the only type of mutation whose rate is higher in these regions. These hypermutable regions plausibly affect diversity more than divergence (and thus our observations) given that their effects are stronger in recent human evolution than in the more distant past and in the lineages of closely related species (Ipsita Agarwal and Molly Przeworski, personal communication). This may reflect the fact that these regions were identified in extant humans and/or a dependence of their effects on life history (*34, 69*). Setting the causes aside, even when we remove these regions from the set of putatively neutral sites used in our inference, observed levels of all types are still markedly higher than the revised predictions near $B=1$ (Fig. S46).

In summary, while we cannot rule out that there are other mutational processes that contribute to the departure from predictions near $B=1$, our analysis suggests that known mutational processes and biased gene conversion fall short of explaining these departures.

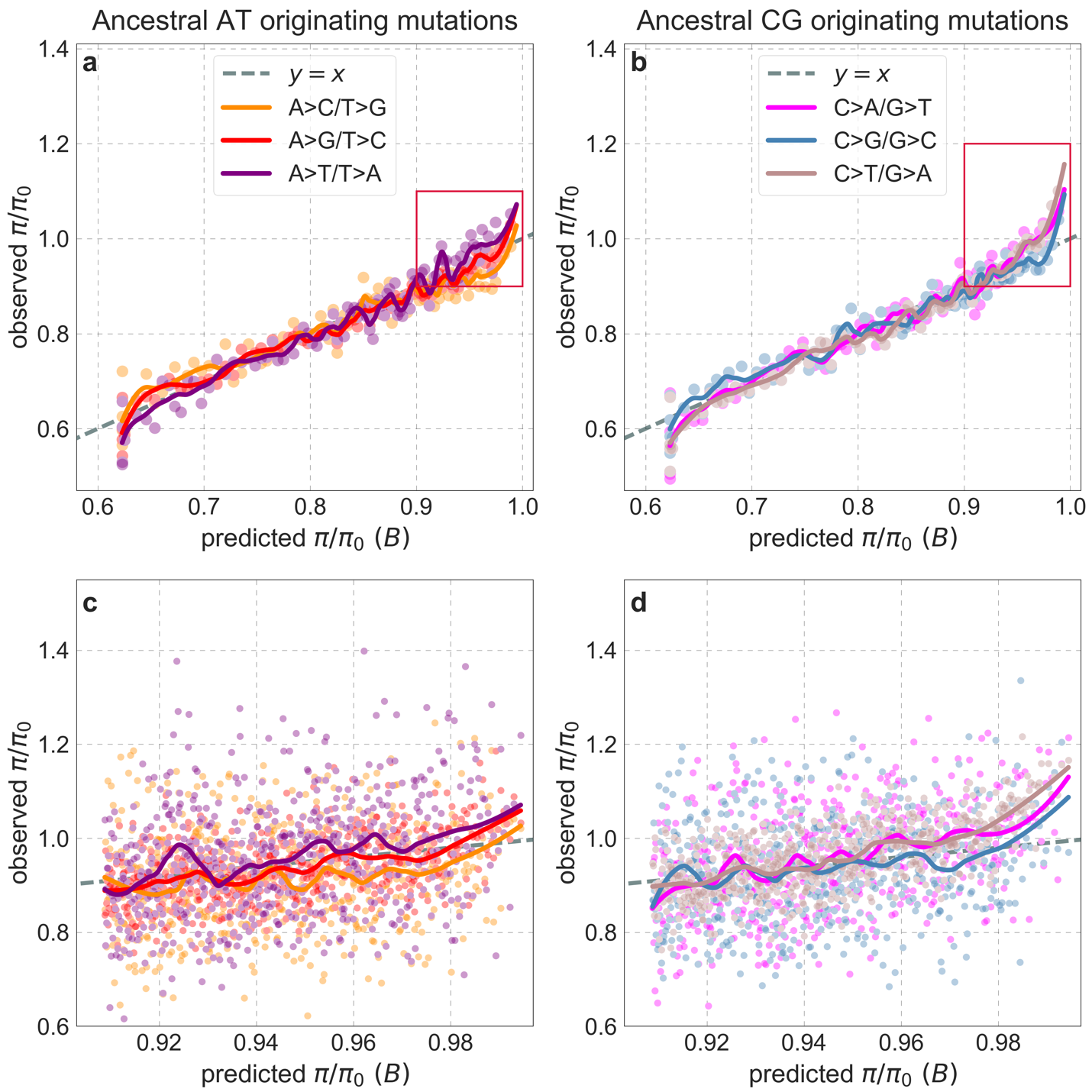

**Figure S45.** Observed vs. predicted neutral diversity levels for different types of substitutions after removing C>G hypermutable regions (from both the inference and observations). Other than removing ~12% of putatively neutral sites in these regions, the details are as in Fig. S44. We note that while a greater proportion of sites is removed from bins near $B=1$ (~20% for the 100^th^ percentile), this in itself has a minor effect on the departures from predictions in these bins.

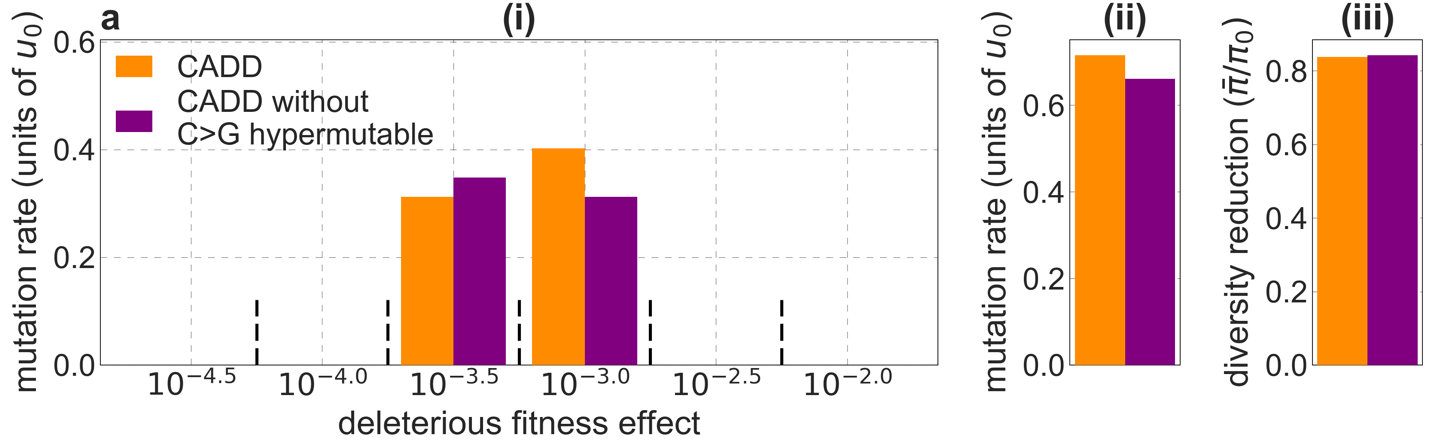

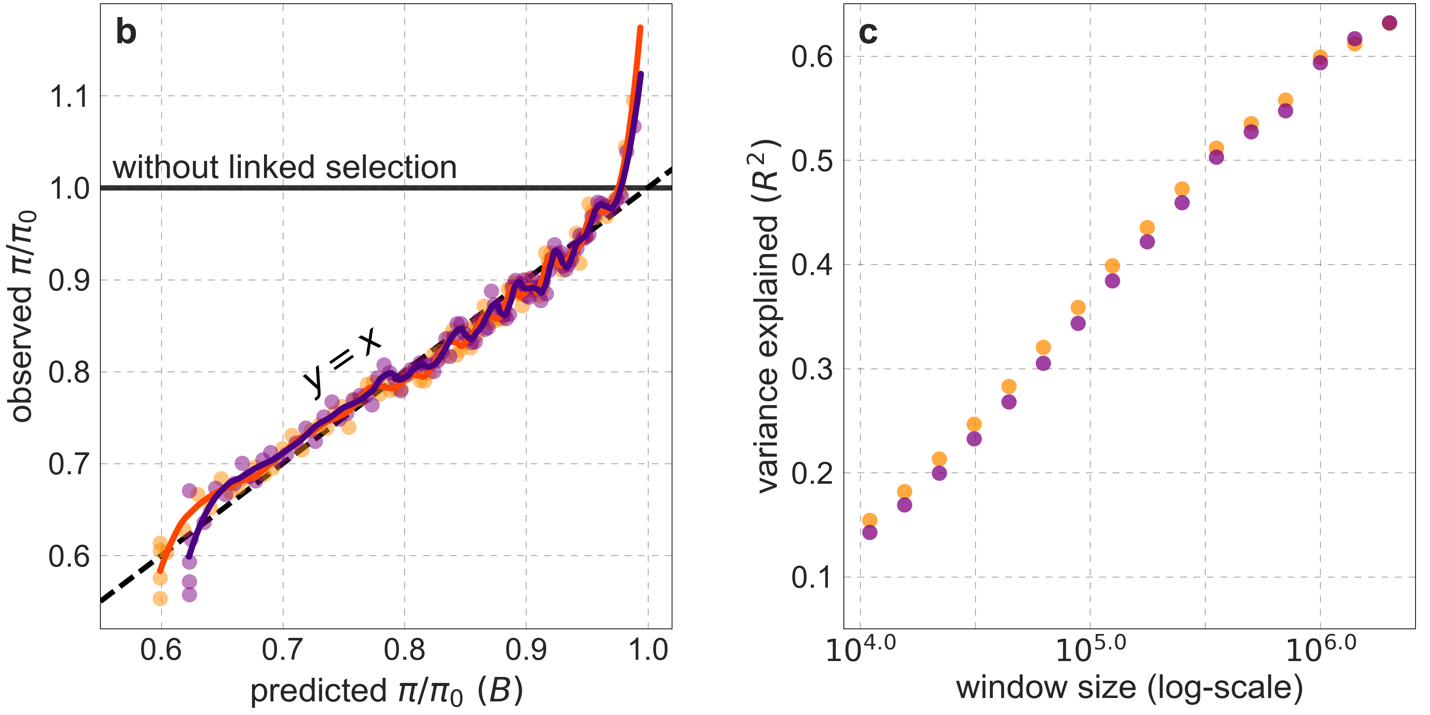

**Figure S46.** Comparison of the best-fitting CADD-based models with and without C>G hypermutable regions. All panels are as described for the corresponding ones in Fig. S14.

#### 8.3 A footprint of archaic introgression?

Next, we consider whether the excess diversity observed near $B=1$ could reflect a residual signal of archaic introgression. The presence of archaic alleles at a locus increases diversity because their coalescence with modern human alleles traces back to the ancestors of modern humans and the archaic hominin from which they originated. Archaic introgression could help to explain the excess diversity in regions with $B\approx1$, if archaic alleles were more common in these regions. As we argue below, there are good reasons to believe this to be the case.

Aside from evidence for positive selection on introgressed alleles in a few cases (*70-72*), the pattern of Neanderthal and Denisovan introgression in contemporary human populations appears to be dominated by purifying selection to remove archaic ancestry from the human genome, as evidenced by the depletion of archaic introgression in and around genes (*70, 71, 73-75*). The causes for this purifying selection are still being deliberated (*73, 75, 76*). One hypothesis is that selection acts against introgressed alleles that are incompatible with the genetic background in modern humans, e.g., alleles that are part of Dobzhansky-Muller incompatibilities between archaic hominins and modern humans (*70, 76*). Another hypothesis is that selection acts against alleles that were deleterious in both archaic hominins and modern humans, which were more common in archaic hominins because their long-term effective population sizes were smaller than in modern humans (*35, 73, 75*).

Regardless of its cause, we expect purifying selection to remove archaic alleles, including neutral variants, more rapidly in genomic regions under stronger background selection (*73, 76, 77*). This is because these regions harbor more selected sites (Fig. S39b) in which archaic alleles could be selected against, and because they have lower rates of recombination (Fig. S39a) causing selection against archaic alleles to remove larger archaic segments. Conversely, we expect the highest, residual proportion of archaic neutral variants in regions with $B\approx1$—precisely where we observe a 10-15% excess of diversity above our predictions (Fig. S38).

In order to test this expectation, we use fine-scale maps of archaic introgression inferred for European (CEU) and East-Asian (CHB/CHS) individuals from the 1000 Genomes Project (*35*). These maps assign a probability of Neanderthal ancestry to contiguous 500 bp segments tiling individual genomes based on the high-coverage Altai Neanderthal genome (*78*). We use them to estimate the average proportion of archaic ancestry per putatively neutral site in bins of predicted *B*. As expected, we find that the estimated proportion of archaic alleles increases with predicted *B* (Fig. S47). The power to identify introgressed segments using this and other methods decreases substantially in regions with $B\approx1$, because higher recombination rate in these regions results in much shorter archaic segments (*35, 79*). We therefore expect that the actual proportion of archaic ancestry increases more sharply near $B=1$ than our analysis suggests, and may therefore better trace the sharp increase in diversity relative to predictions near $B=1$.

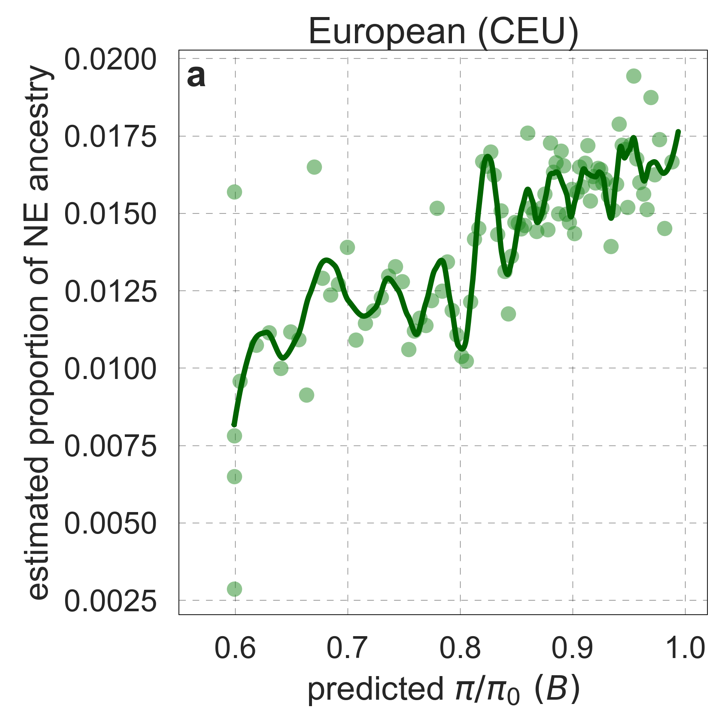

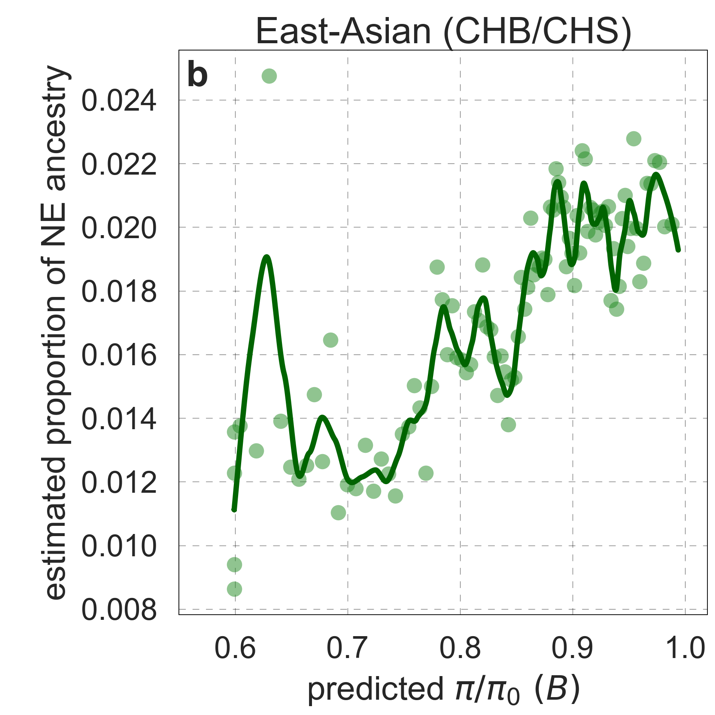

**Figure S47.** Estimated proportion of Neanderthal (NE) ancestry as a function of predicted *B* in Europeans (CEU) (a) and East-Asians (CHB/CHS) (b). See text for the estimation procedure. The bins and LOESS curves were calculated as in Fig. S38.

Current inferences about archaic introgression are divided into those that incorporate sequenced Neanderthal and Denisovan genomes (*35, 70, 71, 80, 81*), such as the maps we used in Fig. S47, and those that are based only on patterns of variation in contemporary humans (*79, 82-84*). When we repeat the analysis in Fig. S47 using ancestry-maps based on the latter approach in both Africans and non-Africans (*79, 83*), we find that levels of archaic ancestry either increase and level off at intermediate values of *B*, or peak at intermediate values and decrease as *B* approaches 1. We believe that this departure from our expectation reflects a decrease in the power of these methods near $B=1$ (due to higher rates of recombination), which is greater than the decrease for methods based on sequenced archaic genomes. An additional caveat is that the evidence for the contribution of archaic introgression to the African gene pool is based solely on patterns in contemporary genetic variation (*82-85*) and remain more speculative in lieu of more direct evidence. Thus, while it seems plausible that the greater retention of neutral archaic variants in regions with the highest ~2% of predicted *B* values contributes substantially to the departure from our predictions in both African and non-African populations, at present, the evidence for such a contribution remains equivocal.

### 9. Additional Figures

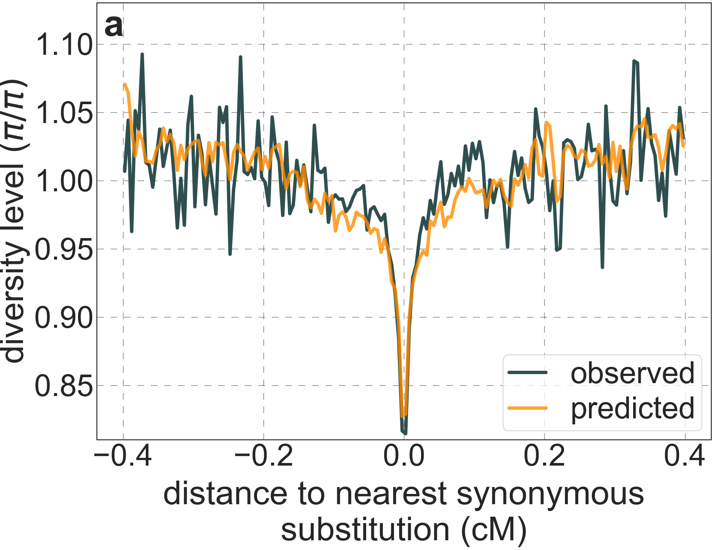

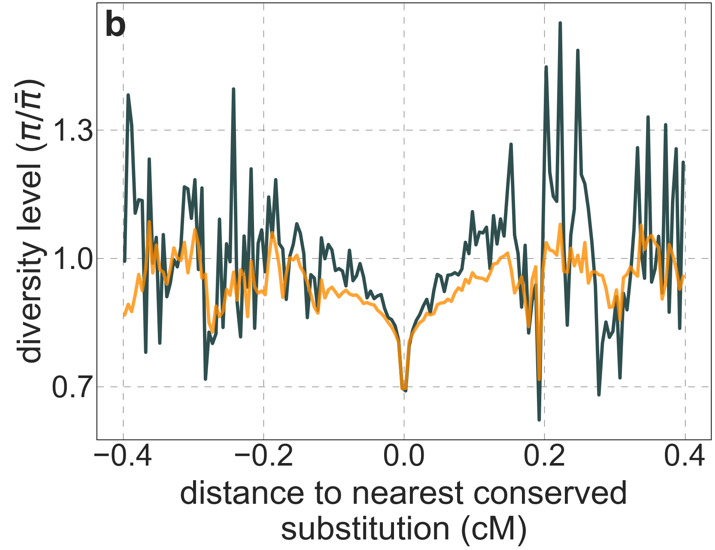

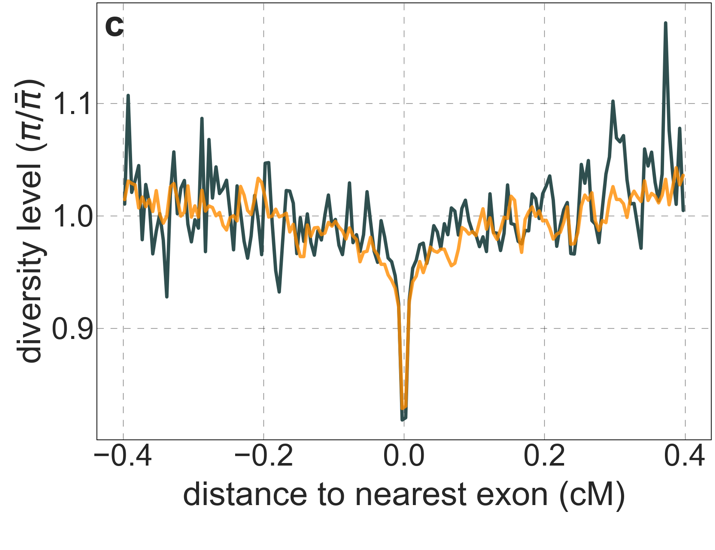

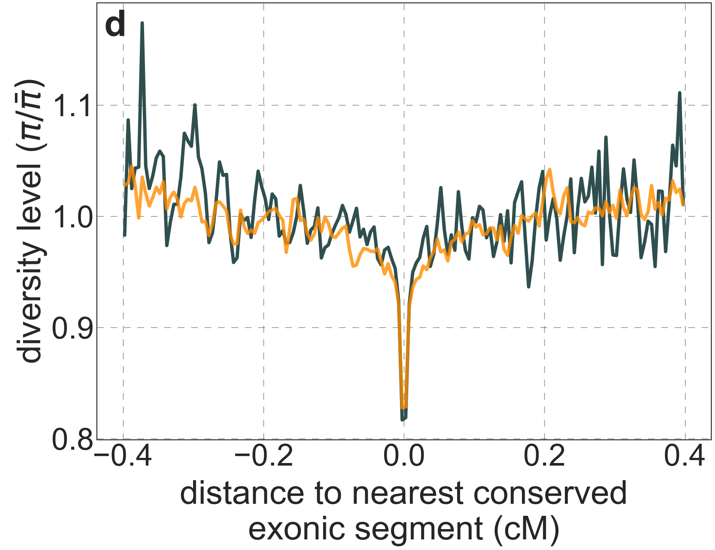

**Figure S48.** A background selection model predicts neutral diversity levels around different genomic features. Here we use our best-fitting CADD-based model and show diversity levels around: a) human-specific synonymous substitutions; b) human-specific substitutions in conserved regions; c) exons; and d) conserved exonic regions. The inference of human-specific substitutions is described in Section 2.7. Conserved regions are based on autosomal sites with the top 6% phastCons scores in the 99-vertebrate alignment (Section 4.1). The set of exons is described in Section 2.4. The genetic distance to the nearest element (e.g., exon) is measured to its closest edge. Other details are similar to Fig. 3.

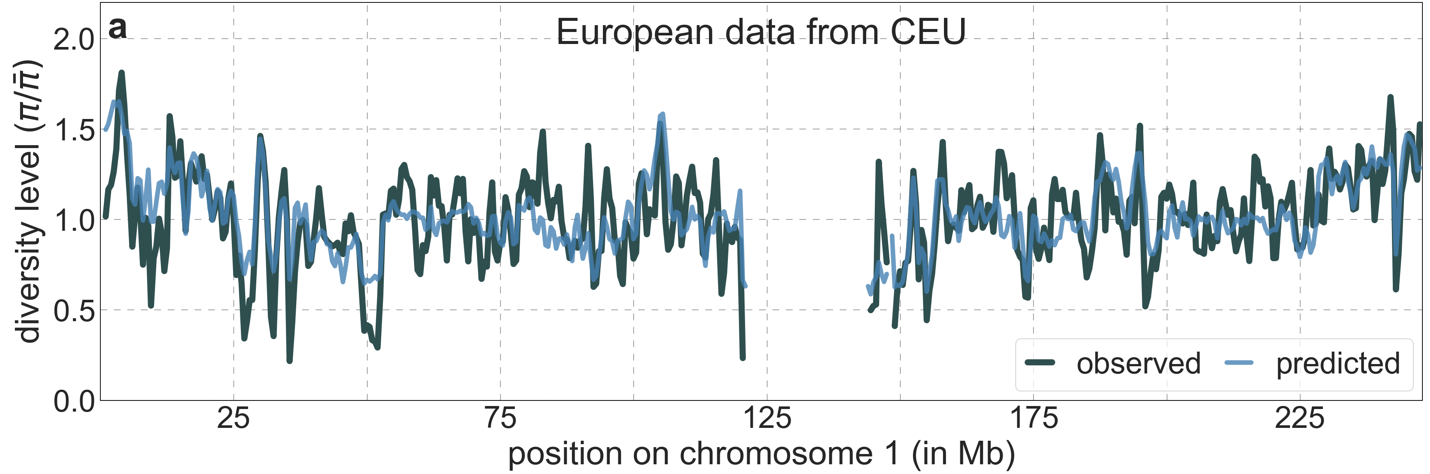

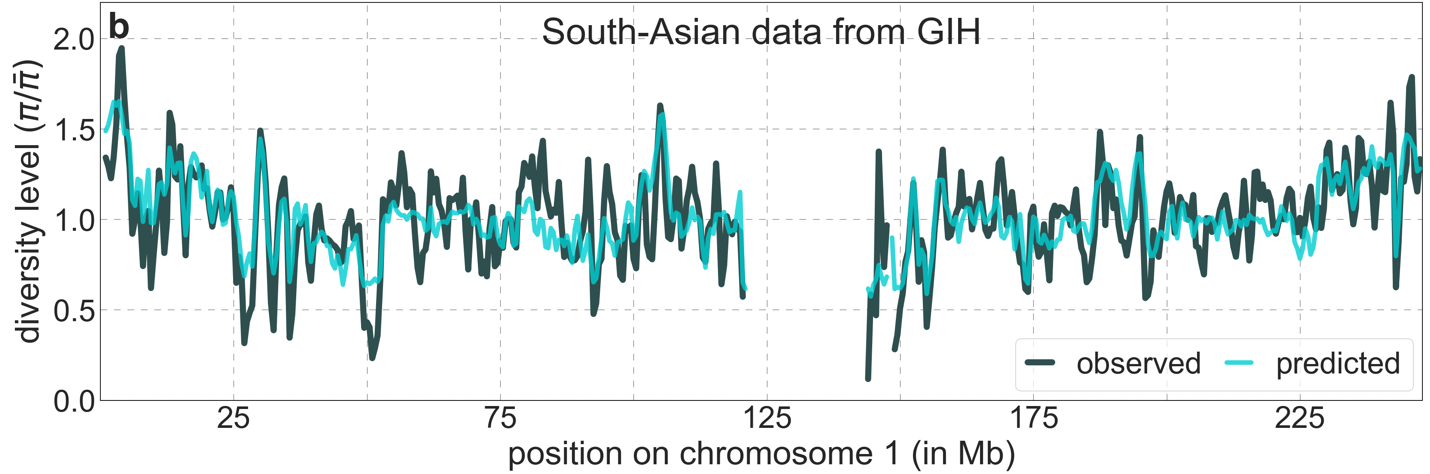

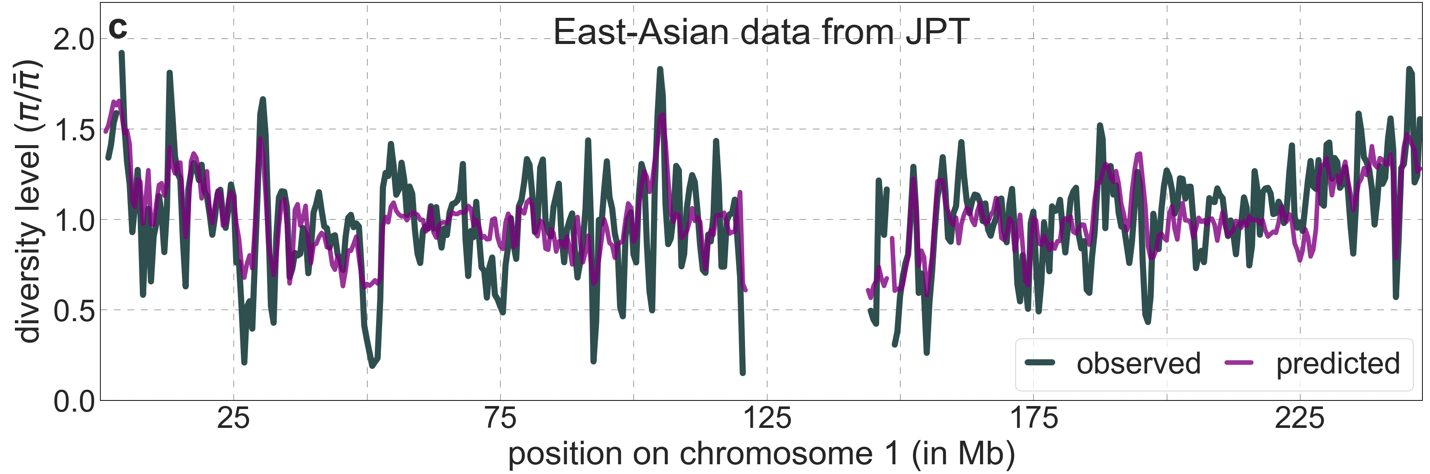

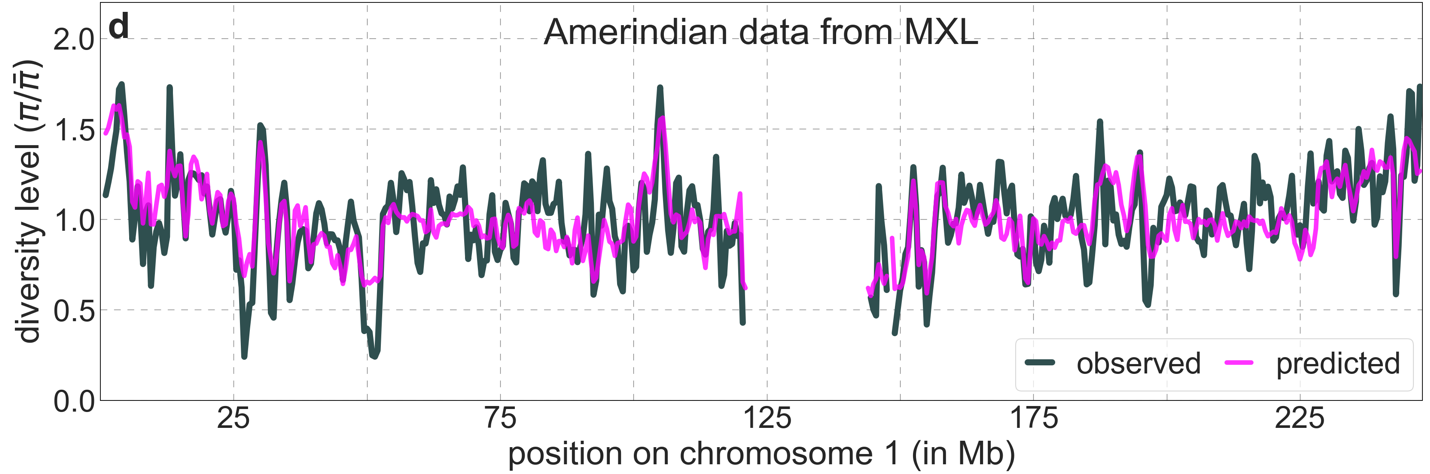
 **Figure S49.** Predicted and observed neutral diversity levels along chromosome 1 based on data from representative continental populations. Plots are generated as detailed in Fig. 2A.

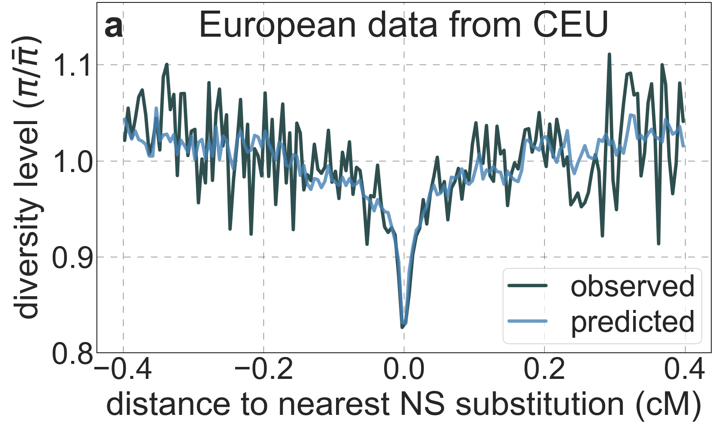

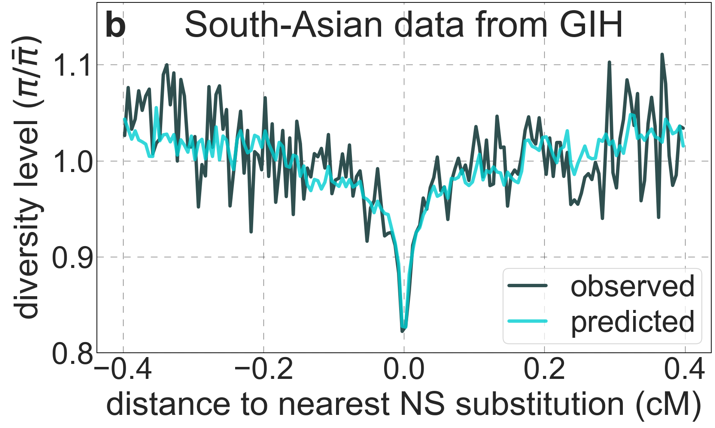

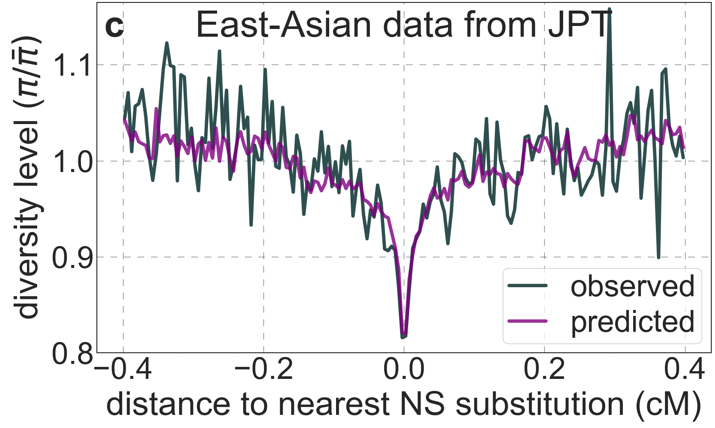

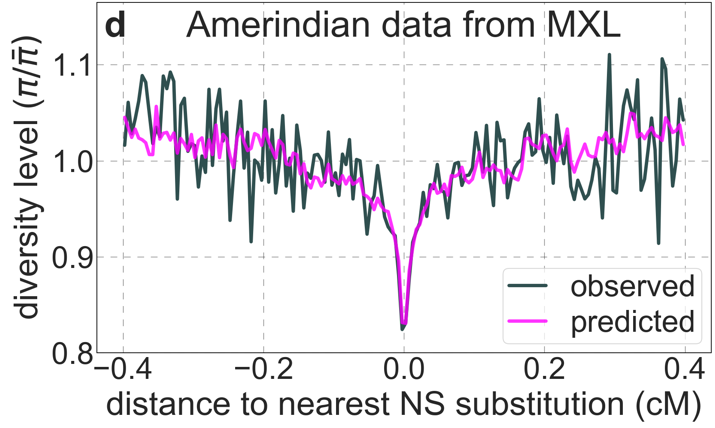

**Figure S50.** A background selection model predicts neutral diversity levels around human-specific nonsynonymous (NS) substitutions in representative continental populations. Plots are constructed as detailed in Fig. 3, using polymorphism data from each population for both inferences and observations.

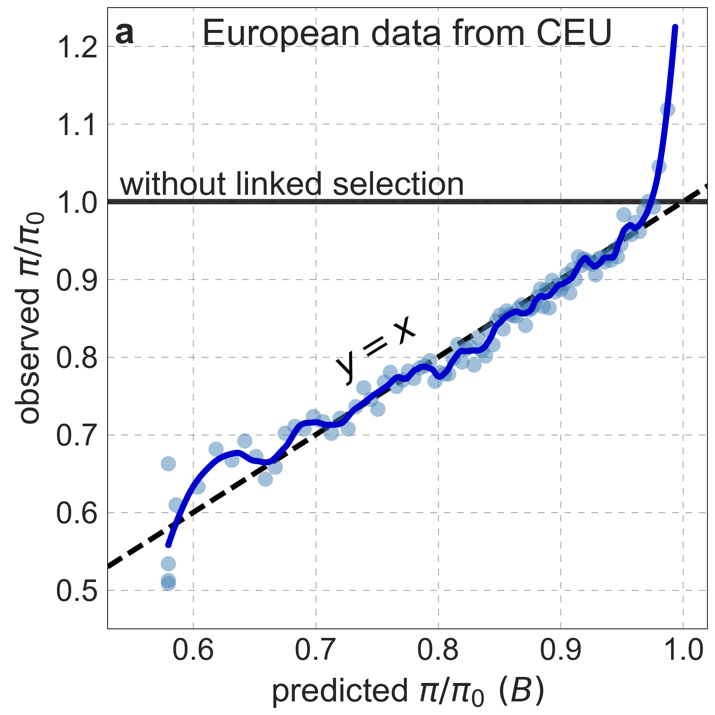

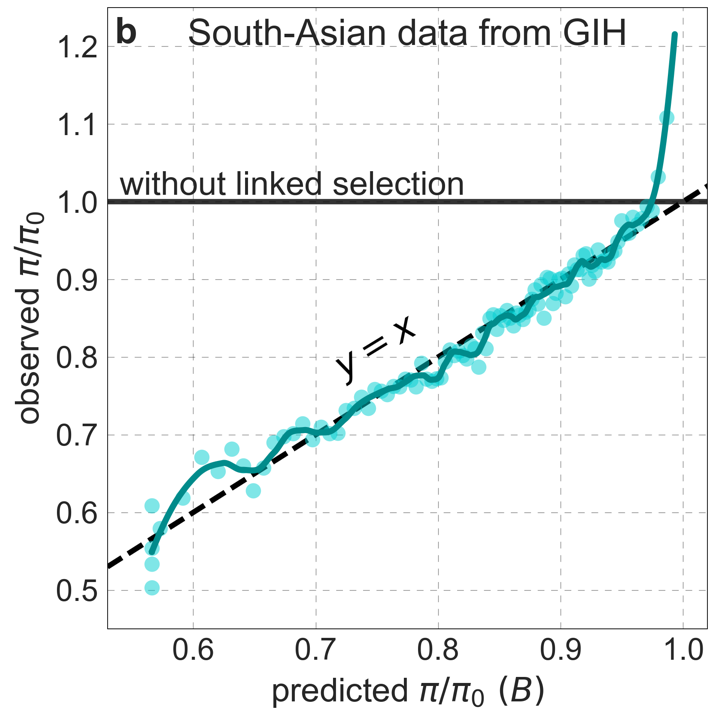

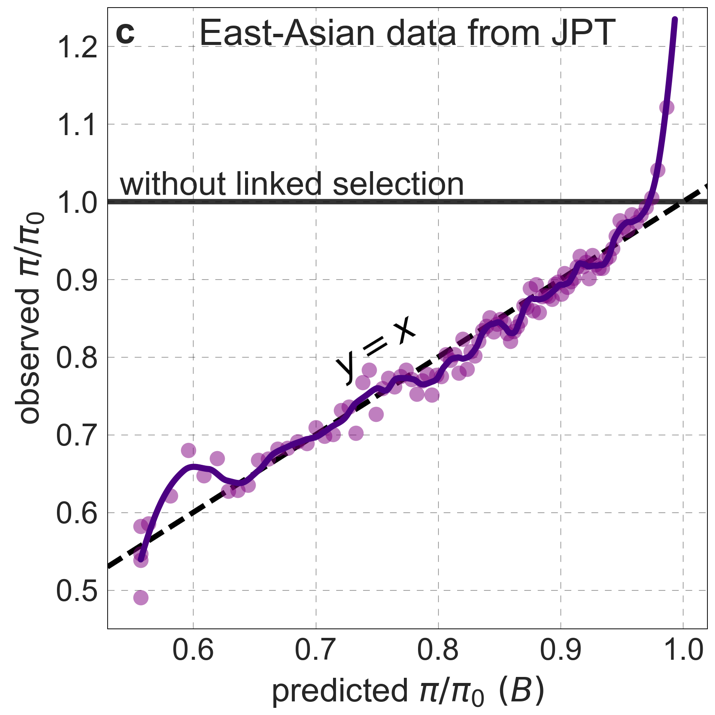

**Figure S51.** Observed vs. predicted neutral diversity levels across the autosomes in representative continental populations. Plots are constructed as detailed in Fig. 5, using polymorphism data from each population for both inferences and observations.
